# (3*R*, 7*S*)-11-hydroxy-jasmonic acid is a major oxidative shunt product of jasmonate catabolism in *Arabidopsis thaliana*

**DOI:** 10.1101/2025.10.03.680387

**Authors:** Kotaro Matsumoto, Maria Mitsui, Takuya Kaji, Riqui Gao, Naoki Kitaoka, Taketomo Otaki, Yuho Nishizato, Hideyuki Matsuura, Minoru Ueda

## Abstract

Jasmonoyl-L-isoleucine (JA-Ile) is a pivotal oxylipin plant hormone that regulates numerous physiological processes and stress responses. Because JA-Ile is synthesized from jasmonic acid (JA), the cellular level of JA is tightly controlled. Although the oxidative catabolism of JA to 12-hydroxy-JA (12-OH-JA) is well-characterized, the biosynthesis, accumulation dynamics, and biological functions of 11-hydroxy-JA (11-OH-JA) remain poorly understood. Here, we conducted a comprehensive investigation of 11-OH-JA in *Arabidopsis thaliana*, encompassing its chemical synthesis, accumulation patterns, biological activity, and biosynthetic pathways.

Using chemically synthesized 11-OH-JA of naturally occurring (3*R*,7*S*)-stereochemistry, we identified 11-OH-JA as the predominant hydroxylated JA derivative that accumulates following wounding. Enzymatic assays and *in silico* docking studies revealed that JOX1/2/3/4 exclusively catalyze the conversion of JA to 11-OH-JA, whereas 12-OH-JA is produced by distinct pathways. Notably, the quadruple mutant *joxQ* completely abolished 11-OH-JA accumulation without affecting 12-OH-JA levels. Furthermore, 11-OH-JA exhibited no binding affinity for the COI1-JAZ co-receptor, indicating that it acts as a biologically inactive shunt product of JA catabolism.

Our findings establish JOX-mediated 11-hydroxylation of JA as a primary inactivation pathway, separate from 12-hydroxylation. Parallel, non-redundant inactivation routes deliver layered signal termination of costly defense signals. This study provides new insights into JA catabolism, highlighting 11-OH-JA as a crucial factor in the attenuation of jasmonate signaling, and thereby redefining the framework of JA turnover in *A. thaliana*.

## INTRODUCTION

The oxylipin hormone (3*R*,7*S*)-jasmonoyl-L-isoleucine (JA-Ile, **Figure 1A**) is synthesized from C18/C16 polyunsaturated fatty acids in response to wounding, pathogen infection, or insect herbivory ^1-5^. JA-Ile governs a wide array of physiological processes, collectively referred to as jasmonate responses, including growth, fertility, senescence, cold acclimation, and specialized metabolites production etc. JA-Ile exerts these effects by initiating profound changes in the cellular transcriptome through a unique co-receptor complex comprising the F-box protein CORONATINE INSENSITIVE 1 (COI1) and the JASMONATE ZIM DOMAIN (JAZ) transcriptional repressor ^6,7^. Because constitutive JA-Ile signaling can compromise plant fitness ^8^, signal termination must be as robust as activation. Across organisms, termination rarely relies on a single switch; rather, layered architectures prevent the costs of prolonged signaling. Jasmonate responses are especially sensitive to this economy of activation and shut-off.

**Figure 1.**
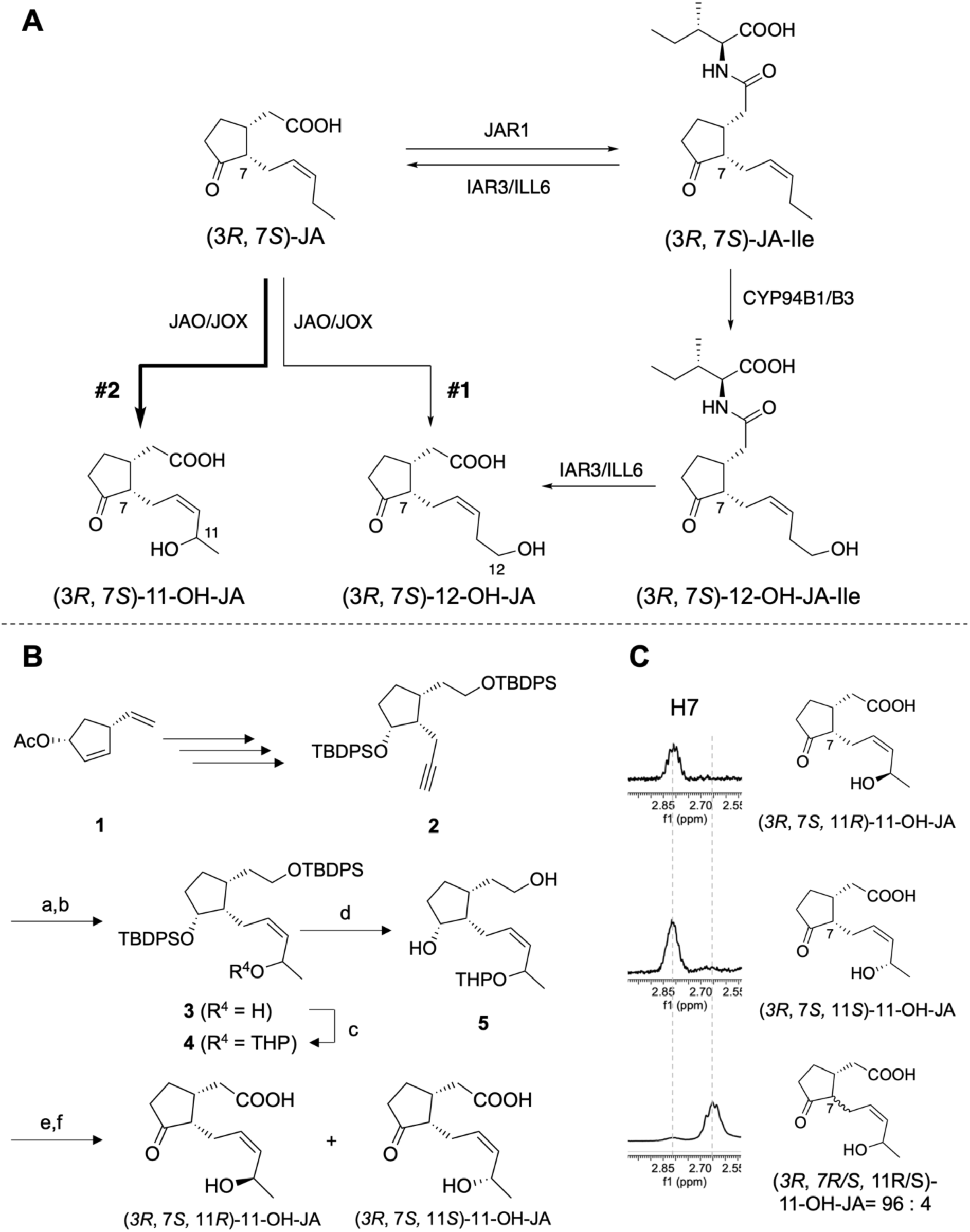
Current understanding of catabolic pathway of (3*R*, 7*S*)-JA: (**A**) Current (**#1**) and updated (**#2**) understanding of catabolic pathway of (3*R*, 7*S*)-JA : (3*R*, 7*S*)-JA is converted to (3*R*, 7*S*)-11-OH-JA and (3*R*, 7*S*)-12-OH-JA through distinct pathways. (3*R*, 7*S*)-12-OH-JA and (3*R*, 7*S*)-12-OH-JA-Ile are further converted respectively into corresponding carboxylic acids, (3*R*, 7*S*)-12-COOH-JA and (3*R*, 7*S*)-12-COOH-JA-Ile (not shown). (**B**) Outline of chemical synthesis of (3*R*, 7*S*, 11*R*/*S*)-11-OH-JA. The synthetic (3*R*, 7*S*, 11*R*/*S*)-11-OH-JA was obtained as a mixture of 11*R*/*S*-stereoisomers. (a) *n*BuLi, acetaldehyde, THF, -78 °C, 69%; (b) H_2_, Pd/BaSO_4_, quinoline, MeOH, 89%; (c) DHP, *p*-TsOH•H_2_O, CH_2_Cl_2_, 98%; (d) TBAF, THF 91%; (e) Jones reagent, acetone, -20 °C, 96%; (f) MgBr_2_, Et_2_O 38%. (**C**) The stereochemistry of 11-OH group of an isolated stereoisomer of **3** was determined as *11S* by modified Mosher method based on the chemical shift difference between (*S*)-MTPA ester and (*R*)-MTPA ester (8^S^–8^R^); (**D**) Comparison of expanded view of ^1^H-NMR of the synthesized (3*R*, 7*S*, 11*R*)-11-OH-JA (top), (3*R*, 7*S*, 11*S*)-11-OH-JA (middle), and a separately synthesized mixture of (3*R*, 7*S*, 11*R*/11*S*)-11-OH-JA and (3*R*, 7*R*, 11*R*/11*S*)-11-OH-JA (4:96) around δ 3.0-2.5 (H7 region).

In *Arabidopsis thaliana*, the principal mechanism for JA-Ile attenuation is catabolic inactivation ^3,9,10^. JA-Ile is sequentially oxidized by cytochrome P450 monooxygenases CYP94B1, CYP94B3, and CYP94C1 to (3*R*,7*S*)-12-hydroxy-jasmonoyl-L-isoleucine (12-OH-JA-Ile, **Figure 1A**) and then to (3*R*,7*S*)-12-carboxy-jasmonoyl-L-isoleucine (12-COOH-JA-Ile, not shown) ^11-16^. Whereas 12-OH-JA-Ile retains weak bioactivity ^17-19^, 12-COOH-JA-Ile is inactive. Additionally, JA-Ile and 12-OH-JA-Ile are hydrolyzed by the amidohydrolases ILL6/IAR3 to form (3*R*,7*S*)-jasmonic acid (JA) and (3*R*,7*S*)-12-hydroxy-JA (12-OH-JA), respectively (**Fig. 1A)** ^20^. Parallel to JA-Ile turnover, free JA is also oxidized, historically attributed to 2-oxoglutarate-dependent dioxygenases JASMONIC ACID ^21,22^ (**Figure 1A, #1**). Because 12-OH-JA does not assemble the COI1–JAZ complex ^18^, 12-OH-JA is viewed as a shunt that helps quench signaling upstream of JA-Ile oxidation. Notably, another oxidized product, (3*R*,7*S*)-11-hydroxy-JA (11-OH-JA, **Figure 1A**), has been detected in multiple plant species ^23-26^, and accumulates in *A. thaliana* leaves following wounding, at levels even higher than 12-OH-JA ^26^. Analytical conflation of 11- and 12-hydroxy isomers of JA has further obscured product assignment.

Here we resolve this ambiguity and reposition jasmonate turnover within a layered termination logic. By using stereochemically defined 11-OH-JA standards, we reveal a **parallel** inactivation route in which JOX dioxygenases oxidize JA selectively at C11 to generate biologically inactive 11-OH-JA in upstream of JA-Ile oxidation. This framework reframes jasmonate catabolism overview from a single terminal outlet to parallel outlets of layered architectures, which provide fast, fail-safe shut-off of costly defense signaling.

## RESULTS

### Chemical Syntheses of (3*R*,7*S*, 11*R*)-11-OH-JA and (3*R*,7*S*, 11*S*)-11-OH-JA

To elucidate the biological role of 11-OH-JA, we first planned to synthesize a stereochemically pure reference standard. Previous synthetic efforts yielded a mixture of (3*R*,7*R*)- and (3*R*,7*S*)- isomers at the C7 position, predominantly furnishing thermodynamically stable (3*R*,7*R*)-11-OH-JA ^24^. Here, we optimized the synthetic route by introducing ketone functional groups in the cyclopentane ring at later stages of the synthesis, thereby minimizing isomerization to the thermodynamically more stable (3*R*,7*R*)-form (**Figure 1B**).

Following the approach of Nonaka *et al.* for *cis*-12-oxo-phytodienoic acid (*cis*-OPDA) synthesis ^27^, we started the synthesis with the hydroboration of **1** (**Figure 1B**). After a series of transformations (**Figures 1B** and **S1**), we obtained the key intermediate **3**, which contained the characteristic (*Z*)-alkene side chain of 11-OH-JA. Nuclear Overhauser effect (NOE) correlations between H11 and H8/H8’ in synthetic intermediate **3** confirmed the *cis*-geometry of olefine moiety (**Figure S2A)**. Subsequent oxidation and deprotection of **4** successfully provided 11-OH-JA, from which two stereoisomers, (3*R*,7*S*, 11*R*)-11-OH-JA and (3*R*,7*S*, 11*S*)-11-OH-JA, were isolated by reverse-phase HPLC (**Figures 1B** and **S1**). NOE correlation between H3 and H7 verified the natural (3*R*, 7*S*)-stereochemistry (**Figures S2B**). Comparison of the ^1^H-NMR spectra of the isolated stereoisomers, (3*R*,7*S*, 11*R*)-11-OH-JA and (3*R*,7*S*, 11*S*)-11-OH-JA, with those of the a separately synthesized mixture of (3*R*, 7*S*, 11*R*/11*S*)-11-OH-JA and (3*R*, 7*R*, 11*R*/11*S*)-11-OH-JA (4:96) ^24^ indicated the absence of (3*R*, 7*R, 11R/S*)-11-OH-JA in our synthetic samples (**Figures 1C and S3**). In addition, the C11 stereochemistry was determined by using intermediate **3**: two diastereomers of **3** was separated and each stereochemistry was determined by using a modified Mosher’s method (**Figure S4A**) ^28^. The (3*R*,7*S*, 11*S*)-**3** was converted to (3*R*,7*S*, 11*S*)-11-OH-JA (**Figure S4B**) and the retention time was compared with those of a mixture of (3*R*,7*S*, 11*R*/*S*)-11-OH-JA to determine the stereochemistry (**Figure S4C**). These results allowed us to unambiguously assign the structures of the two synthetic standards: (3*R*, 7*S*, 11*R*)-11-OH-JA and (3*R*, 7*S*, 11*S*)-11-OH- (**Figure S4C**).

This refined synthetic strategy provided sufficient quantities of highly pure (3*R*,7*S*, 11*R*)-11-OH-JA and (3*R*,7*S*, 11*S*)-11-OH-JA, enabling accurate analytical separation from its isomeric counterpart, 12-OH-JA, and facilitating functional assays that had been previously hampered by lack of a suitable standard.

### Time-Course Accumulation of 11-OH-JA After Wounding in *Arabidopsis thaliana*

Having established a stereochemically defined standard, we next monitored the endogenous levels of 11-OH-JA in *A. thaliana* (Col-0) leaves following wounding. We employed UPLC–MS/MS to detect 11-OH-JA and compared its accumulation to that of 12-OH-JA. As references, we employed synthetic standards of (3*R*,7*S*,11*R*)-11-OH-JA, (3*R*,7*S*,11*S*)-11-OH-JA, a mixture of (3*R*,7*S*)/(3*R*,7*R*)-11*R*-OH-JA, a mixture of (3*R*,7*S*)/(3*R*,7*R*)-11*S*-OH-JA, and a previously characterized mixture of (3*R*,7*S*)/(3*R*,7*R*)-12-OH-JA (5:95) ^18^.

Under optimized analytical conditions with a pentafluorophenyl UPLC column, we achieved sufficient resolution of all isomers: (3*R*,7*S*,11*R*)-11-OH-JA at Rt = 16.0 min, (3*R*,7*R*,11*R*)-11-OH-JA at Rt = 20.5 min, (3*R*,7*S*,11*S*)-11-OH-JA at Rt = 20.4 min, (3*R*,7*R*,11*S*)-11-OH-JA at Rt = 21.5 min, (3*R*,7*S*)-12-OH-JA at Rt = 19.3 min, and (3*R*,7*R*)-12-OH-JA at Rt = 22.2 min (**Figure 2**). However, the peaks of (3*R*,7*R*,11*R*)-11-OH-JA and (3*R*,7*S*,11*S*)-11-OH-JA overlapped under this condition.

**Figure 2.**
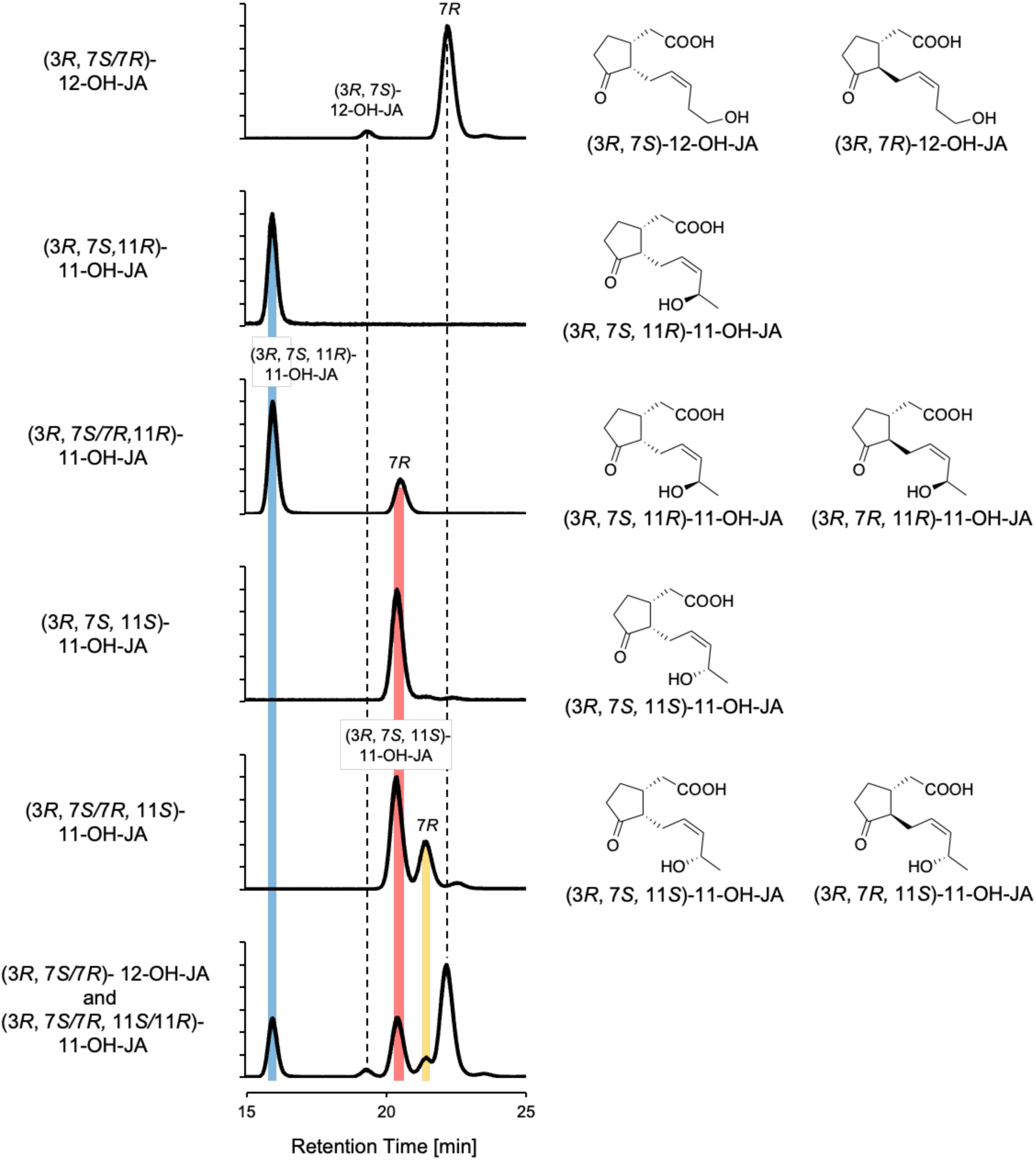
(**A**) UPLC-MS/MS chromatogram (*m*/*z* 225 > 59) of a mixture of synthetic (3*R*, 7*R*/*S*)-12-OH-JA, synthetic (*3R*, 7*S,* 11*S*)-11-OH-JA, a mixture of synthetic (*3R*, 7*R*/*S,* 11*S*)-11-OH-JA, synthetic (*3R*, 7*S,* 11*R*)-11-OH-JA, a mixture of synthetic (*3R*, 7*R*/*S,* 11*R*)-11-OH-JA, a mixture of (3*R*, 7*R*/*S*)-12-OH-JA and (*3R*, 7*R*/*S,* 11*R*/11*S*).

We observed the accumulations of OH-JAs in wounded leaves of wild-type *A. thaliana* (Col-0) at 60, 120, 300, and 480 min (**Fig. 3A**). The level of (3*R*,7*S*)-12-OH-JA started to accumulate from 60 min and peaked at 120 minutes post-wounding (**Figures 3A&3B**). In contrast, the four possible stereoisomers of 11-OH-JA, (3*R*,7*S*,11*R*)-11-OH-JA, (3*R*,7*R*,11*R*)-11-OH-JA, (3*R*,7*S*,11*S*)-11-OH-JA, and (3*R*,7*R*,11*S*)-11-OH-JA, exhibited delayed accumulation from 120 min, reaching maximal levels at 300 minutes post-wounding (**Figures 3A&3B**). Interestingly, both 11*R* and 11*S* stereoisomers were detected, suggesting that oxidation at the C11 position in wounded *A. thaliana* occurs in a non-stereoselective manner. Furthermore, 11-OH-JAs with both 7*S* and 7*R* configurations were found in plant extracts, although the 7*S* form is the biosynthetically derived stereoisomer. This observation is likely due to the inherent lability of the C7 stereocenter in 11-OH-JA. Notably, at 300 minutes post-wounding, the total level of 11-OH-JA (ca. 2,000 pmol/gFW) exceeded that of 12-OH-JA (ca. 770 pmol/gFW) (**Figure 3B**), consistent with previous GC-MS analyses indicating that 11-OH-JA is the major hydroxylated JA derivative in wounded *Arabidopsis* leaves (26). And these distinct time-course accumulation profiles raise the hypothesis that 11- and 12-hydroxylation of JA will proceed via separate biosynthetic pathways.

**Figure 3.**
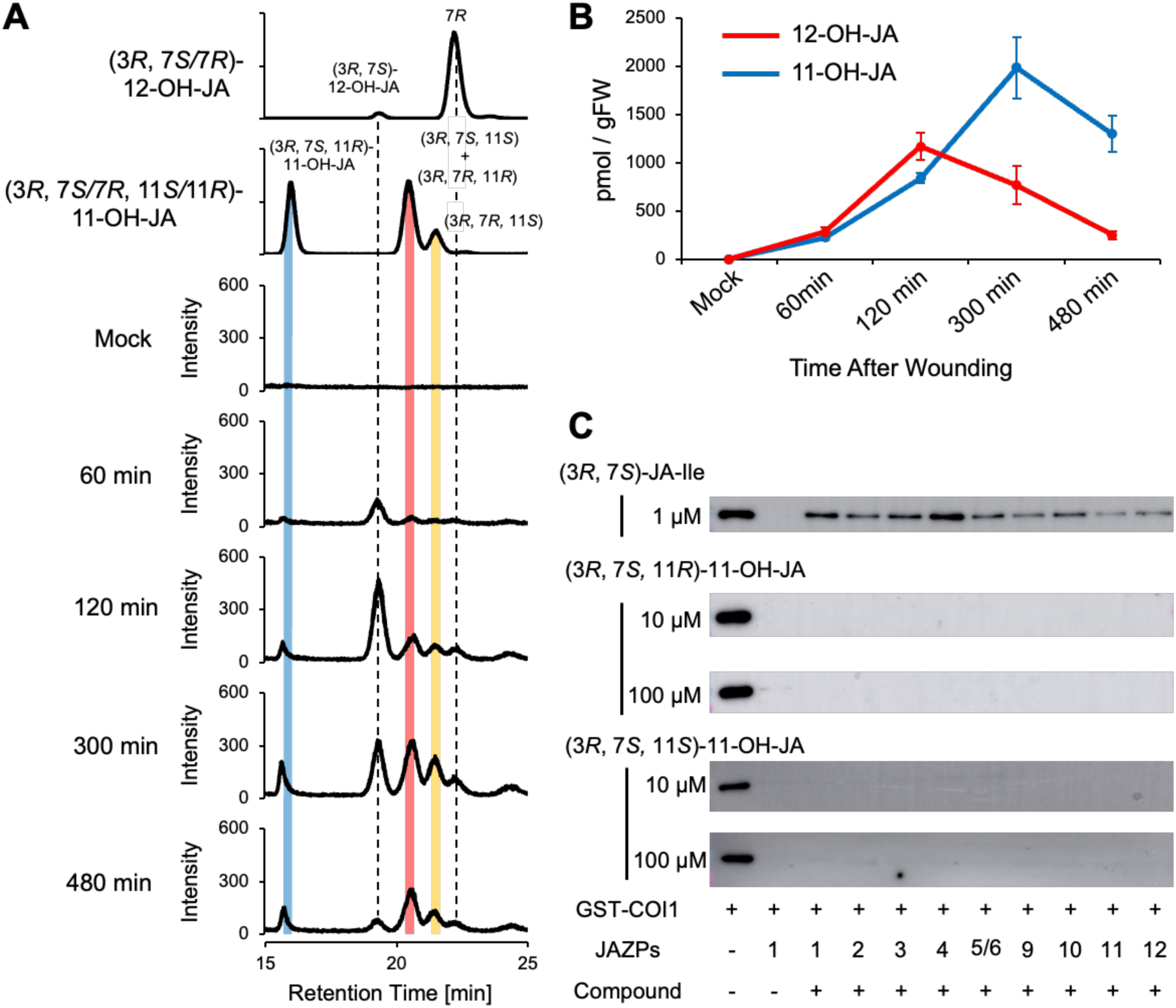
(**A**) Time course change of 11/12-OH-JAs accumulation in wounded *A.thaliana* leaves. 11/12-OH-JAs were extracted from the leaves of 5-week-old *A.thaliana* and analyzed using UPLC-MS/MS. Experiments are repeated three times with similar results. (**B**) Time-course of 11-OH-JA and 12-OH-JA accumulation in wounded leaves. Values are presented as mean ±SE (n = 4). (**C**) A pulldown assay of GST-COI1 with JAZPs was performed in the presence of (3*R*,7*S*)-JA-Ile (1 μM), (3*S*,7*S,* 11*R*)-11-OH-JA-Ile (10/100 μM), or (3*S*,7*S,* 11*S*)-11-OH-JA-Ile (10/100 μM). GST-COI1 bound to JAZP was pulled down with anti-fluorescein antibody and protein G beads, and analyzed by immunoblotting (anti-GST-HRP conjugate for detection of GST-COI1).

### Biological Assays using 11-OH-JA

Given that jasmonic acid (JA) elicits diverse defense-related and developmental responses via its bioactive conjugate, JA-Ile, we next assessed the ability of 11-OH-JA to bind the COI1–JAZ co-receptor complex, which is essential for perceiving bioactive jasmonates. A pull-down assay using GST-tagged COI1 (GST–COI1) and various JAZ degron peptides (JAZP1–6/9–12, containing Jas-motif required for interaction with JA-Ile) ^29,30^ revealed that (3*R*,7*S*,11*R*)-11-OH-JA and (3*R*,7*S*,11*S*)-11-OH-JA (up to 100 µM) did not promote COI1–JAZ interaction (**Fig. 3C and S6**). This result align with prior work on 12-OH-JA, which also lacks binding affinity for COI1–JAZ (16). Thus, similar to 12-OH-JA, 11-OH-JA acts as a non-bioactive catabolite of JA in *A. thaliana*.

### *In Silico* and *In Vitro* Analysis of JOX2-Mediated Oxidation of JA to 11-OH-JA

Multiple studies have implicated a family of 2-oxoglutarate-dependent dioxygenases, termed JASMONATE-INDUCED OXYGENASE/JASMONIC ACID OXIDASE (JAO/JOX), in the direct oxidation of JA ^21,31,32^. However, these earlier reports generally attributed the enzymatic product to “12-OH-JA” without explicitly ruling out 11-OH-JA, largely due to the absence of an authentic 11-OH-JA standard. To clarify the precise enzymatic product, we focused on JOX2, a key member JAO/JOX that is highly selective for JA ^32^.

We began by examining the crystal structure of JOX2 bound to (3*R*,7*R*)-JA (PDB ID: 6LSV) ^32^. Because the commercially available JA is typically a mixture of *cis*- and *trans*-isomers, the crystallized ligand was predominantly (3*R*,7*R*)-JA. To model the more biologically relevant (3*R*,7*S*)-JA, we performed *in silico* docking simulations that placed (3*R*,7*S*)-JA within the JOX2 active site (**Figures 4A–4D, S6**). This analysis revealed a set of conserved arginine residues (R225, R350, R354) forming hydrogen bonds with the carboxy and keto groups of JA, as well as a hydrophobic triad (F157, F317, F346) stabilizing the ligand through π-stacking and Van der Waals interactions ^32^. Additional hydrophobic interactions are provided by Y135, L142, L241, P247, and I353 (**Figures 4C and S6B-S6C**). These interactions ensure the precise positioning of (3*R*,7*S*)-JA, critical for oxidation. The iron center in JOX2 was located 4.04 Å and 4.37 Å from both C11 and C12 of JA, respectively (**Figures 4D**). According to the previous study ^32^, these distances are consistent with being within the range where oxidation can occur, starting with the abstraction of the first hydrogen atom, followed by rapid hydroxylation ^33^. However, the allylic nature of C11 makes it more prone to hydrogen abstraction, favoring the formation of 11-OH-JA.

**Figure 4.**
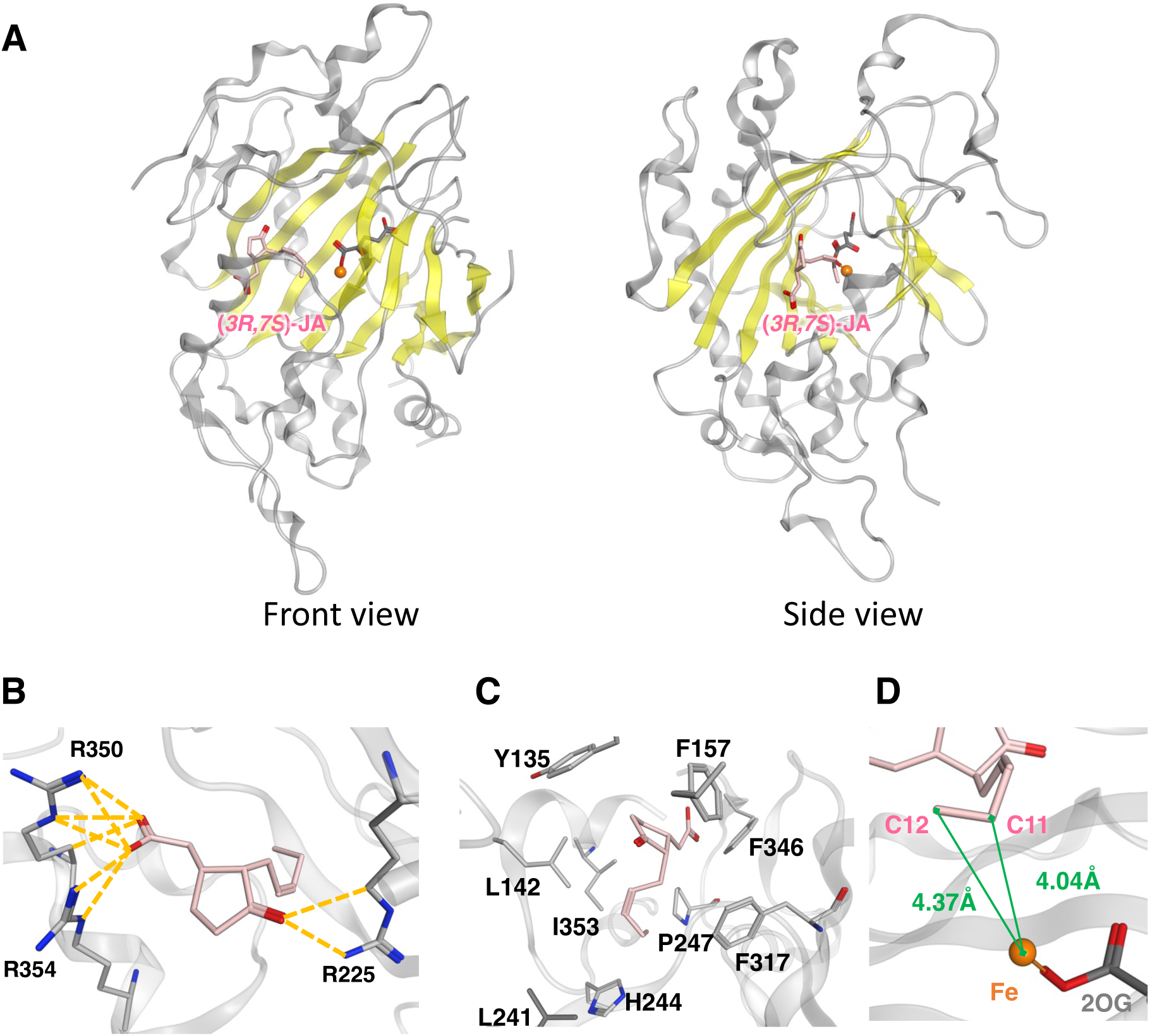
*In silico* docking models for (3*R*,7*S*)-JA as perceived by JOX2: (**A**) Docking model overview. Replaced (3*R*,7*S*)-JA is shown in pink sticks with JOX (6LSV). β seat is shown in yellow ribbons. (**B**) Hydrogen bonds between arginine residues and (3*R*,7*S*)-JA in the JOX2. (**C**) Hydrophobic interactions around (3*R*,7*S*)-JA in the JOX2. (**D**) Distances between the iron ion and the C12 and C11 carbon atoms of (3*R*,7*S*)-JA (pink). They are 4.37 Å and 4.04 Å, respectively. JOX2 and 2OG is represented in gray. Hydrogen bonds are shown as yellow dotted lines. Iron ion is showed as orange sphere. Distances are marked in green

To test this docking-guided prediction experimentally, we heterologously expressed full-length JOX2 (residues 1–371) in *Escherichia coli* and purified the recombinant protein (**Figures S7A, S7B**). When incubated with a mixture of (3*R*,7*S*)-JA and (3*R*,7*R*)-JA, the enzyme yielded major products at the same retention times with our synthetic 11-OH-JAs, while no 12-OH-JAs were detected (**Figures 5A, 5B**). These results directly contradict previous understanding that JOX2 catalyzes the formation of 12-OH-JA (19, 20, 32, 35).

**Figure 5.**
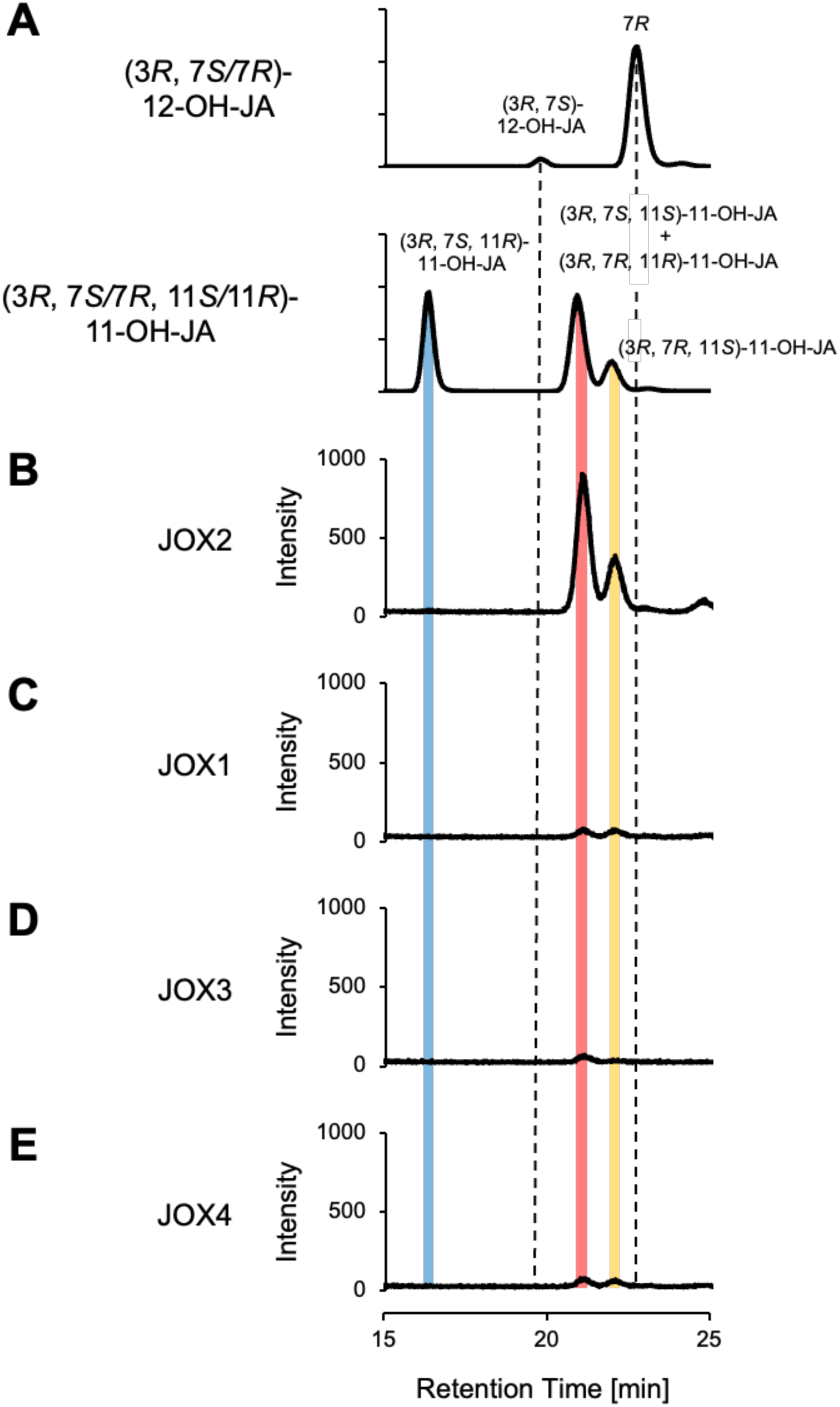
UPLC-MS/MS analyses of the products of heterologously expressed JOX1/2/3/4. (**A**) UPLC-MS/MS analyses of synthetic (*3R, 7S*)-12-OH-JA, (*3R, 7R*)-12-OH-JA, and (*3R, 7S, 11R/*11*S*)-11-OH-JA. UPLC-MS/MS analyses of products from the JOX-mediated enzymatic conversion of JA: (**B**) JOX2, (**C**) JOX1, (**D**) JOX3, and (**E**) JOX4.

We next asked whether other members of the JOX family (JOX1, JOX3, JOX4) also produce 11-OH-JA exclusively. Each isoform was expressed, purified (**Figures S7A, S7C–S7E**), and assayed with (3*R*,7*S*)-JA and (3*R*,7*R*)-JA in parallel reactions. Like JOX2, all three enzymes converted JA solely into 11-OH-JA, without 12-OH-JA production (**Figures 5C–5E**). These findings demonstrate a shared substrate specificity and catalytic outcome across the JOX family, firmly establishing 11-hydroxylation as their primary route for JA turnover.

### 11-OH-JA Accumulation Is Abolished in the *joxQ* Quadruple Mutant

To confirm the physiological relevance of JOX-mediated 11-hydroxylation in *planta*, we measured the accumulation of 11-OH-JA in the *joxQ* quadruple mutant, in which *JOX1*, *JOX2*, *JOX3*, and *JOX4* genes are impaired (**Figures 6A, 6B, and S8**) ^21^. Under mechanical wounding, the level of 11-OH-JA (ca. 2,000 pmol/gFW) in WT plants peaked at 300 min (**Figures 3A**). In contrast, the level of 11-OH-JAs in a *joxQ* mutant line significantly decreased, despite accumulating normal levels of 12-OH-JA (**Figures 6B**). This phenotype underscores the nonredundant role of JOX1/2/3/4 in catalyzing 11-hydroxylation of JA. Meanwhile, the retention of 12-OH-JA production in *joxQ* suggests that 12-hydroxylation proceeds through a distinct pathway, primarily CYP94-mediated oxidation of JA-Ile, followed by amide hydrolysis to 12-OH-JA (**Figure 1**).

**Figure 6.**
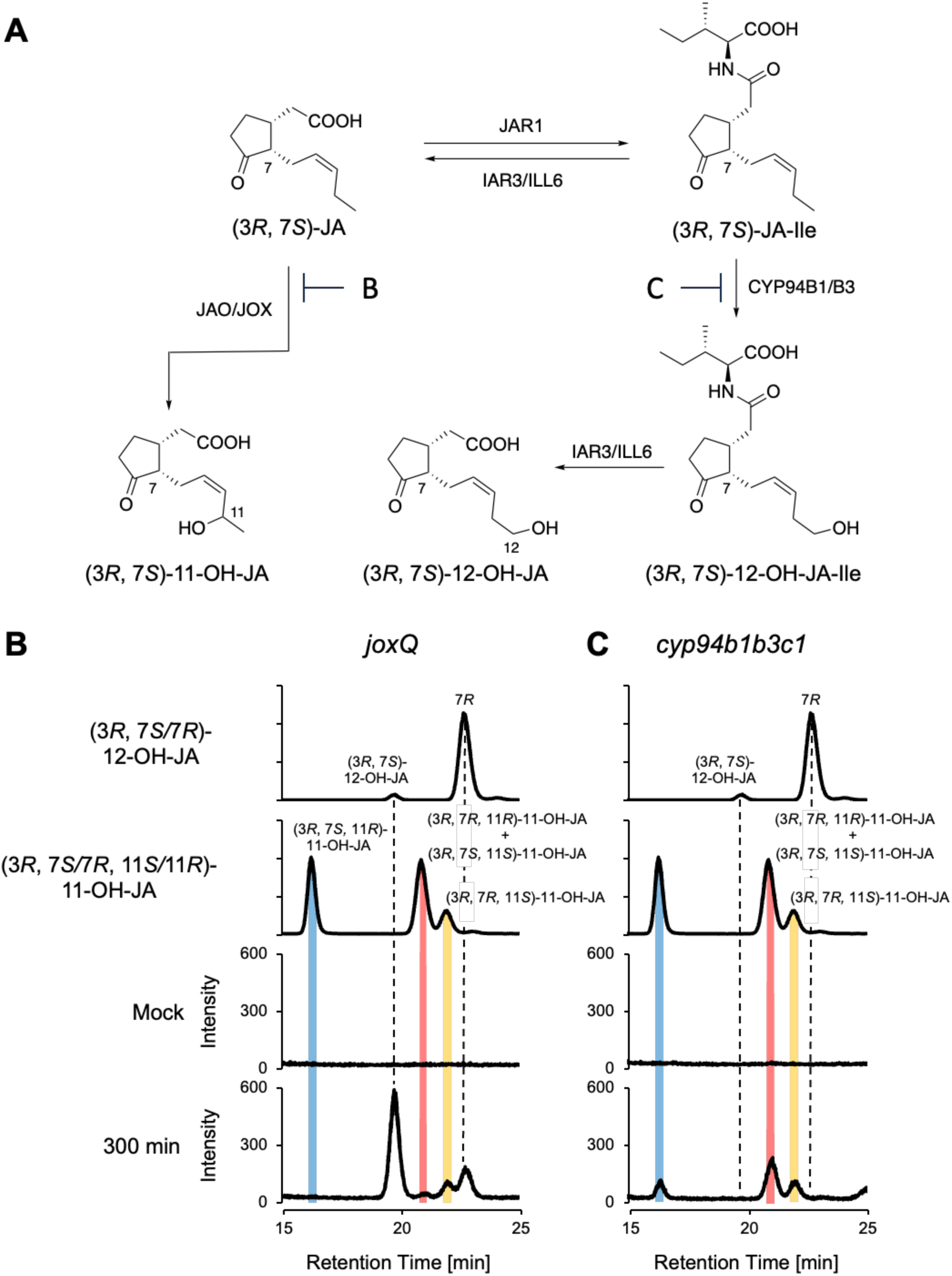
JA catabolite analyses in *joxQ* and *cypb1b3c1* mutant lines. **(A)** JA catabolite pathways were confirmed by using **(B**) *joxQ* and **(C**) *cypb1b3c1* mutant lines. UPLC-MS/MS analyses of 11/12-OH-JAs in wounded Col-0**, (B**) UPLC-MS/MS analyses of 11/12-OH-JAs in wounded Col-0 and wounded *joxQ*, (**C**) Time course changes in the accumulation of 11/12-OH-JAs in wounded Col-0 and wounded *cypb1b3c1*. The isocratic UPLC analysis was performed using water-MeOH (95:5, v/v) containing 0.2% acetic acid (v/v) with Poroshell 120 PFP 1.9 μm UPLC column (ϕ2.1 × 50 mm; Agilent Technologies). Multiple reaction monitoring (MRM) in the negative-ion mode was performed to identify 11-OH-JA and 12-OH-JA (*m*/*z* 225 > 59).

### 11-OH-JA Levels Are Unchanged in the *cyp94b1b3c1* Triple Mutant

Finally, we examined whether CYP94 enzymes, which are responsible for stepwise oxidation of JA-Ile to 12-OH-JA-Ile and 12-COOH-JA-Ile (9–14), might also contribute to 11-OH-JA formation via a possible “bypass” route—namely, 11-hydroxylation of JA-Ile followed by hydrolysis. To address this question, we profiled 11-OH-JA and 12-OH-JA in the *cyp94b1b3c1* triple mutant (**Figure 6A**) ^11,34^. As anticipated, 12-OH-JA levels were significantly reduced in this background (**Figure 6C**), reflecting impaired conversion of JA-Ile to 12-OH-JA-Ile. However, 11-OH-JA remained at wild-type levels (**Figure 6C**). This finding refutes the existence of a CYP94-dependent bypass that could produce 11-OH-JA. Instead, it affirms that JOX enzymes are solely responsible for oxidizing JA to 11-OH-JA, whereas CYP94 act upon JA-Ile to form 12-OH-JA-Ile (**Figure 1**).

In addition, we confirmed that the no detectable MS/MS (MRM) signal corresponding to 12- or 11-OH-JA-Ile (*m*/*z* 338→130; **Figure S9**) was observed in the extract of the wounded *cyp94b1cyp94b3cyp94c1* triple mutant. This observation demonstrated that the level of 11-OH-JA produced is under detection limits, and indicate that JOX-mediated 11-OH-JA is not converted to 11-OH-JA-Ile under wounding conditions.

In summary, our results establish that 11-OH-JA is synthesized exclusively by the JOX family through direct 11-hydroxylation of JA. Unlike previous understanding, none of the JOX isoforms appreciably yield 12-OH-JA. Coupled with our evidence that 11-OH-JA lacks jasmonate-like activity, these data highlight a major yet previously underrecognized route for JA inactivation in *A. thaliana*.

## DISCUSSION

In this study, we synthesized stereochemically pure (3*R*,7*S*,11*R*)-11-OH-JA and (3*R*,7*S*,11*S*)-11-OH-JA and leveraged it to uncover a previously underappreciated pathway in JA catabolism. By establishing UPLC analytical conditions that sufficiently separate 11-OH-JAs from 12-OH-JAs, we revealed that 11-OH-JAs are the major hydroxylated JA catabolite that accumulates in *A. thaliana* after wounding. Moreover, our biochemical and genetic experiments demonstrate that JOX1/2/3/4 specifically convert JA into 11-OH-JA (**Figures 1A**, **#2**). This finding contrasts with earlier understanding that JOX enzymes primarily produce 12-OH-JA ^21,22,32,35,36^, highlighting the importance of chemically synthesized authentic reference compounds and careful analytical separation in elucidating hormone catabolism pathways.

Our time-course analyses indicate that 12-OH-JA begins to accumulate around 60 min post-wounding, whereas 11-OH-JA levels rise later (300 min) (**Figure 3A**). This stepwise pattern underscores the distinct enzymatic routes leading to 11-OH-JA and 12-OH-JA in *A. thaliana*. Specifically, JOX-mediated 11-hydroxylation of JA emerges as a late step in diverting JA away from bioactive JA-Ile. In contrast, 12-OH-JA mostly arises from JA-Ile via CYP94-dependent oxidation (to form 12-OH-JA-Ile) followed by hydrolysis (**Figure 1**). The decreased level of 11-OH-JA in the *joxQ* quadruple mutant (**Figure 6B**), which lacks JOX1/2/3/4, further confirms the role of JOX enzymes in 11-hydroxylation of JA. Notably, *cyp94b1b3c1* mutants remain fully capable of producing 11-OH-JA (**Figure 6C**), ruling out a bypass route via 11-hydroxylation of JA-Ile.

Our biochemical and structural analyses suggest why JOX family members favor the allylic C11 of JA over the more terminal C12. *In silico* docking simulations of JOX2 with (3*R*,7*S*)-JA indicate that C11 is optimally positioned for hydrogen abstraction and subsequent hydroxylation, yielding 11-OH-JA exclusively (**Figure 4**). The marked preference for C11 oxidation is also supported by *in vitro* assays in which recombinant JOX2 (as well as JOX1, JOX3, and JOX4) failed to produce 12-OH-JA (**Figure 5B**). This finding underscores the potential pitfalls: earlier studies likely misattributed 11-OH-JA to 12-OH-JA because the two compounds are coelute under conventional UPLC–MS/MS conditions ^21,22,32,35,36^.

Beyond defining a key oxidative route, our data establish 11-OH-JA as a biologically inactive metabolite. Like 12-OH-JA, 11-OH-JA shows no affinity for the COI1–JAZ co-receptor (**Figure 3B**). Thus, 11-OH-JA functions as a shunt product, terminating JA signaling by depleting the pool of JA available for conversion into bioactive JA-Ile. This inactivation mechanism parallels the CYP94-driven pathway that targets JA-Ile itself; however, the two routes operate on different substrates (JA vs. JA-Ile) and rely on different enzymes (JOX vs. CYP94). In doing so, *A. thaliana* deploys multiple, nonredundant oxidative strategies to fine-tune jasmonate levels in response to developmental cues and environmental stresses.

The prominence of 11-OH-JA accumulation after wounding suggests that JOX-mediated oxidation plays an especially important role in modulating JA signaling under stress. Once formed, 11-OH-JA may also serve as a precursor to further modifications—such as conjugation or oxidation—that remain to be fully characterized. Indeed, while 12-OH-JA is further metabolized to forms such as 12-hydroxy jasmonoyl sulfate (12-HSO_4_-JA) ^3,26,35,37^ or 12-hydroxy jasmonoyl glucoside (12-*O*-Glc-JA) ^3,26,38,39^ (**Figure S10**), effectively removing it from the JA catabolism pathway, while the fate of 11-OH-JA is largely unknown.

The existence of parallel catabolic routes suggests that rapid JA inactivation confers critical fitness advantages. JA-related defense responses, while essential for survival under biotic assault, can inhibit growth and reproduction ^8^. Accordingly, having multiple enzymes that deactivate both JA and JA-Ile helps balance defensive needs against developmental demands. The prevalence of 11-hydroxylated JA in other plant species ^23-26^ indicates that this mechanism is likely widespread. Future comparative studies may illuminate whether JOX-mediated 11-hydroxylation is similarly dominant in other species and how plants modulate the relative fluxes between 11- and 12-hydroxylated intermediates.

In summary, our work demonstrates that 11-hydroxylation of JA by JOX1/2/3/4 is a major and early route of jasmonate inactivation in *A. thaliana* (**Figure 1A**, **#2**). By highlighting the exclusive production of 11-OH-JA, these findings update the previous understanding that JOX enzymes produce 12-OH-JA, and may reposition 11-OH-JA within the overall framework of JA turnover in *A. thaliana*. More broadly, they expand our view of hormone turnover, revealing an additional layer of complexity in how plants attenuate JA responses. Our findings position jasmonate turnover within a layered signal-termination framework. Rather than relying on a single catabolic outlet, plants deploy at least two non-redundant routes that act on different nodes of the jasmonate biosynthetic pathway: oxidation of JA-Ile to 12-OH-JA-Ile and JA to 11-OH-JA. Functionally, such layering increases robustness and speed, distinct modules can be differentially wired to environmental inputs, thereby limiting the duration and amplitude of defense signaling that otherwise imposes growth costs. Conceptually, this echoes a wider understanding in biology: signals are often terminated by distributed safeguards including ligand degradation, receptor desensitization or destruction, and transcriptional shut-off, which together create fail-safe attenuation. In jasmonate bioscience, placing 11-hydroxylation next to JA-Ile oxidation expands this repertoire and suggests testable predictions about how plants tune the growth–defense trade-off under fluctuating stress.

## MATERIALS AND METHODS

### Chemical Syntheses of (3*R*,7*S*, 11*R*)-11-OH-JA and (3*R*,7*S*, 11*S*)-11-OH-JA

To a solution of diol **5** (18.5 mg, 62 umol) in acetone (15.0 mL) was added Jones reagent (4.0 M solution, 600 uL, 2.4 mmol) at -20 °C. After 1 h of stirring at -20 °C, *i*-PrOH (800 uL) was added to quench the remaining reagent. After 1 h of stirring at -20 °C, EtOAc/*n*-hexane (1/1, 10 mL) and H_2_O (20 mL) were added, and the water layer was extracted with EtOAc. The combined organic layers were washed with brine, dried over Na_2_SO_4_, and concentrated under reduced pressure. The residue was purified by silica gel medium-pressure chromatography (*n*-hexane+0.1%AcOH/EtOAc+0.1%AcOH = 84/16 to 0/100) to give a carboxylic acid intermediate (18.7 mg, 97%) as a colorless oil.

To a solution of the carboxylic acid intermediate (18.7 mg) in Et_2_O (10 mL) was added MgBr_2_•OEt_2_ (31.1 mg, 120 umol). The resulting white suspension was stirred at room temperature for 1.5 h and dried under reduced pressure. The residue was dissolved with 35% aq. MeOH containing 0.1% acetic acid and syringe-filtered through PTFE membrane. The crude product was directly purified by preparative RP-HPLC (ODS-HG-5 Φ20x250, 210 nm, 6.0 mL/min, eluent: 35% aq. MeOH + 0.1%AcOH) to give (*3R*,*7S*,*11R*)-OH-JA (t_R_ = 27 min, 2.34 mg, 17%) and (*3R*,*7S*,*11S*)-OH-JA (t_R_ = 32 min, 2.92 mg, 21%) as a colorless amorphous solid.

(*3R*,*7S*,*11R*)-11-OH-JA: ^1^H NMR (400 MHz, CD_3_OD) 8_H_: 5.46-5.40 (m, 2H), 4.63-4.55 (m, 1H), 2.86-2.75 (m, 1H), 2.49-2.34 (m, 3H), 2.32-2.21 (m, 2H), 2.21-2.01 (m, 3H), 1.92-1.82 (m, 1H), 1.20 (d, *J* = 6.0 Hz, 3H); ^13^C NMR (100 MHz, CD_3_OD, 35 °C) δ_C_: 221.4, 176.6* (* determined by HMBC trace, see SI), 136.4, 128.5, 64.2, 53.8, 37.2, 36.1, 26.7 (2C), 24.2, 23.7; HRMS (ESI, negative) m/z [M-H]^-^ Calcd. for C_12_H_17_O_4_^-^: 225.1132, Found : 225.1130.

(*3R*,*7S*,*11S*)-11-OH-JA: ^1^H NMR (400 MHz, CD_3_OD) 8_H_: 5.49-5.39 (m, 2H), 4.63-4.54 (m, 1H), 2.87-2.75 (m, 1H), 2.53-2.29 (m, 3H), 2.29-2.20 (m, 2H), 2.20-1.95 (m, 3H), 1.92-1.81 (m, 1H), 1.19 (d, *J* = 6.4 Hz, 3H); ^13^C NMR (100 MHz, CD_3_OD) δ_C_: 221.2, 175.7* (* determined by HMBC trace, see SI), 135.5, 127.2, 63.3, 52.9, 36.0, 35.0, 33.9, 25.6, 23.3, 22.8; HRMS (ESI, negative) m/z [M-H]^-^ Calcd. for C_12_H_17_O_4_^-^: 225.1132, Found : 225.1145.

Detailed materials and methods of chemical syntheses are described in attached SI.

### Determination of Stereochemistry of (3*R*,7*S*, 11*R*)-11-OH-JA and (3*R*,7*S*, 11*S*)-11-OH-JA by modified Mosher’s method

Single stereoisomer of **3** was isolated from a mixture of (*3R, 7S, 11R/S*)-**3** by multiple silica gel medium-pressure chromatography. From a mixture of (*3R, 7S, 11R/S*)-**3** (39.2 mg), (*3R, 7S, 11R*)-**3** (14.2 mg) and (*3R, 7S, 11S*)-**3** (17.5 mg) were isolated. (*3R, 7S, 11S*)-3-(*R*/*S*)-MTPA ester was synthesized and the stereochemistry at the C11 was determined as (*S*) from the chemical shift difference (8^S^-8^R^) of the thus obtained MTPA esters (**Figure S4A**). Detailed materials and methods of chemical syntheses are described in attached SI.

To determine the stereochemistry at the C11 of RP-HPLC isolated two stereoisomers of (*3R,7S, 11R/S*)-11-OH-JA, (*3R,7S, 11S*)-11-OH-JA was synthesized from (*3R,7S, 11S*)-**3** (**Figure S4B**). Detailed materials and methods of chemical syntheses are described in attached SI. The synthesized (*3R,7S, 11S*)-11-OH-JA was subjected to UPLC-MS/MS analysis to clarify the stereochemistry of (*3R,7S, 11R*)-11-OH-JA and (*3R,7S, 11S*)-11-OH-JA (**Figure S4C**).

### Plant materialas

*Arabidopsis thaliana* Col-0 is the genetic background of wild-type and mutant lines used in this study. The seeds of the *cyp94b1b3c1* triple mutant was kindly provided by Prof. Thierry Heitz (University of Strasbourg, France). The seed of *joxQ* was kindly provided from Prof. Guido Van den Ackerveken (Utrecht University). All seeds were grown on Jiffy-7 (Sakata Seed Corporation, Kanagawa, Japan) in a Biotron LPH-240SP growth chamber (Nippon Medical & Chemical Instruments Co., Ltd., Osaka, Japan) at 22 °C under 16 h light (ca. 2000 lux) and 8 h dark conditions. The humidity was set to 50%.

### UPLC-MS/MS analyses of endogenous 11-OH-JAs and 12-OH-JAs in *A. thaliana* and the mutante lines

The leaves (ca. 100–200 mg FW) of the 5-week-old *A. thaliana* Col-0 were nipped tightly with a pair of tweezers. Damaged leaves were harvested at 0, 15, 30, 60, and 300 min after wounding. Each sample was weighed and frozen in liquid nitrogen. The frozen sample was ground using a Micro Homogenizing System (Micro Smash™; TOMY, Tokyo, Japan) with a zirconia bead and extracted in 1.5 mL EtOH for 16 h at 4 °C. After centrifugation at 4,300 rpm for 10 min, the supernatants were collected. The volatile components of the extract were removed by using nitrogen blowdown evaporator, and a solution of 150 μL 80% aq. MeOH was then added to the residue. The mixture was applied to an SPE C18 cartridge column (Sep-Pak C18 Plus Short Cartridge, 360 mg, Waters, Milford, MA, USA) and the column was eluted with a solution of 80% aq. MeOH (700 μL × 3). The volatile components and water in the eluents were removed by using nitrogen blowdown evaporator. The residue was dissolved with a solution of 100 μL MeOH, and the sample was examined by LC-MS/MS using an Ultivo Triple Quadrupole LC/MS system (Agilent Technologies, Santa Clara, CA, USA) operating in the negative mode. Poroshell 120 PFP 1.9 μm (Φ2.1 × 50 mm; Agilent Technologies) was used at 23 °C and a flow rate of 0.300 mL/min. The elution was performed using water-MeOH (95:5, v/v, isocratic), both containing 0.2% acetic acid (v/v). The liquid chromatograms were extracted by multiple reaction monitoring with the following ion transitions: 11-OH-JA 225 > 59 and 12-OH-JA 225 > 59.

### *In silico* Docking study of (3*R*,7*S*)-JA with JOX1-4

The initial structures of the (3*R,*7*R*)-JA with JOX2 that were obtained were based on a crystal structure (PBD ID: 6LSV) ^32^. Defects in the crystal structure were corrected using Structure Preparation in MOE (version 2024.06, Chemical Computing Group Inc., Montreal, QC, Canada) and hydrogen atoms were added and structurally optimized. (3*R*,7*S*)-JA were modeled with ChemDraw and Chem3D software (PerkinElmer, Inc., Waltham, MA, USA) and were subjected to moe to add partial charges and minimize their energy. The best-docking pose, those which have similar interactions as (*3R,7R*)-JA on a crystal structure, was selected. The homology modeling of JOX1/3/4 were obtained based on the crystal structure of the JOX2-2OG-iron-JA complex (PDB ID: 3OGL) using moe. Docking study was performed similarly as JOX2.

### Enzymatic Activity assay of JOX1/2/3/4 with UPLC-MS/MS analyses

The 20 μM (-)-JA was dissolved in 10 mM HEPES buffer (pH 7.4), containing 150 mM NaCl, 5 mM dithiothreitol (DTT), 2 mM α-ketoglutarate, 5 mM ascorbic acid, and 1 mM FeSO_4_. To this solution, JOX1/2/3/4 (10 μg) was added respectively and the solution was adjusted to 100 μl with storage buffer. The sample solution was incubated at 30 °C for 2 h and then preserved on ice. After filtration, sample was diluted 10-fold with methanol and 10 µL of the diluted sample solution was subjected to UPLC-MS/MS analysis observing produced 11-OH-JA or 12-OH-JA. A negative mode UPLC-MS/MS analysis was performed on an Ultivo Triple Quadrupole LC/MS system (Agilent Technologies) equipped with Poroshell 120 PFP 1.9 μm UPLC column (ϕ2.1 × 50 mm; Agilent Technologies) (column temperature: 23 °C, flow rate: 0.300 mL/min). The isocratic UPLC analysis was performed using water-MeOH (95:5, v/v) containing 0.2% acetic acid (v/v). The chromatograms were extracted by multiple reaction monitoring with the following ion transitions: 11-OH-JA 225 > 59 and 12-OH-JA 225 > 59.

## Acknowledgments

The seeds of the *cyp94b1b3c1* triple mutant was kindly provided by Prof. Thierry Heitz (University of Strasbourg, France). The seed of *joxQ* was kindly provided from Prof. Guido Van den Ackerveken (Utrecht University). And we also thank to Prof. Thierry Heitz (University of Strasbourg, France) for valuable discussion on our manuscript.

## Funding

This work was financially supported by Grant-in-Aid for Scientific Research from JSPS (Japan) projects No. 23H00316, JPJSBP120239903, and a Grant-in-Aid for Transformative Research Areas (A) “Latent Chemical Space” [JP23H04880 and JP23H04883] from the Ministry of Education, Culture, Sports, Science and Technology, Japan (to MU).

## Author contributions

Conceptualization, M.U.; methodology, M.U., K. M., T.K., N. K.; validation, M.U., T.K., and Y. N.; formal analysis, K.M., M.M., T.K., R.G., T.O.; investigation, K.M., M.M., T.K., R.G., T.O.; resources, K.M., T.K., R.G., T. O., N. K., and H. M.; data curation, M.U. and T. K.; writing—original draft preparation, M.U.; writing—review and editing, M.U. and T.K.; visualization, K.M., M.M., T.K., T.O.; supervision, M.U.; project administration, M.U.; funding acquisition, M.U. All authors have read and agreed to the published version of the manuscript.

## Competing interests

The authors declare no competing interests.

## Data availability statement

The data generated in this study are provided in the Supplementary Information.

## Notes

### Competing Interest Statement

The authors have declared no competing interest.

