## Supplementary Information for "(3*R*, 7*S*)-11-hydroxy-jasmonic acid is a major oxidative shunt product of jasmonate catabolism in *Arabidopsis thaliana*"

**Title**

**Contents:**

Fig. S1-S9 S2-S13

Supplementary Materials and Methods S14-S34

[1]H-, [13]C-, and 2D NMR spectra SX34-S48

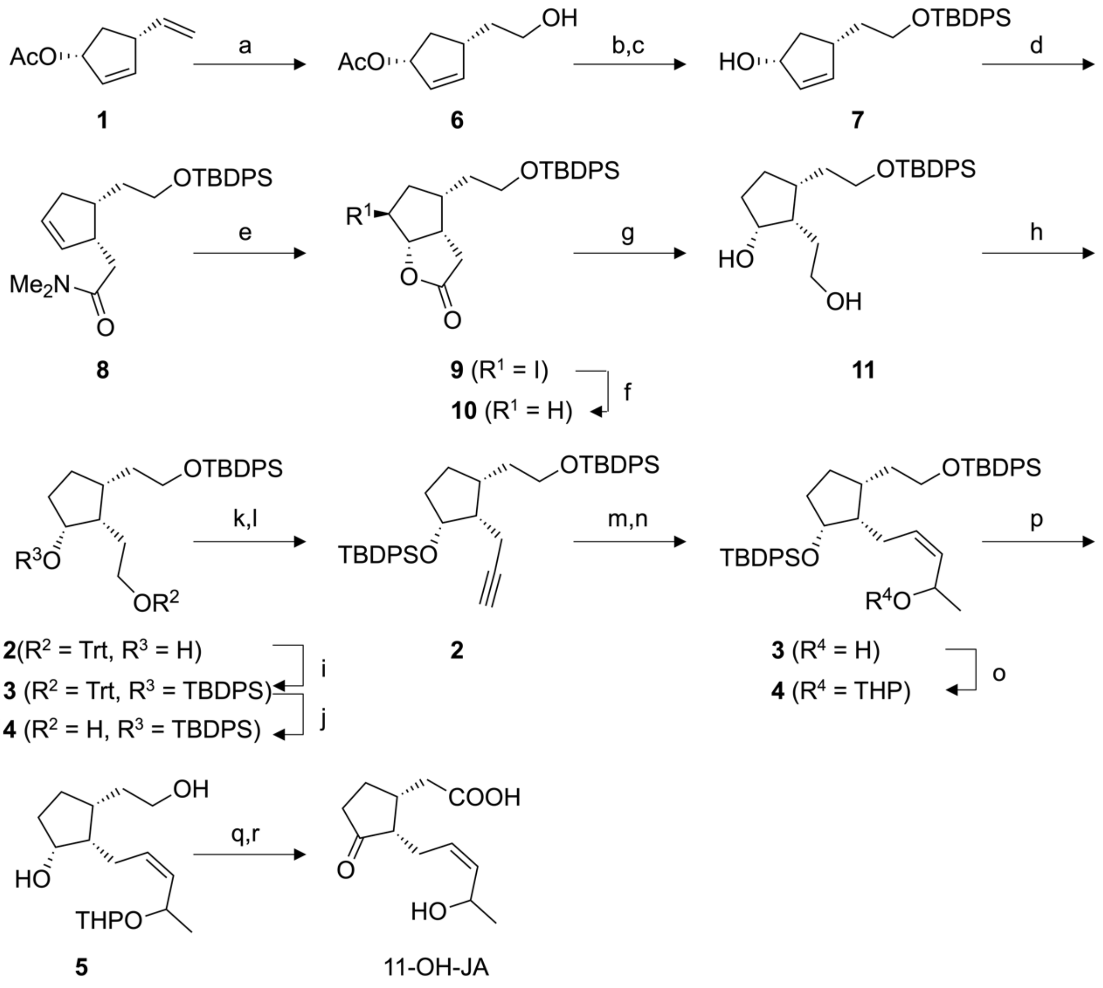


**Fig. S1 Synthetic scheme of 11-OH-JA**: (a) $Cy_2BH$, THF, 0ºC; $NaBO_3$, $H_2O$, 0 ºC, 60%; (b) TBDPSCl, imidazole, DMF; (c) NaOH, MeOH/$H_2O$ , 66% (2 steps); (d) $MeC(OMe)_2NMe_2$, xylene, 150 ºC, 56%; (e) $I_2$, $KH_2PO_4$, $Na_2HPO_4$, THF/$H_2O$ , 84%; (f) *n*$Bu_3SnH$, AIBN, benzene, reflux; (g) LiAlH, THF, 0 ºC, 93% (2 steps); (h) TrtCl, DMAP, $Et_3N$, $CH_2Cl_2$, 74%; (i) TBDPSCl, imidazole, DMF, 86%; (j) *p*-TsOH•$H_2O$, $CH_2Cl_2$, MeOH, 77%; (k) DMSO, $(COCl)_2$, DCM, -65 ºC; $Et_3N$, -65 ºC to rt; (l) Ohira-Bestmann reagent, $K_2CO_3$, MeOH, 69% (2 steps); (m) *n*BuLi, acetaldehyde, THF, -78 ºC, 69%; (n) $H_2$, Pd/$BaSO_4$, quinoline, MeOH, 89%; (o) DHP, *p*-TsOH•$H_2O$, $CH_2Cl_2$, 98%; (p) TBAF, THF 91%; (q) Jones reagent, acetone, -20 ºC, 96%; (r) $MgBr_2$, $Et_2O$, 38%.

**A**

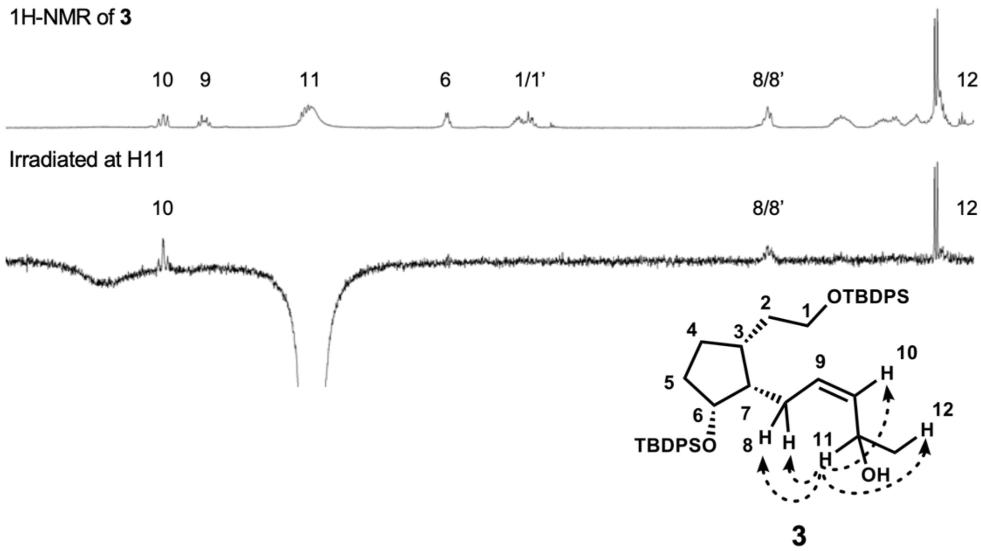
1H-NMR of 3
10
9
11
6
1/1'
8/8'
12
Irradiated at H11
10
8/8'
12
OTBDPS
TBDPSO
OH
3

**B**

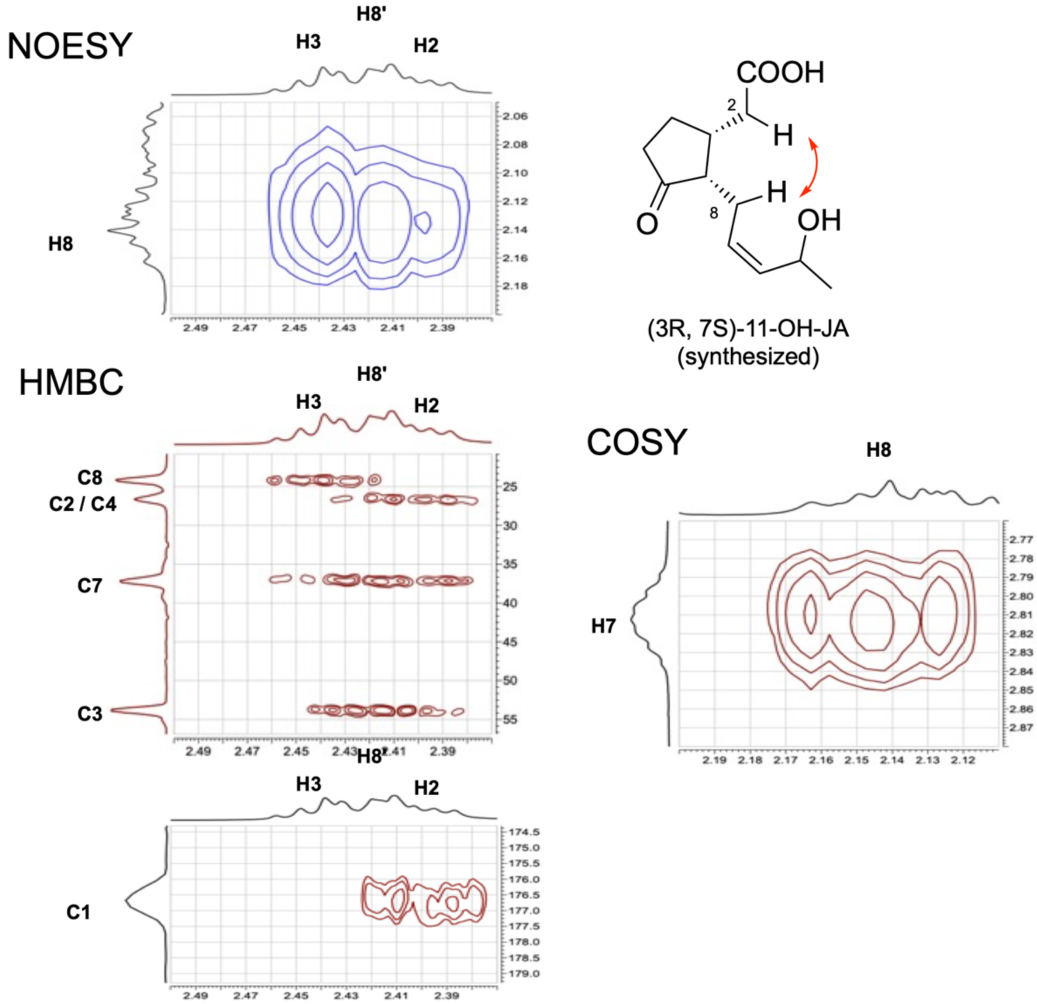
NOESY
H3
H8'
H2
H8
COOH
OH
(3R, 7S)-11-OH-JA
(synthesized)
HMBC
H3
H8'
H2
C8
C2 / C4
C7
C3
C1
COSY
H8
H7

**Fig. S2** (**A**) Differential NOE experiment on intermediate **3**. 1D-differential-noe spectrum was measured in pyridine-*d5* (64 scans). H11 (δ 5.06) was selectively irradiated. The key NOE correlation between H11 and H8/H8' was 0.3%. (**B**) Expanded view of NOESY (700 MHz) cross peak of the synthesized (3*R*, 7*S*)-11-OH-JA between H2 and H8 and related HSQC and HMBC (700 MHz for $^{1}$H and 175 MHz for $^{13}$C) used to assign H2.

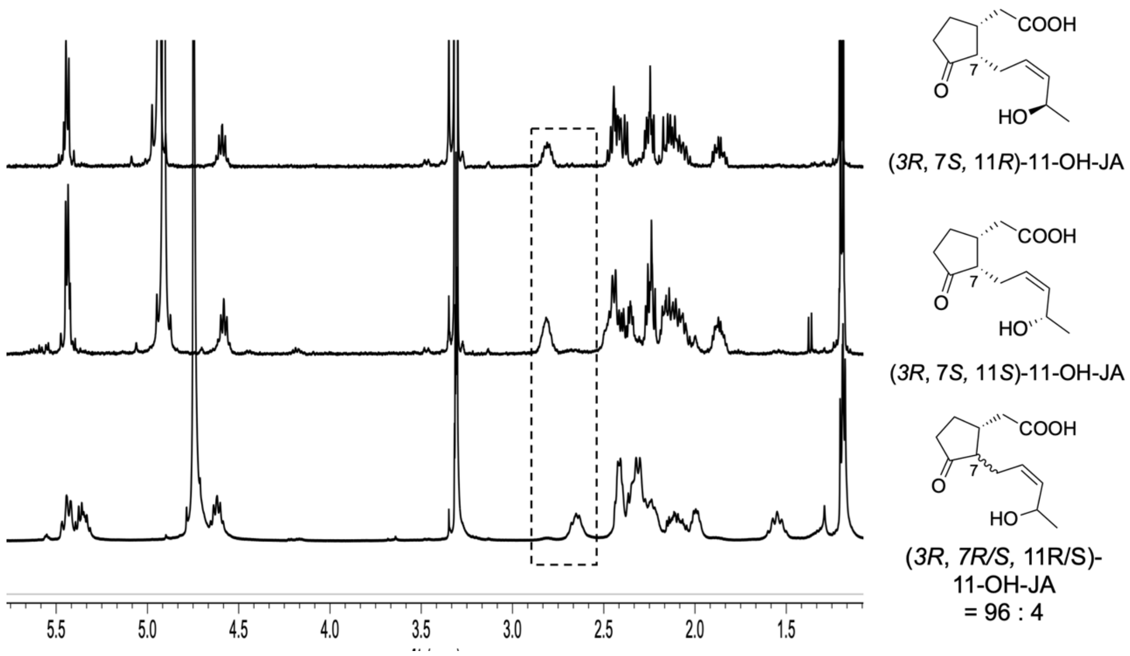


**Fig. S3.** Comparison of [1]H NMR between (3*R*, 7*S*, 11*R*)-11-OH-JA (top), (3*R*, 7*S*, 11*S*)-11-OH-JA (middle), and a separately synthesized mixture of (3*R*, 7*S*, 11*R*/11*S*)-11-OH-JA and (3*R*, 7*R*, 11*R*/11*S*)-11-OH-JA (4:96). The region within the dotted line was shown in **Figure 1C**.

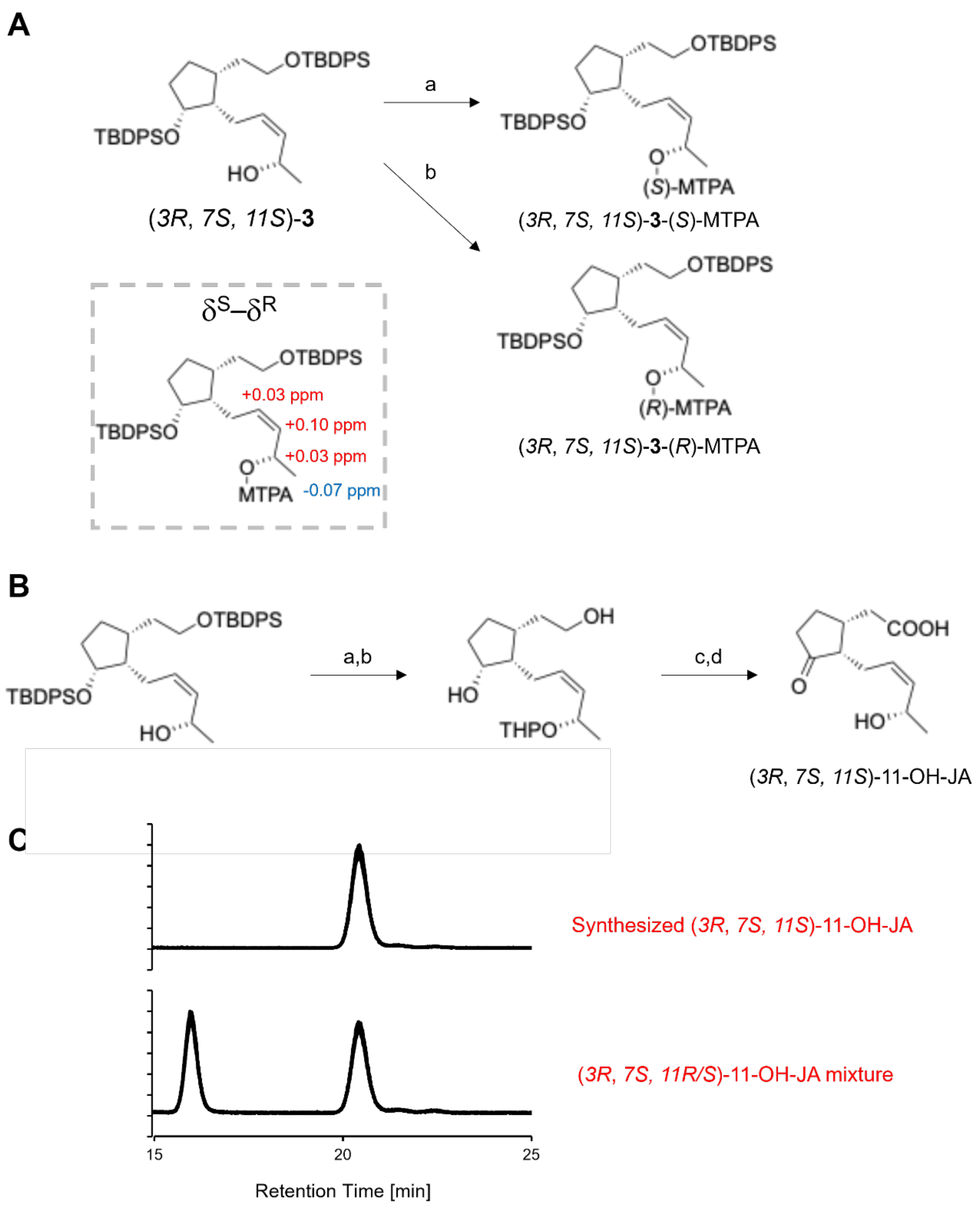


**Fig. S4.** (**A**) Determination of the stereochemistry at the C11 position of one isolated stereoisomer of **3** by modified Mosher method : (a) (*R*)-MTPA-Cl, pyridine, DCM; (b) (*S*)-MTPA-Cl, pyridine, DCM; (B) Synthesis of (*3R*, *7S, 11S*)-11-OH-JA from (*3R*, *7S, 11S*)-**3** : (a) DHP, *p*-TsOH•$H_2O$, $CH_2Cl_2$, 91%; (b) TBAF, THF 89%; (c) Jones reagent, acetone, -20 ºC, 78%; (d) $MgBr_2$, $Et_2O$ 44%; (C) Comparison of LC-MS/MS chromatogram suggested that the latter peak of (*3R*, *7S, 11R/S*)-11-OH-JA mixture was (*3R*, *7S, 11S*)-11-OH-JA isomer.

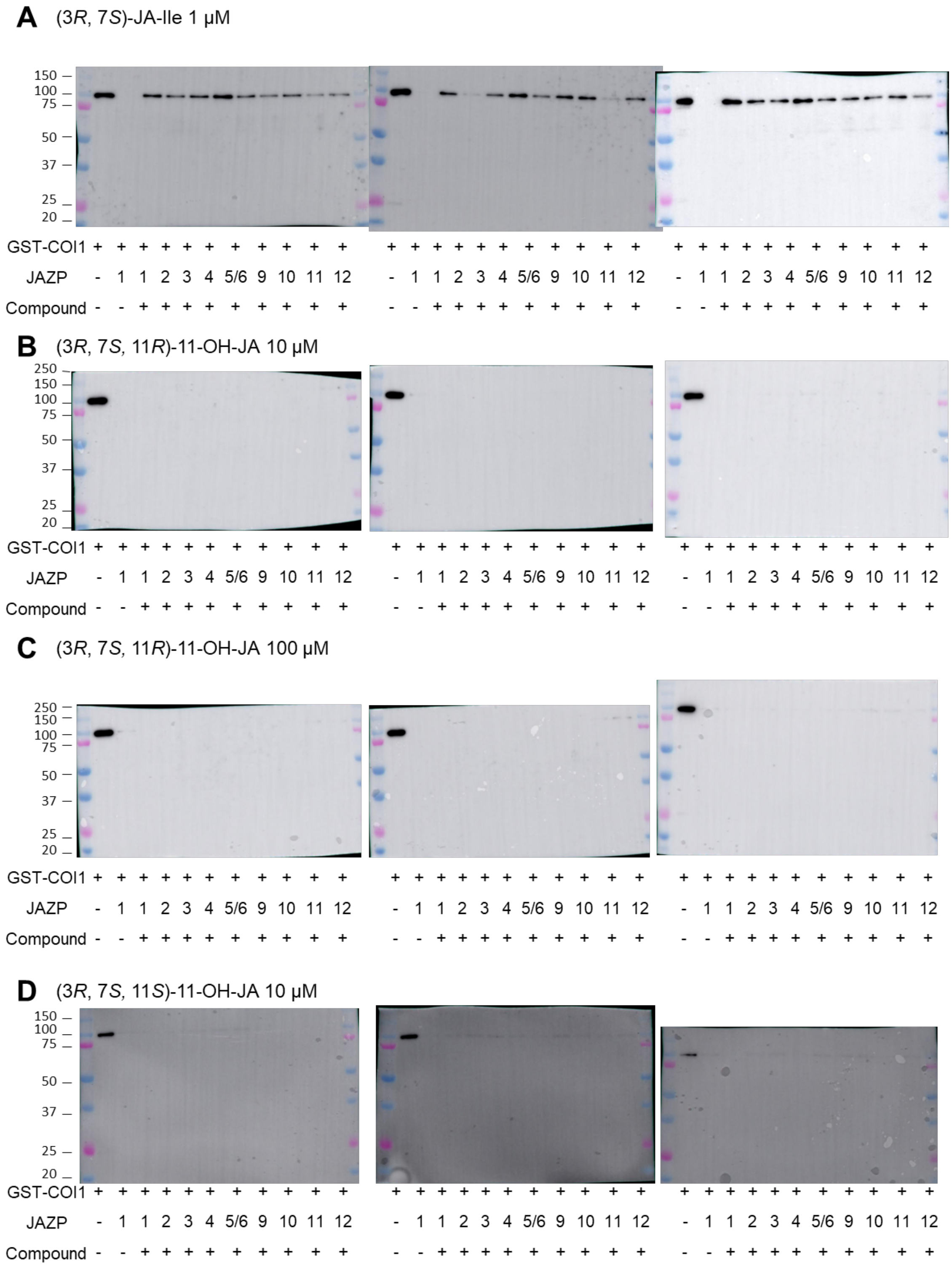
A (3R, 7S)-JA-Ile 1 μM
150
100
75
50
37
25
20
GST-COI1 + + + + + + + + + + + + + + + + + + + + + + + + + + + + + + + + +
JAZP - 1 1 2 3 4 5/6 9 10 11 12 - 1 1 2 3 4 5/6 9 10 11 12 - 1 1 2 3 4 5/6 9 10 11 12
Compound - - + + + + + + + + + - - + + + + + + + + + - - + + + + + + + + +
B (3R, 7S, 11R)-11-OH-JA 10 μM
250
150
100
75
50
37
25
20
GST-COI1 + + + + + + + + + + + + + + + + + + + + + + + + + + + + + + + + +
JAZP - 1 1 2 3 4 5/6 9 10 11 12 - 1 1 2 3 4 5/6 9 10 11 12 - 1 1 2 3 4 5/6 9 10 11 12
Compound - - + + + + + + + + + - - + + + + + + + + + - - + + + + + + + + +
C (3R, 7S, 11R)-11-OH-JA 100 μM
250
150
100
75
50
37
25
20
GST-COI1 + + + + + + + + + + + + + + + + + + + + + + + + + + + + + + + + +
JAZP - 1 1 2 3 4 5/6 9 10 11 12 - 1 1 2 3 4 5/6 9 10 11 12 - 1 1 2 3 4 5/6 9 10 11 12
Compound - - + + + + + + + + + - - + + + + + + + + + - - + + + + + + + + +
D (3R, 7S, 11S)-11-OH-JA 10 μM
150
100
75
50
37
25
20
GST-COI1 + + + + + + + + + + + + + + + + + + + + + + + + + + + + + + + + +
JAZP - 1 1 2 3 4 5/6 9 10 11 12 - 1 1 2 3 4 5/6 9 10 11 12 - 1 1 2 3 4 5/6 9 10 11 12
Compound - - + + + + + + + + + - - + + + + + + + + + - - + + + + + + + + +

**E** (3*R*, 7*S*, 11*S*)-11-OH-JA 100 μM

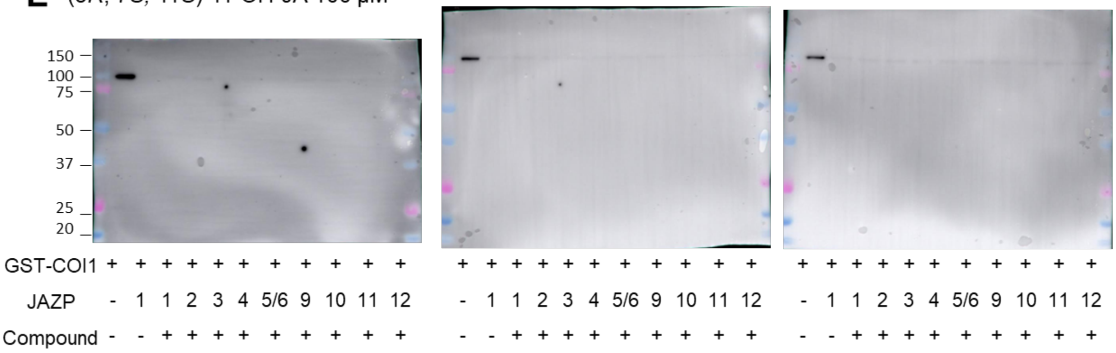


**F** No compunds

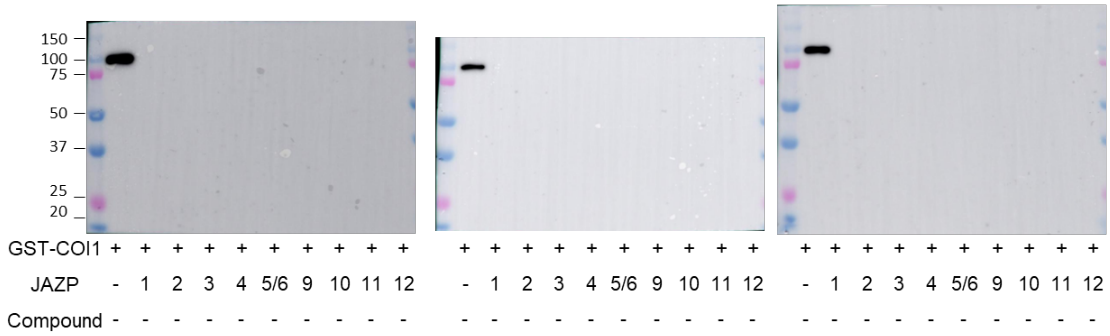


**Fig. S5** Uncropped images and three independent pull-down results shown in Figure 2C. Pull-down assays between GST-COI1 and Fl-JAZPs1-6/9-12 under (3R,7S)-JA-Ile (1 μM, A), (3R, 7S, 11R)-11-OH-JA (10/100 μM, B,C for each result), (3R, 7S, 11S)-11-OH-JA (10/100 μM, D,E for each result) or no compounds (F for each result) . GST-COI1 which bound to JAZPs was pulled down with anti-fluorescein antibody and protein A beads. They are analyzed by immunoblotting (anti-GST-HRP conjugate for detection of GST-COI1).

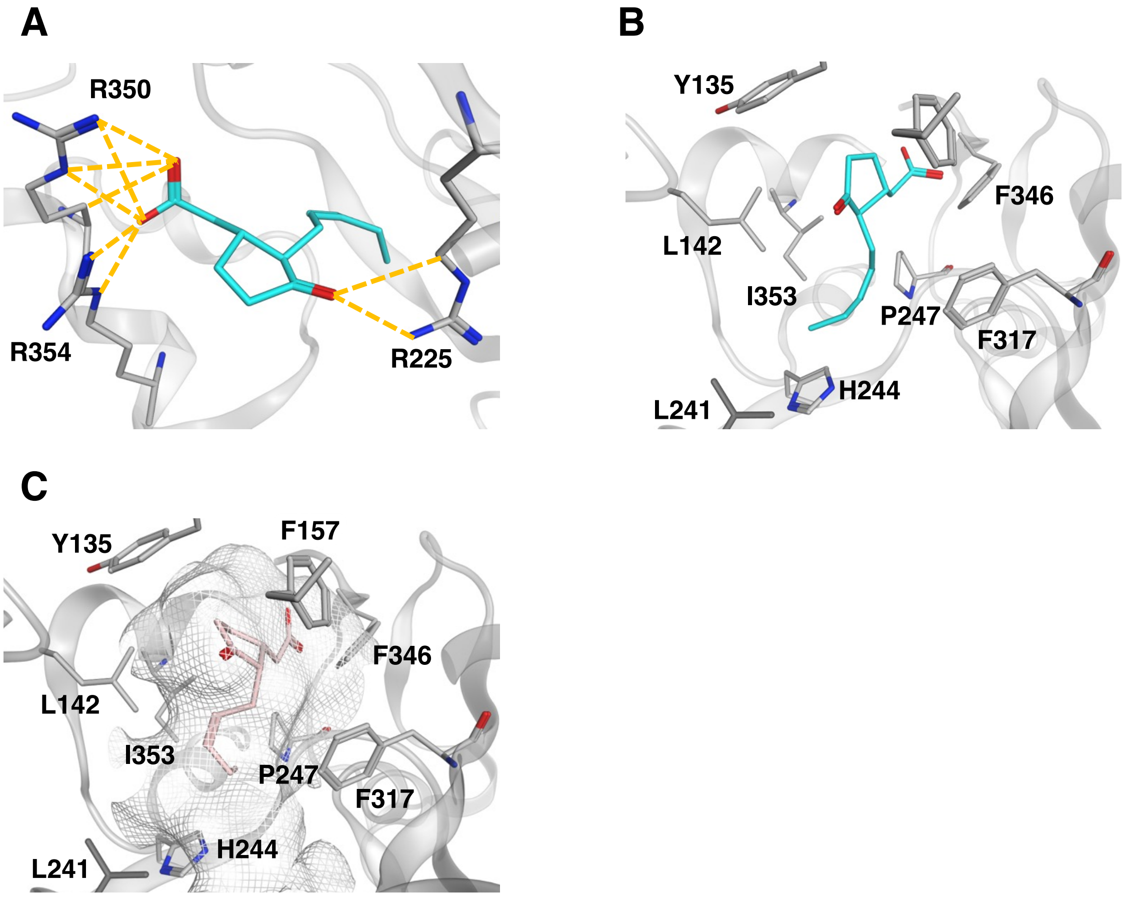


**Fig. S6** (**A**) Hydrogen bond network and (**B**) hydrophobic interactions in the crystal structure of (3*R*,7*R*)-JA (cyan) and JOX2 (gray) (PDB: 6LSV). (**C**) Hydrophobic residues form the ligand-binding pocket (shown in gray mesh).

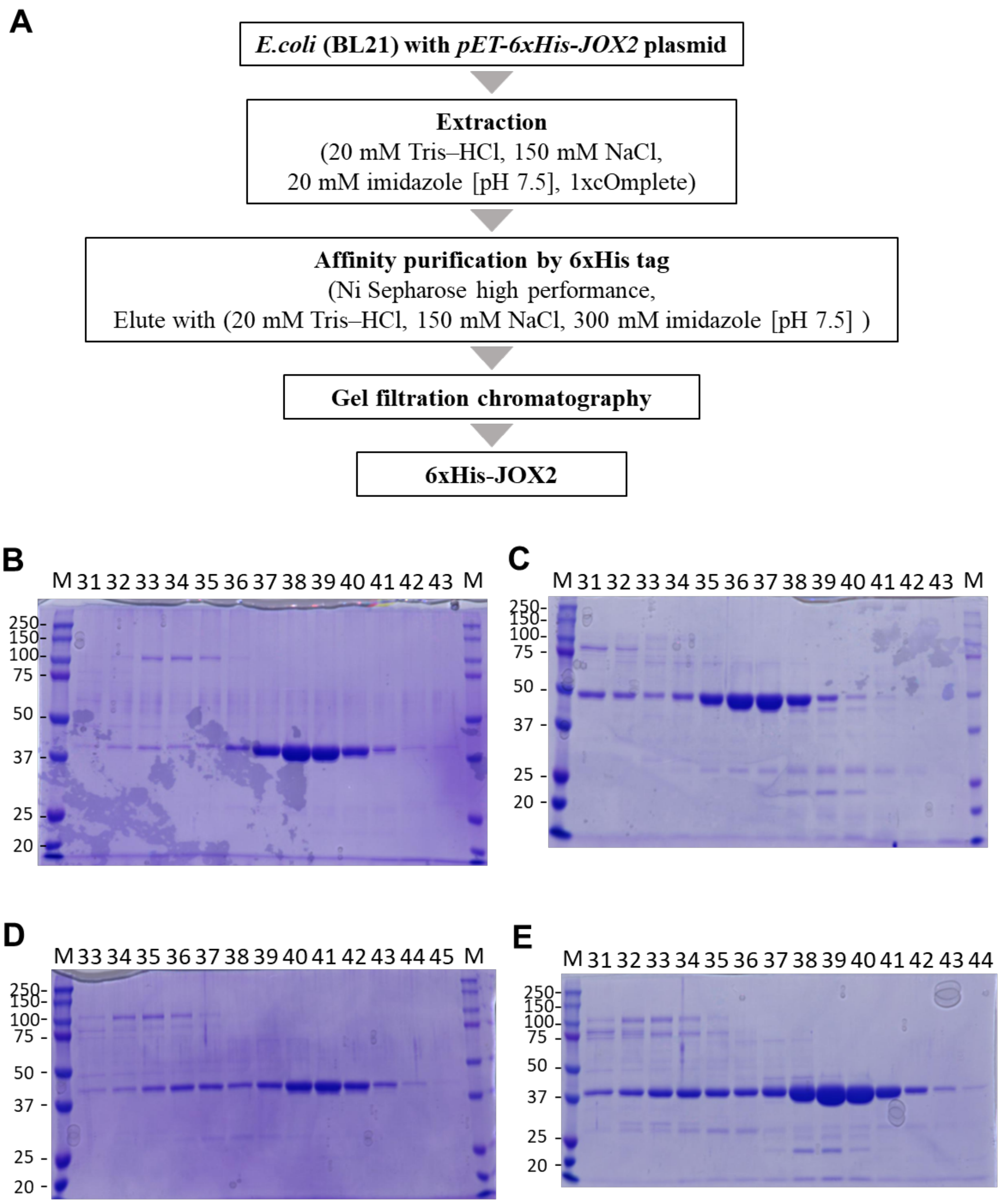


**Fig. S7 Expression and purification of 6xHis-JOX1/2/3/4** (**A**) A schematic representation of JOX2 expression and purification. JOX1/3/4 were similarly expressed and purified. (**B**) CBB-stained SDS-PAGE analysis of 6xHis-JOX2 after gel filtration chromatography. Fractions 37-41 were collected as pure 6xHis-JOX2 protein. (**C**) CBB-stained SDS-PAGE analysis of JOX1 after gel filtration chromatography. Fraction XXX were collected as pure 6xHis-JOX1 protein. (**D**) CBB-stained SDS-PAGE analysis of

JOX3 after gel filtration chromatography. Fraction 37-41 were collected as pure 6xHis-JOX3 protein. (**E**) CBB-stained SDS-PAGE analysis of JOX4 after gel filtration chromatography. Fraction 37-41 were collected as pure 6xHis-JOX4 protein.

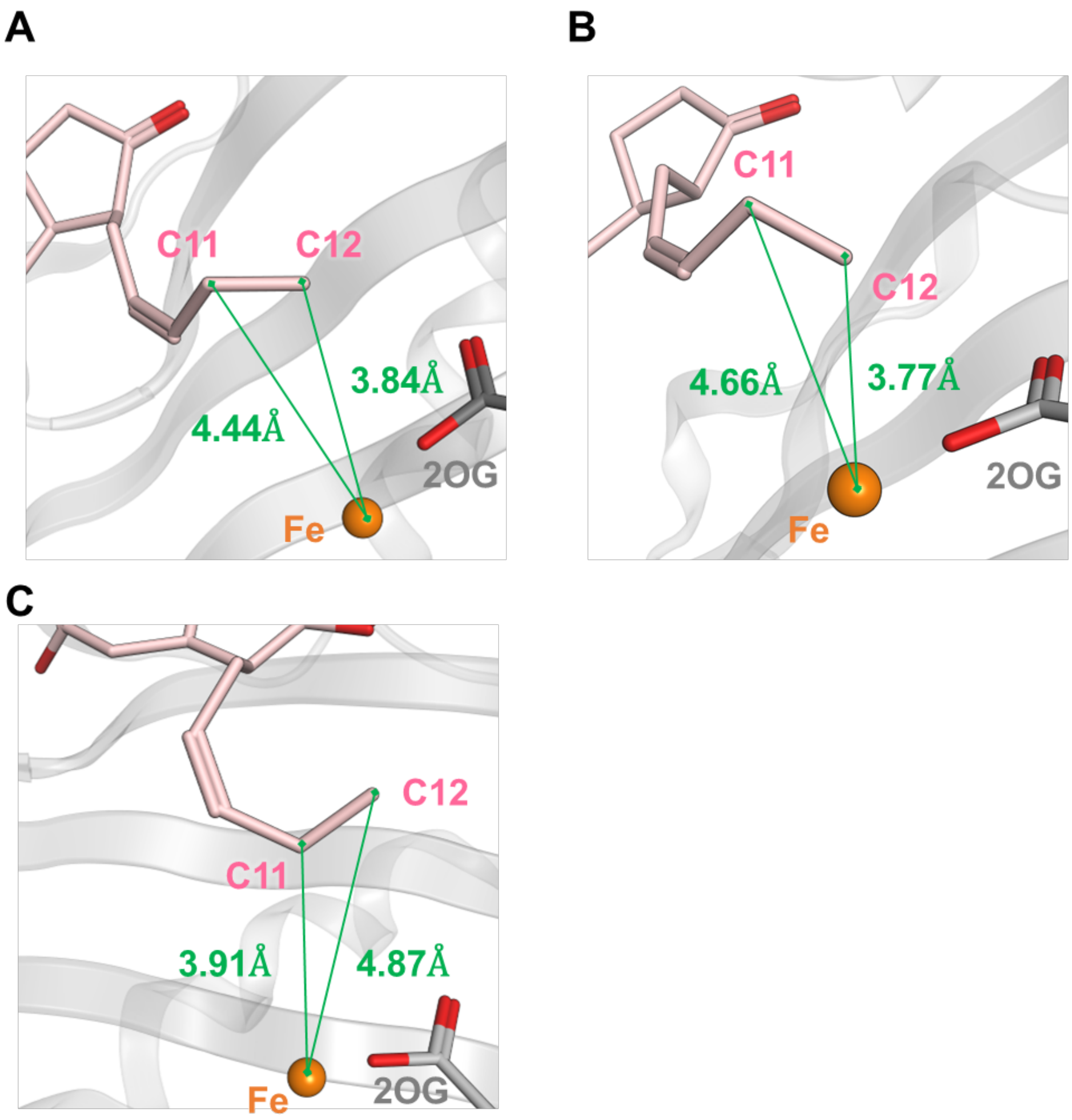


**Fig. S8** ***In silico*** **docking models of (A) JOX1, (B) JOX3, (C) JOX4**

Distances between the iron ion and the C12 and C11 carbon atoms of (*3R*,*7S*)-JA (pink). (A) In the JOX1, they are 3.84 Å for C12 and 4.44 Å for C11. (B) In the JOX3, they are 3.77 Å for C12 and 4.66 Å for C11. (C) In the JOX4, they are 3.91 Å for C12 and 4.87 Å for C12. JOXs and 2OG are represented in gray. Iron ion is shown as orange sphere. Distances are marked in green.

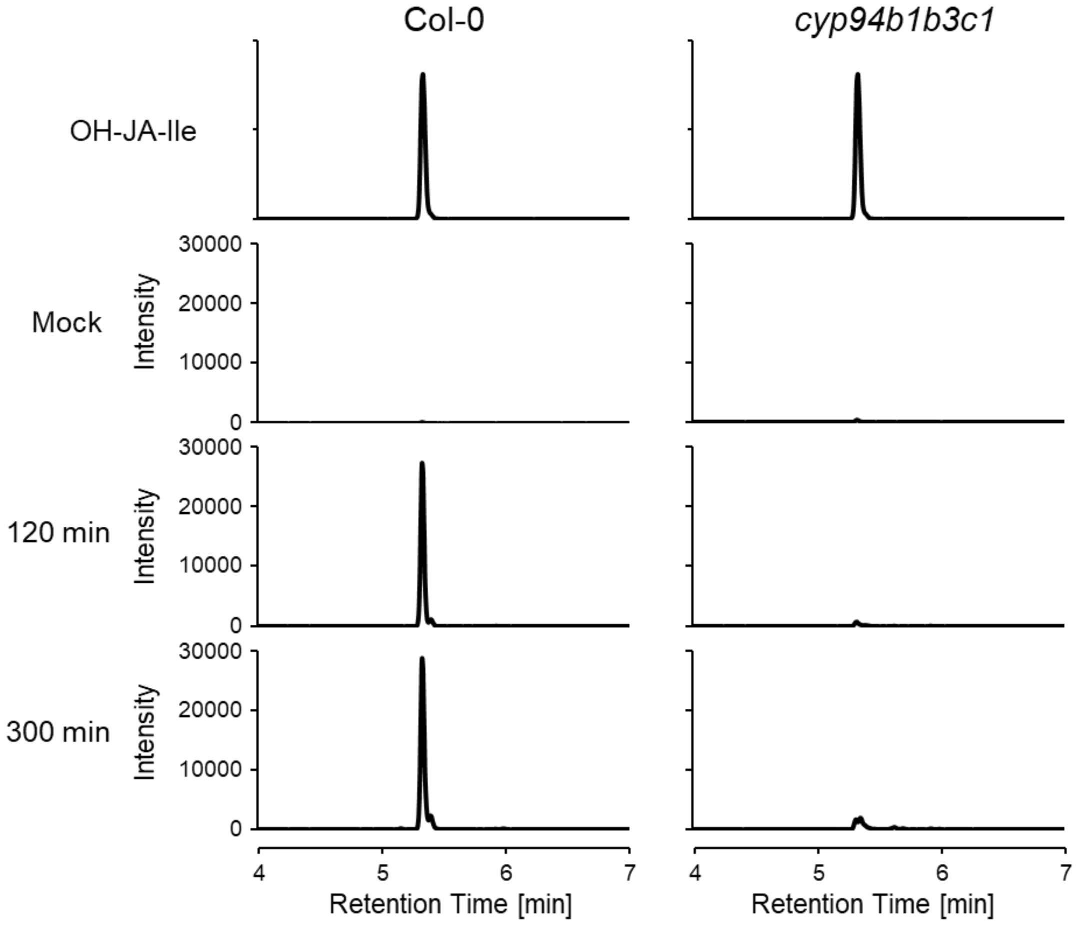


**Fig. S9 MS/MS (MRM) signal corresponding to 12- or 11-OH-JA-Ile in *cyp94b1cyp94b3cyp94c1* triple mutant**

UPLC-MS/MS chromatogram (*m*/*z* 338 > 130) of a mixture of 12- or 11-OH-JA-Ile. ZORBAX RRHD Eclipse Plus C18 1.8 μm (Φ2.1 × 50 mm; Agilent Technologies) was used at 23 °C and a flow rate of 0.350 mL/min. The elution was performed using water-$CH_3CN$ (95:5, v/v, isocratic), both containing 0.1% formic acid (v/v).

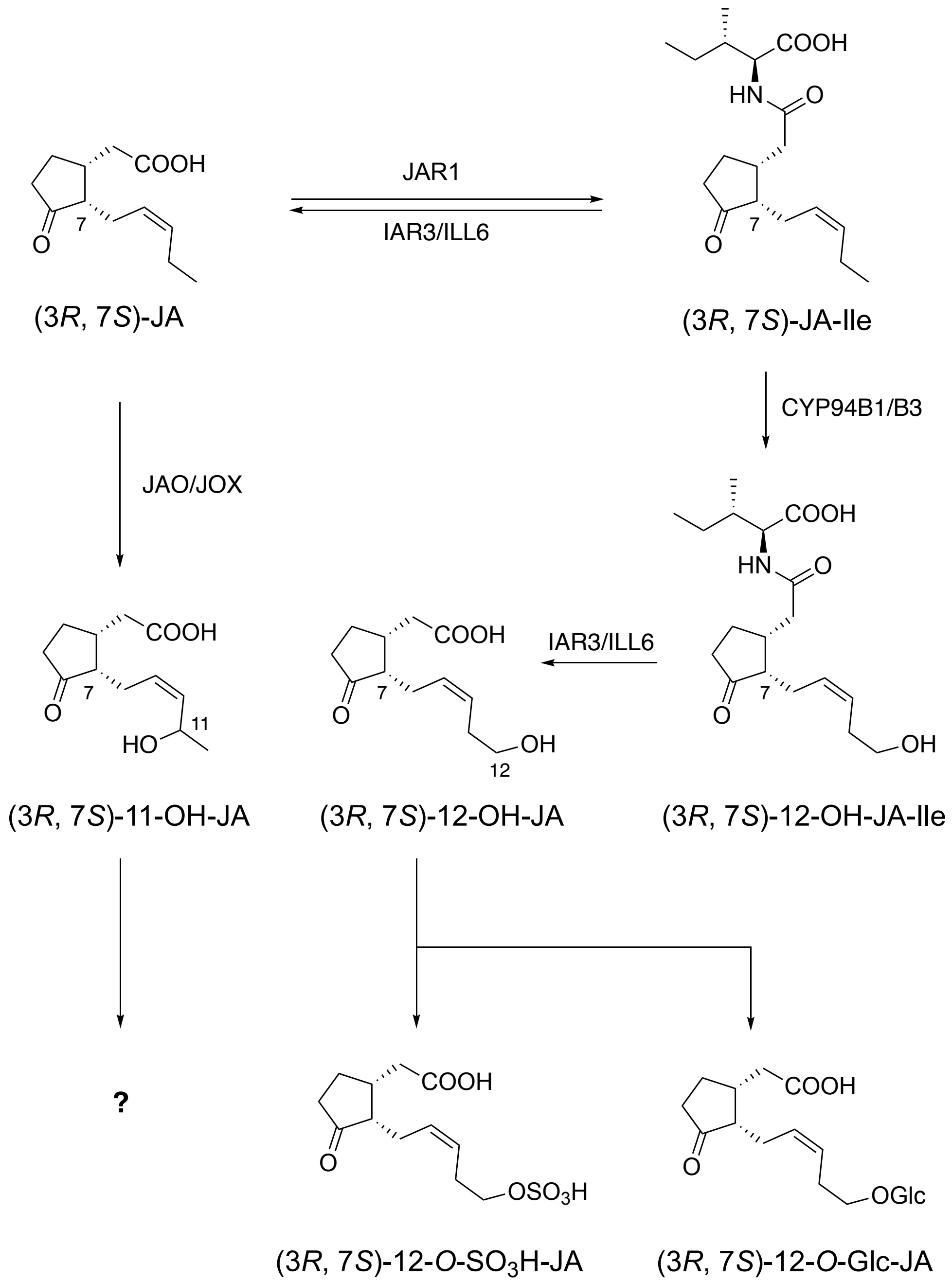


**Fig. S10** Current understanding of JA catabolism in *A. thaliana*. Further conversion of (3*R*, 7*S*)-11-OH-JA remains unknown.

### Supplementary Materials and Methods

### Chemical Syntheses

#### General materials and methods

All chemical reagents and solvents were obtained from commercial suppliers (Kanto Chemical Co. Ltd., Wako Pure Chemical Industries Co. Ltd., Nacalai Tesque Co. Ltd., Tokyo Chemical Industry Co. Ltd., Sigma-Aldrich Co. LLC., GE Healthcare) and used without further purification. All anhydrous solvents were either dried by standard techniques and freshly distilled before use or purchased in anhydrous form and used as supplied. Reversed-phase high–performance liquid chromatography (HPLC) was carried out on a PU–4180 plus pump equipped with UV–4075 and MD–4010 detectors (JASCO, Tokyo, Japan). $^{1}H$ and $^{13}C$ NMR spectra were recorded on a JNM–ECS–400 spectrometer (JEOL, Tokyo, Japan). HMBC/HSQC and NOESY experiments were carried out on a Bruker AVANCE NEO700 spectrometer equipped with cryoprobe (Bruker BioSpin Inc., Billerica, MA, US). Chemical shifts are denoted in δ (ppm) relative to TMS or residual solvent peaks as internal standard (TMS, $^{1}H$ δ 0.00; $CDCl_3$, $^{13}C$ δ 77.0; $CD_3OD$, $^{1}H$ δ 3.31, $^{13}C$ δ 49.0; pyridine-*d5*, $^{1}H$ δ 8.74, $^{13}C$ δ 150.4). Fourier transforms infrared (FT/IR) spectra were recorded on an FT/IR–4100 (JASCO, Tokyo, Japan). High–resolution (HR) electrospray ionization (ESI)–mass spectrometry (MS) analyses were conducted using a microTOF II (Bruker Daltonics Inc., Billerica, MA, US). Optical rotations were measured using a JASCO P–2200 polarimeter (JASCO, Tokyo, Japan). Flash chromatography was performed on an Isolera system (Biotage Ltd., North Carolina, US). TLC analyses were performed on Silica gel F254 (0.25 mm or 0.5 mm, MERCK, Germany) or RP–18F254S (0.25 mm, MERCK). All reactions were carried out under air unless stated otherwise.

**Synthesis of (*3R*, *7S*)-11-OH-JA and (*3R*, *7S*)-11-OH-JA-Ile**

**Synthesis of alcohol 6**

To a solution of $BH_3$·THF (63 mL, 0.91 M in THF, 57.3 mmol) in THF (100 mL) was added cyclohexene (11.8 mL, 116 mmol) at 0 °C under an argon atmosphere. The solution was stirred at 0 °C for 2 hours, and the supernatant was removed. The mixture was added to a solution of **1** (2.24 g, 14.7 mmol) which was prepared by previous method (27) in THF (50 mL). After stirring at 0 °C for 1 h, the reaction mixture was added $NaBO_3 \cdot 4H_2O$ (29.4 g, 191 mmol) and $H_2O$ (150 mL) . After stirring at room temperature for 3 h, the reaction mixture was quenched with saturated aqueous $NH_4Cl$ and extracted with EtOAc. The organic layer was washed with saturated aqueous NaCl, dried over $Na_2SO_4$, and concentrated under reduced pressure. The residue was purified by silica gel medium-pressure chromatography (*n*-hexane/EtOAc = 88/12 to 0/100) to give **6** (1.33 g, 60%) as a colorless oil. $^1$H NMR (400 MHz, $CDCl_3$) $\delta_H$ 6.04 (d, $J$ = 5.5 Hz, 1H), 5.81 (dt, $J$ = 5.5, 2.3 Hz, 1H), 5.66-5.52 (m, 1H), 3.75 (dt, $J$ = 10.6, 6.4 Hz, 1H), 3.70 (dt, $J$ = 10.6, 6.4 Hz, 1H), 2.77 (brt, $J$ = 6.4 Hz, 1H), 2.55 (dt, $J$ = 14.2, 7.8 Hz, 1H), 2.03 (s, 3H), 1.78 (q, $J$ = 6.4 Hz, 1H), 1.65 (q, $J$ = 6.4 Hz, 1H), 1.47 (dt, $J$ = 14.2, 4.6 Hz, 1H); $^{13}$C NMR (100 MHz, $CDCl_3$) $\delta_C$: 170.9, 140.6, 129.2, 79.7, 61.4, 40.95, 38.9, 36.4, 21.3; IR (neat) cm$^{-1}$: 3423, 2933, 2878, 1734, 1716, 1244; HRMS (ESI, positive) m/z $[M+Na]^+$ Calcd. for $C_9H_{14}NaO_3$: 193.0835, Found : 193.0834.

**Synthesis of alcohol 7**

To a solution of **6** (1.33 g, 7.79 mmol) in DMF (60 mL) was added imidazole (1.60 g, 23.5 mmol) and TBDPSCl (3.0 mL, 11.6 mmol) under an argon atmosphere. After being stirred at room temperature (rt) for 23 h, the reaction mixture was added MeOH and $H_2O$ and stirred 30 min. The solution was extracted with *n*-hexane, then organic layer was washed with brine, dried over $Na_2SO_4$, and concentrated under reduced pressure to afford a TBDPS-protected intermediate, which was used for the following reaction without further purification.

To a solution of above intermediate in MeOH (10 mL) was added NaOH (3 M in $H_2O$, 10 mL, 30 mmol). After being stirred at room temperature for 15 h, the reaction mixture was extracted with ether. The organic layer was washed with brine, dried over $Na_2SO_4$, and concentrated under reduced pressure. The residue was purified by silica gel medium-pressure chromatography (*n*-hexane/EtOAc = 93/7 to 40/60) to give **7** (1.87 g, 66% in 2 steps) as a yellow oil: $[\alpha]_D^{28}$ = -2 (*c* 1.02, $CHCl_3$); $^1H$ NMR (400 MHz, $CD_3OD$) $\delta_H$: 7.67 (dt, *J* = 7.8, 1.4 Hz, 4H), 7.38-7.45 (m, 6H), 5.81 (dt, *J* = 5.5, 1.8 Hz, 1H), 5.71 (dt, *J* = 5.5, 1.8 Hz, 1H), 4.72 (ddt, *J* = 7.8, 5.5, 1.8 Hz, 1H), 3.75 (dt, *J* = 10.1, 6.4 Hz, 1H), 3.72 (dt, *J* = 10.1, 6.4 Hz, 1H), 2.71 (*J* = 6.0, 1.8 Hz, 1H), 2.39 (dt, *J* = 13.3, 7.8 Hz, 1H), 1.79 (dt, *J* = 19.7, 6.4 Hz, 1H), 1.58 (dt, *J* = 19.7, 7.8 Hz, 1H), 1.19 (dt, *J* = 13.3, 6.0 Hz, 1H), 1.04 (s, 9H); $^{13}C$ NMR (100 MHz, $CD_3OD$) $\delta_C$: 138.7, 136.7 (4C), 135.0 (2C), 134.3, 130.9 (2C), 128.8(4C), 77.7, 63.8, 42.4, 41.1, 40.6, 27.4 (3C), 20.0; IR (neat) $cm^{-1}$: 3335, 2930, 2858, 1428, 1111, 705; HRMS (ESI, positive) m/z $[M+Na]^+$ Calcd. for $C_{23}H_{30}NaO_2Si$: 389.1913, Found : 389.1900.

**Synthesis of dimethylamide 8**

To a solution of **7** (1.87 g, 5.10 mmol) in xylene (18 mL) was added $MeC(OMe)_2NMe_2$ (3.8 mL, 26.0 mmol) under an argon atmosphere. The mixture was stirred at 150 °C, and MeOH was removed in a Dean Stark apparatus with MS4A. After

16 h, the reaction mixture was concentrated under reduced pressure. The residue was purified by silica gel medium-pressure chromatography (*n*-hexane/EtOAc = 93/7 to 40/60) to give **8** (1.24 g, 56%) as a brown oil: $[\alpha]_D^{28}$ = -62 (*c* 1.19, $CHCl_3$); $^1H$ NMR (400 MHz, $CDCl_3$) $\delta_H$: 7.65-7.68 (m, 4H), 7.36-7.44 (m, 6H), 5.79-5.82 (m, 1H), 5.68-5.71 (m, 1H), 3.64-3.74 (m, 2H), 3.04-3.11 (m, 1H), 2.95 (s, 3H), 2.94 (s, 3H), 2.36-2.46 (m, 1H), 2.29-2.35 (m, 1H), 2.28 (dd, *J* = 14.7, 5.0 Hz, 1H), 2.07(dd, *J* = 14.7, 10.1 Hz, 1H), 1.92 (ddq, *J* = 16.0, 8.2, 2.3 Hz, 1H), 1.71-1.80 (m, 1H), 1.51 (dddd, *J* = 12.9, 9.7, 7.0, 5.0 Hz, 1H), 1.04 (s, 9H); $^{13}C$ NMR (100 MHz, $CDCl_3$) $\delta_C$: 172.5, 135.6 (4C), 135.5, 134.0 (2C), 130.2, 129.6 (2C), 127.6 (4C), 63.4, 43.6, 37.8, 37.4, 37.1, 35.4, 33.6, 33.4, 26.9 (3C), 19.2; IR (neat) $cm^{-1}$: 3049, 2931, 1651, 1394, 1111, 705; HRMS (ESI, positive) m/z $[M+Na]^+$ Calcd. for $C_{27}H_{37}NNaO_2Si$: 458.2486, Found : 458.2502.

**Synthesis of iodolactone 9**

To a solution of **8** (1.24 g, 2.86 mmol) in THF/$H_2O$ (6.4 mL, 1:1, v/v) was added $KH_2PO_4$ (511 mg, 3.75 mmol), $Na_2HPO_4 \cdot 4H_2O$ (138 mg, 773 μmol) and $I_2$ (1.45 g, 5.73 mmol). The solution was stirred room temperature for 26 h. The reaction mixture was quenched with saturated aqueous $Na_2S_2O_3$ and extracted with EtOAc. The organic layer was washed with brine, dried over $Na_2SO_4$, and concentrated under reduced pressure. The residue was purified by silica gel medium-pressure chromatography (*n*-hexane/EtOAc = 95/5 to 60/40) to give dimethylamide **9** (1.29 g, 84%) as a yellow oil: $[\alpha]_D^{28}$ = +5 (*c* 1.15, $CHCl_3$); $^1H$ NMR (400 MHz, $CDCl_3$) $\delta_H$: 7.65-7.68 (m, 4H), 7.38-7.47 (m, 6H), 5.24 (d, *J* = 6.4 Hz, 1H), 4.43 (d, *J* = 5.0 Hz, 1H), 3.68 (dt, *J* = 10.1, 6.4 Hz, 1H), 3.64 (dt, *J* = 10.1, 6.4 Hz, 1H), 2.97-3.04 (m, 1H), 2.85-2.95 (m, 1H), 2.47 (dd, *J* = 18.8, 10.1 Hz, 1H), 2.38 (dd, *J* = 18.8, 4.1 Hz, 1H), 2.08 (dd, *J* = 14.7, 5.5 Hz, 1H), 1.58-1.74 (m, 3H), 1.06

(s, 9H); $^{13}$C NMR (100 MHz, $CDCl_3$) $\delta_C$: 176.4, 135.6 (4C), 133.5, 133.4, 129.8 (2C), 127.8 (4C), 92.7, 62.4, 40.1, 38.9, 37.1, 32.6, 28.9, 28.0, 26.9 (3C), 19.1; IR (neat) $cm^{-1}$: 2930, 2857, 1784, 1427, 1111, 704; HRMS (ESI, positive) m/z $[M+Na]^+$ Calcd. for $C_{25}H_{31}INaO_3Si$: 557.0979, Found : 557.0967.

**Synthesis of diol 11**

To a solution of **9** (3.08 g, 5.75 mmol) in benzene (35 mL) was added *n*-$Bu_3SnH$ (4.65 mL, 17.3 mmol), AIBN (188 mg, 1.14 mmol). The mixture was stirred at 90 °C for 40 min, then cooled to rt. The reaction mixture was diluted with $CH_2Cl_2$ and quenched with 1M NaOHaq. , then was extracted with $CH_2Cl_2$. The organic layer was washed with saturated aqueous $NH_4Cl$, brine, dried over $Na_2SO_4$, and concentrated under reduced pressure to afford **10**, which was used for the following reaction without further purification.

To a solution of crude **10** in THF (60 mL) was added $LiAlH_4$ (658.2 mg, 17.3 mmol) at -20 ºC. After being stirred at 0 ºC for 40 min, the reaction mixture was re-cooled to -20 ºC and then a homogeneous mixture of $SiO_2$ (68 g) and $H_2O$ (20 mL) was added. The mixture was stirred for 40 min and the solids were removed by filtration and washed thoroughly with EtOAc and concentrated under reduced pressure. The residue was purified by Silica gel medium-pressure chromatography (*n*-hexane/EtOAc = 88/12 to 0/100) to give diol **11** (2.20 g, 93%) as a colorless oil: $[\alpha]_D^{22} = +11$ (*c* 1.56, $CHCl_3$); $^1$H NMR (400 MHz, $CDCl_3$) $\delta_H$: 7.68-7.65 (m, 4H), 7.45-7.35 (m, 6H), 4.27-4.22 (m, 1H), 3.81 (dt, *J* = 10.1, 5.0 Hz, 1H), 3.69 (ddd, *J* = 10.1, 7.3, 5.0 Hz, 1H), 3.65-3.57 (m, 2H), 2.10-2.00 (m, 1H), 1.91-1.81 (m, 2H), 1.79-1.67 (m, 2H), 1.66-1.53 (m, 3H), 1.52-1.36 (m, 2H), 1.04 (s, 9H); $^{13}$C NMR (100 MHz, $CDCl_3$) $\delta_C$: 135.6 (4C), 134.0 (2C), 129.5

(2C), 127.6 (4C), 77.2, 74.6, 63.1, 63.0, 46.3, 36.8, 34.1, 32.7, 27.3, 26.9 (3C), 19.2; IR (neat) $cm^{-1}$: 3335, 2932, 2858, 1428, 1111, 823, 739, 703; HRMS (ESI, positive) m/z $[M+Na]^+$ Calcd. for $C_{25}H_{36}NaO_3Si$: 435.2331, Found : 435.2323.

**Synthesis of 12**

To a solution of **11** (2.20 g, 5.33 mmol) in $CH_2Cl_2$ (30 mL) was added DMAP (65.2 mg, 0.534 mmol), Trt-Cl (1.56 g, 5.60 mmol) and $Et_3N$ (1.48 mL, 10.7 mmol). The solution was stirred at rt for 11 h. The reaction mixture was quenched with saturated aqueous $NH_4Cl$ and extracted with $CH_2Cl_2$. The organic layer was washed with brine, dried over $Na_2SO_4$, and concentrated under reduced pressure. The residue was purified by silica gel medium-pressure chromatography (*n*-hexane/EtOAc = 98/2 to 80/20) to give **12** (2.59 g, 74%) as a pale yellow oil: $[\alpha]_D^{26}$ = +23 (*c* 0.95, $CHCl_3$); $^1H$ NMR (400 MHz, $CDCl_3$) $\delta_H$: 7.67-7.63 (m, 4H), 7.45-7.33 (m, 11H), 7.32-7.19 (m, 10H), 4.10 (m, 1H), 3.65 (ddd, *J* = 10.1, 7.8, 5.0 Hz, 1H), 3.55 (dt, *J* = 10.1, 7.3 Hz, 1H), 3.34 (dt, *J* = 9.2, 5.0 Hz, 1H), 3.04 (ddd, *J* = 9.2, 7.8, 5.0 Hz, 1H), 2.29 (d, *J* = 3.2 Hz, 1H), 1.99-1.89 (m, 1H), 1.88-1.53 (m, 8H), 1.49-1.37 (m, 2H), 1.03 (s, 9H); $^{13}C$ NMR (100 MHz, $CDCl_3$) $\delta_C$: 144.0 (3C),135.6 (4C), 134.1 (2C), 129.5 (2C), 129.4, 128.7 (2C), 128.6 (2C), 129.9 (2C), 127.8 (2C), 127.7 (2C), 127.6 (2C), 127.5 (2C), 127.1, 126.9 (3C), 87.3, 74.3, 63.2, 46.4, 36.8, 34.2, 32.9, 28.9, 26.9 (3C), 25.3, 19.2; IR (neat) $cm^{-1}$: 3474, 2933, 1959, 1890, 1823, 1735, 1596, 1490, 701; HRMS (ESI, positive) m/z $[M+Na]^+$ Calcd. for $C_{44}H_{50}NaO_3Si$: 677.3421, Found : 677.3427.

**Synthesis of 13**

To a solution of **12** (2.59 g, 3.95 mmol) in DMF (20 mL) was added imidazole

(802 mg, 11.8 mmol) and TBDPSCl (2.00 mL, 7.79 mmol). The solution was stirred at rt for 19 h. The reaction mixture was quenched with saturated aqueous $NaHCO_3$ and extracted with hexane. The organic layer was washed with brine, dried over $Na_2SO_4$, and concentrated under reduced pressure. The residue was purified by silica gel medium-pressure chromatography (*n*-hexane/EtOAc = 100/0 to 94/6) to give **13** (3.04 g, 86%) as a pale yellow oil: $[\alpha]_D^{26}$ = +6.4 (*c* 1.89, $CHCl_3$); $^1H$ NMR (400 MHz, $CDCl_3$) $\delta_H$: 7.65-7.61 (m, 4H), 7.60-7.54 (m, 4H), 7.42-7.27 (m, 19H), 7.25-7.15 (m, 8H), 4.00 (m, 1H), 3.58 (ddd, *J* = 9.6, 7.3, 4.8 Hz, 1H), 3.50 (dt, *J* = 10.1, 6.9 Hz, 1H), 3.11-3.04 (m, 1H), 3.03-2.96 (m, 1H), 1.99-1.87 (m, 1H), 1.86-1.74 (m, 1H), 1.73-1.60 (m, 3H), 1.53-1.30 (m, 3H), 1.01 (s, 9H), 1.00 (s, 9H); $^{13}C$ NMR (100 MHz, $CDCl_3$) $\delta_C$: 144.6 (3C), 135.9 (4C), 135.5 (4C), 134.8, 134.2, 134.1 (2C), 129.5 (2C), 129.4 (2C), 128.7 (6C), 127.6 (6C), 127.5 (4C), 127.4 (2C), 127.3 (2C), 126.7 (3C), 86.3, 77.2, 63.4, 63.3, 44.6, 35.9, 34.5, 32.9, 28.2, 27.1 (3C), 26.9 (3C), 25.3, 19.2 (2C); IR (neat) $cm^{-1}$: 2856, 1959, 1890, 1824, 1590, 1490, 1110; HRMS (ESI, positive) m/z $[M+Na]^+$ Calcd. for $C_{60}H_{68}NaO_3Si_2$: 915.4599, Found : 915.4617.

**Synthesis of 14**

To a solution of **13** (3.04 g, 3.40 mmol) in $CH_2Cl_2$ (10 mL) and MeOH (10 mL) was added *p*-TsOH･$H_2O$ (194.1mg, 1.02 mmol). The solution was stirred at rt for 15 min. The reaction mixture was quenched with saturated aqueous $NaHCO_3$ and extracted with $CH_2Cl_2$. The organic layer was washed with brine, dried over $Na_2SO_4$, and concentrated under reduced pressure. The residue was purified by silica gel medium-pressure chromatography (*n*-hexane/EtOAc = 98/2 to 80/20) to give **14** (1.70 g, 77%) as a colorless oil: $[\alpha]_D^{25}$ = +7.9 (*c* 0.93, $CHCl_3$); $^1H$ NMR (400 MHz, $CDCl_3$) $\delta_H$: 7.68-7.63 (m, 8H),

7.44-7.33 (m, 12H), 4.14 (m, 1H), 3.68 (ddd, $J$ = 10.1, 7.3, 5.0 Hz, 1H), 3.63-3.49 (m, 3H), 1.96-1.85 (m, 1H), 1.83-1.69 (m, 3H), 1.65-1.30 (m, 6H), 1.06 (s, 9H), 1.04 (s, 9H); $^{13}$C NMR (100 MHz, $CDCl_3$) $\delta_C$: 136.0 (2C), 135.9 (2C), 135.6 (4C), 134.4, 134.1, 134.0, 133.8, 129.7, 129.6, 129.5 (2C), 127.6 (6C), 127.5 (2C), 77.2, 76.9, 63.1, 62.6, 44.6, 36.0, 34.4, 32.3, 28.0, 27.1 (3C), 26.9 (3C), 19.2 (2C); IR (neat) cm$^{-1}$: 3397, 3071, 2932, 1739, 1589, 1472, 1427, 1390, 1241, 1108; HRMS (ESI, positive) m/z [M+Na]$^+$ Calcd. for $C_{41}H_{54}NaO_3Si_2$: 673.3504, Found : 673.3483.

**Synthesis of 2**

To a solution of DMSO (0.56 mL, 7.88 mmol) in $CH_2Cl_2$ (5.0 mL) was added oxalyl chloride (0.40 mL, 4.66 mmol) at -78 °C under an argon atmosphere. After the reaction mixture was stirred at -78 °C for 40 min, a solution of **14** (463 mg, 0.711 mmol) in $CH_2Cl_2$ (14 mL) was slowly added. After the reaction mixture was stirred at -65 °C for 2 h, $Et_3N$ (0.99 mL, 7.14 mmol) was slowly added. The mixture was gradually warmed to room temperature for 1 h with stirring. The reaction mixture was quenched with saturated aqueous $NH_4Cl$. The mixture was extracted with *n*-hexane. The organic layer was washed with saturated aqueous NaCl, dried over $Na_2SO_4$, and filtered. The reaction mixture was concentrated under reduced pressure. The residue was purified by silica gel medium-pressure chromatography (*n*-hexane/EtOAc = 95/5) to give aldehyde intermediate (410 mg, 89%) as a pale yellow oil and was used for the next reaction immediately.

To a solution of aldehyde intermediate in anhydrous MeOH (9.0 mL) was added $K_2CO_3$ (171 mg, 1.24 mmol) and Ohira-Bestmann reagent (290 μL, 1.93 mmol). The mixture was stirred for 6 h. The reaction mixture was quenched with saturated aqueous

$NaHCO_3$ and extracted with *n*-hexane. The organic layer was washed with brine, dried over $Na_2SO_4$, and concentrated under reduced pressure. The residue was purified by silica gel medium-pressure chromatography (*n*-hexane/EtOAc = 98/2) to give **2** (313 mg, 77%) as a pale yellow oil: $[\alpha]_D^{29}$ = +11.0 (*c* 0.97, $CHCl_3$); $^1H$ NMR (400 MHz, $CDCl_3$) $\delta_H$: 7.71-7.61 (m, 8H), 7.44-7.32 (m, 12H), 4.19 (q, *J* = 5.0 Hz, 1H), 3.77-3.60 (m, 2H), 2.46 (ddd, *J* = 16.9, 5.5, 2.8 Hz, 1H), 2.24 (ddd, *J* = 16.9, 8.7, 2.3 Hz, 1H), 2.14-2.03 (m, 1H), 2.01-1.92 (m, 2H), 1.85 (t, *J* = 2.8 Hz, 1H), 1.68-1.36 (m, 5H), 1.06 (s, 9H), 1.04 (s, 9H); $^{13}C$ NMR (100 MHz, $CDCl_3$) $\delta_C$: 136.0 (2C), 135.9 (2C), 135.6 (4C), 134.6, 134.2, 134.1, 133.8, 129.5 (4C), 127.6 (8C), 85.1, 76.1, 68.2, 63.1, 62.6, 47.5, 35.8, 34.3, 32.8, 29.7, 28.1, 27.0 (3C), 26.9 (3C), 26.8, 19.3, 19.2; IR (neat) $cm^{-1}$: 3309, 3071, 2998, 2857, 2117, 1959, 1889, 1824, 1732, 1471, 1427; HRMS (ESI, positive) m/z $[M+Na]^+$ Calcd. for $C_{42}H_{52}NaO_2Si_2$: 667.3398, Found: 673.3405.

**Synthesis of 3**

To a solution of **2** (313 mg, 0.485 mmol) in THF (10 mL) was added *n*BuLi (0.76 mL, 1.22 mmol, 1.6 M in hexane) at -78 °C under an argon atmosphere. The reaction mixture was warmed to 0 ºC and stirred for 1.5 h. The reaction mixture was re-cooled to -78 ºC. After the reaction mixture was stirred at -78 °C for 30 min, a solution of acetaldehyde (141 μL, 2.52 mmol) in THF (1.27 mL) was slowly added. The reaction mixture was warmed to room temperature and stirred for 6 h. The reaction mixture was quenched with saturated aqueous $NH_4Cl$. The mixture was extracted with $CH_2Cl_2$. The organic layer was washed with saturated aqueous NaCl, dried over $Na_2SO_4$, and filtered. The residue was purified by silica gel medium-pressure chromatography (*n*-hexane/EtOAc = 98/2 to 80/20) to give alkyne intermediate (231 mg, 69%) as a yellow oil.

To a solution of alkyne intermediate (47.3 mg, 68.6 mmol) in anhydrous MeOH (2.0 mL) was added quinoline (8.2 μL, 69.2 mmol) and Pd/$BaSO_4$ (4.3 mg, 9 wt%) under an argon atmosphere. The reaction vessel was evacuated and recharged with $H_2$. After being stirred at room temperature for 10 min under $H_2$ atmosphere, the reaction mixture was filtered through a pad of Celite with EtOAc. The filtrate was concentrated under reduced pressure. The residue was purified by silica gel medium-pressure chromatography (*n*-hexane/EtOAc = 98/2 to 80/20) to give **12** (42.4 mg, 89%) as a colorless oil. **3** was obtained as the stereoisomer mixture at C-11 position: $[\alpha]_D^{22}$ = +20.2 (*c* 0.75, $CHCl_3$); $^1$H NMR (400 MHz, $CDCl_3$) $\delta_H$: 7.60-7.69 (m, 8H), 7.32-7.46 (m, 12H), 5.30-5.44 (m, 2H), 4.65 (quin, *J* = 6.2 Hz, 0.5H), 4.57 (quin, *J* = 6.4 Hz, 0.5H), 4.15 (q, *J* = 5.0 Hz, 1H), 3.68 (dtd, *J* = 10.1, 5.0, 2.3 1H), 3.60 (dt, *J* = 10.1, 6.9 Hz, 1H), 2.08-2.31 (m, 2H), 1.90-2.02 (m, 1H), 1.75-1.88 (m, 1H), 1.63-1.71 (m, 1H), 1.30-1.63 (m, 5H), 1.22 (d, *J* = 6.4 Hz, 1.5H), 1.22 (d, *J* = 6.2 Hz, 1.5H), 1.07 (s, 4.5H), 1.06 (s, 4.5H), 1.04 (s, 9H); $^{13}$C NMR (100 MHz, $CDCl_3$) $\delta_C$: 135.9, 135.5, 134.6, 134.5, 134.2, 134.1 (2C), 133.9 (2C), 133.5, 133.3, 131.5, 131.0, 129.5, 127.6, 127.5, 127.4, 77.2, 76.6, 63.9, 63.8, 63.1 (2C), 48.5, 48.4, 35.8, 35.7, 34.5, 34.2, 32.9, 28.1, 27.0 (3C), 26.8 (3C), 23.3, 23.2, 23.0 (2C), 19.2 (2C); IR (neat) cm$^{-1}$: 3368, 3072, 2931, 2858, 1471, 1428, 1390, 1110, 822, 740; HRMS (ESI, positive) m/z $[M+Na]^+$ Calcd. for $C_{44}H_{58}NaO_3Si_2$: 713.3817, Found : 713.3825.

**Synthesis of 4**

To a solution of **3** (202 mg, 0.292 mmol) in $CH_2Cl_2$ (10 mL) was added DHP (40 μL, 0.44 mmol) and *p*-TsOH•$H_2O$ (0.7 mg, 5 μmol) at 0 ºC. The solution was stirred at

0 ºC for 2 h. The reaction mixture was quenched with saturated aqueous $NaHCO_3$ and extracted with $CH_2Cl_2$. The organic layer was washed with brine, dried over $Na_2SO_4$, and concentrated under reduced pressure. The residue was purified by silica gel medium-pressure chromatography (*n*-hexane/EtOAc = 98/2 to 90/10) to give **4** (221 mg, 98%) as a pale yellow oil. **4** was obtained as the stereoisomer mixture at C-11 and THP position: $^1$H NMR (400 MHz, $CDCl_3$) $\delta_H$: 7.59-7.72 (m, 8H), 7.30-7.46 (m, 12H), 5.12-5.61 (m, 2H), 4.46-4.77 (m, 2H), 4.09-4.28 (m, 1H), 3.78-3.94 (m, 1H), 3.56-3.74(m, 2H), 3.35-3.51 (m, 1H), 2.14-2.35 (m, 2H), 1.936-2.05 (m, 1H), 1.75-1.91 (m, 2H), 1.36-1.74 (m, 11H), 1.25 (d, *J* = 6.4 Hz, 0.75H), 1.22 (d, *J* = 6.9 Hz, 0.75H), 1.19 (d, *J* = 6.0 Hz, 0.75H), 1.16 (d, *J* = 6.4 Hz, 0.75H), 1.07 (s, 9H), 1.04 (s, 9H); $^{13}$C NMR (100 MHz, $CDCl_3$) $\delta_C$: 136.1, 135.7, 133.4, 132.9, 132.4, 132.1, 131.5, 131.0, 130.5, 130.1, 129.6, 127.7, 96.7, 96.5, 96.3, 96.0, 76.7, 68.3, 68.2, 66.9, 66.7, 63.4, 63.3, 63.2,62.6, 62.5, 48.9, 48.8, 48.6, 36.1, 36.0, 35.8, 34.6(2C), 34.5, 33.1, 31.3, 31.2, 31.1, 28.3, 27.2, 27.0, 25.7, 25.6, 25.6, 23.4, 23.3, 23.1(2C), 21.7(2C), 20.5(2C), 19.9, 19.8, 19.4, 19.3; IR (neat) $cm^{-1}$: 3072, 2932, 2859, 1428, 1389, 1111, 822, 740; HRMS (ESI, positive) m/z $[M+Na]^+$ Calcd. for $C_{49}H_{66}NaO_4Si_2$: 797.4392, Found : 797.4379.

**Synthesis of 5**

To a solution of **4** (111 mg, 0.144 mmol) in anhydrous THF (10 mL) was added TBAF (0.72 mL, 0.72 mmol, 1.0 M in THF) under an argon atmosphere. The reaction mixture was stirred for 2 h at reflux condition. The reaction mixture was cooled to rt. The reaction mixture was quenched with saturated aqueous $NH_4Cl$. The mixture was extracted with EtOAc. The organic layer was washed with brine, dried over $Na_2SO_4$, and filtered. The residue was purified by silica gel medium-pressure chromatography (*n*-

hexane/EtOAc = 90/10 to 0/100) to give **5** (38.9 mg, 91%) as a pale yellow oil. **5** was obtained as the stereoisomer mixture at C-11 and THP position: $^{1}$H NMR (400 MHz, $CDCl_3$) $\delta_H$: 5.15-5.65 (m, 2H), 4.60-4.84 (m, 2H), 4.09-4.28 (m, 1H), 4.04-4.24 (m, 1H), 3.42-3.96(m, 4H), 2.45-2.61(m, 0.5H), 2.17-2.33(m, 1H), 1.47-2.12(m, 14.5H), 1.27 (d, *J* = 6.4 Hz, 1.5H), 1.22 (d, *J* = 6.0 Hz, 1.5H); $^{13}$C NMR (100 MHz, $CDCl_3$) $\delta_C$: 133.5, 132.8, 132.7, 132.3, 132.2, 131.2, 129.7, 129.6, 98.5, 97.6, 94.6, 93.6, 75.3, 75.2, 73.6(2C), 70.2, 69.4, 66.5, 65.6, 63.2(2C), 62.3, 62.2(2C), 61.8, 60.9, 48.7, 48.3, 48.2, 48.0, 36.3(2C), 35.9(2C), 35.5, 35.3, 35.0, 34.7, 33.7, 33.6, 33.4, 33.3, 31.3, 31.2, 30.8, 30.2, 29.2, 29.0, 28.9, 28.8, 25.7, 25.5(2C), 25.4, 23.8, 23.7, 23.6, 23.5, 21.8, 21.7, 21.4, 20.8, 20.1, 19.9, 19.1, 18.4; IR (neat) cm$^{-1}$: 3409, 2930, 1728, 1442, 1022, 899, 868, 809, 754; HRMS (ESI, positive) m/z $[M+Na]^+$ Calcd. for $C_{17}H_{30}NaO_4$: 321.2036, Found :321.2036.

**Determination of Stereochemistry of (3*R*,7*S*, 11*R*)-11-OH-JA and (3*R*,7*S*, 11*S*)-11-OH-JA by modified Mosher's method**

**Isolation of a single stereoisomer of 3**

Single stereoisomer of **3** was isolated from a mixture of (*3R, 7S, 11R/S*)-**3** by multiple silica gel medium-pressure chromatography. From a mixture of (*3R, 7S, 11R/S*)-**3** (39.2 mg), (*3R, 7S, 11R*)-**3** (14.2 mg) and (*3R, 7S, 11S*)-**3** (17.5 mg) were isolated.

**Synthesis of (*3R, 7S, 11S*)-3-(*S*)-MTPA ester**

To solution of (*3R, 7S, 11S*)-**3** (2.2 mg, 3.2 μmol) in DCM (2.0 mL) and pyridine (0.10 mL) was added (*R*)-MTPA-Cl (4.0 μL, 21 μmol). The solution was stirred at rt for 2 h. The reaction mixture was quenched with 1M aqueous HCl and extracted with $CHCl_3$.

The organic layer was washed with brine, dried over $Na_2SO_4$, and concentrated under reduced pressure. The residue was roughly purified by silica gel chromatography (*n*-hexane/EtOAc = 98/2 to 95/5) to give (*3R, 7S, 11S*)-**3**-(*S*)-MTPA ester (0.5 mg) as a pale yellow amorphous solid: $^1$H NMR (400 MHz, $CDCl_3$) $\delta_H$: 7.68-7.31 (m, 25H), 5.88 (dq, *J* = 9.2, 6.4 Hz, 1H), 5.55 (dt, *J* = 11.0, 6.9 Hz, 1H), 5.34 (dd, *J* = 11.0, 9.2 Hz, 1H), 4.19-4.14 (m, 1H), 3.71-3.55 (m, 2H), 3.51 (s, 3H), 2.44-2.31 (m, 2H), 2.30-2.20 (m, 1H), 2.05-1.95 (m, 2H), 1.85-1.75 (m, 1H), 1.72-1.63 (m, 1H), 1.61-1.47 (m, 3H), 1.26 (d, *J* = 6.4 Hz, 3H), 1.06 (s, 9H), 1.03 (s, 9H); HRMS (ESI, positive) m/z $[M+Na]^+$ Calcd. for $C_{54}H_{65}F_3NaO_5Si_2$: 929.4215, Found : 929.4242.

**Synthesis of (*3R, 7S, 11S*)-3-(*R*)-MTPA ester**

To solution of (*3R, 7S, 11S*)-**3** (2.3 mg, 3.3 μmol) in DCM (2.0 mL) and pyridine (0.10 mL) was added (*S*)-MTPA-Cl (4.0 μL, 21 μmol). The solution was stirred at rt for 1 h. The reaction mixture was quenched with 1M aqueous HCl and extracted with $CHCl_3$. The organic layer was washed with brine, dried over $Na_2SO_4$, and concentrated under reduced pressure to give crude (*3R, 7S, 11S*)-**3**-(*R*)-MTPA ester (1.2 mg) as a pale yellow amorphous solid: $^1$H NMR (400 MHz, $CDCl_3$) $\delta_H$: 7.68-7.31 (m, 25H), 5.85 (dq, *J* = 8.7, 6.4 Hz, 1H), 5.52 (dt, *J* = 11.0, 7.3 Hz, 1H), 5.24 (dd, *J* = 11.0, 9.2 Hz, 1H), 4.17 (q, *J* = 5.0 Hz, 1H), 3.71-3.58 (m, 2H), 3.53 (s, 3H), 2.45-2.35 (m, 2H), 2.28-2.17 (m, 1H), 2.05-1.95 (m, 2H), 1.85-1.75 (m, 1H), 1.72-1.64 (m, 1H), 1.61-1.47 (m, 3H), 1.33 (d, *J* = 6.0 Hz, 3H), 1.06 (s, 9H), 1.03 (s, 9H); HRMS (ESI, positive) m/z $[M+Na]^+$ Calcd. for $C_{54}H_{65}F_3NaO_5Si_2$: 929.4215, Found : 929.4233.

**Synthesis of (*3R,7S, 11S*)-11-OH-JA from (*3R,7S, 11S*)-3 (Figure S3B)**

**Synthesis of (*3R,7S, 11S*)-5**

To a solution of (*3R,7S, 11S*)-**3** (12 mg, 17 μmol) in $CH_2Cl_2$ (4.0 mL) was added DHP (6.0 μL, 67 μmol) and *p*-TsOH·$H_2O$ (0.16 mg, 1 μmol) at 0 ºC. The solution was stirred at 0 ºC for 1 h. The reaction mixture was quenched with saturated aqueous $NaHCO_3$ and extracted with $CH_2Cl_2$. The organic layer was washed with brine, dried over $Na_2SO_4$, and concentrated under reduced pressure. The residue was purified by silica gel chromatography (*n*-hexane/EtOAc = 95/5) to give (*3R,7S, 11S*)-**4** (12 mg, 91%) as a pale yellow oil. (*3R,7S, 11S*)-**4** was obtained as the stereoisomer mixture at THP position and directly used for the next reaction.

To a solution of (*3R,7S, 11S*)-**4** (12 mg, 16 μmol) in anhydrous THF (6.0 mL) was added TBAF (0.10 mL, 0.10 mmol, 1.0 M in THF) under an argon atmosphere. The reaction mixture was stirred for 16 h at rt. The reaction mixture was quenched with saturated aqueous $NH_4Cl$. The mixture was extracted with EtOAc. The organic layer was washed with brine, dried over $Na_2SO_4$, and filtered. The residue was purified by silica gel chromatography (*n*-hexane/EtOAc = 50/50 to 40/60) to give (*3R,7S, 11S*)-**5** (4.2 mg, 89%) as a yellow oil. (*3R,7S, 11S*)-**5** was obtained as the stereoisomer mixture at THP position and directly used for the synthesis of (*3R,7S, 11S*)-11-OH-JA.

**Synthesis of (*3R,7S, 11S*)-11-OH-JA**

To a solution of (*3R,7S, 11S*)-**5** (4.2 mg, 14 umol) in acetone (10 mL) was added Jones reagent (4.0 M solution, 90 uL, 0.36 mmol) at -20 ºC. After 1 h of stirring at -20 ºC, *i*-PrOH (1.4 mL) was added to quench the remaining reagent. After 1 h of stirring at -20 ºC, EtOAc/*n*-hexane (1/1, 10 mL) and $H_2O$ (15 mL) were added, and the aqueous layer was extracted with EtOAc. The combined organic layers were washed with brine, dried over $Na_2SO_4$, and concentrated under reduced pressure. The residue was purified by silica gel chromatography (*n*-hexane+0.1%AcOH/EtOAc+0.1%AcOH = 40/60) to give a carboxylic acid intermediate (3.4 mg, 78%) as a colorless oil.

To a solution of the carboxylic acid intermediate (3.4 mg) in $Et_2O$ (1.4 mL) was added $MgBr_2$•$OEt_2$ (4.2 mg, 16 μmol). The resulting white suspension was stirred at room temperature for 3 h and dried under reduced pressure. The residue was dissolved with 30% aq. MeOH containing 0.1% AcOH and syringe-filtered through PTFE membrane. The crude product was directly purified by RP-HPLC (ODS-HG-5 Φ4.6x250, 210 nm, 0.6 mL/min, eluent: 30% aq. MeOH + 0.1%AcOH) to give (*3R*,*7S*,*11S*)-OH-JA ($t_R$ = 21 min, 1.1 mg, 44%) as a colorless amorphous solid. The synthesized (*3R*,*7S*,*11S*)-OH-JA was the same as the latter isomer of (*3R*,*7S*,*11R/S*)-11-OH-JA in UPLC-MS/MS analysis (**Figure S3C**).

#### Plant materials

*Arabidopsis thaliana* ecotype Col-0 is the genetic background of wild-type and mutant lines used in this study. For biological assay using seedlings, seeds were surface-sterilized in 5% sodium hypochlorite with 0.3% Tween 20 and vernalized for 2–3 days at 4 °C. All seedlings were grown under a 16:8 h light (118 μmol $m^{-2}$ $sec^{-1}$)/dark photocycle at 22 °C in a CL-301 growth chamber (Tomy Seiko Co., Ltd, Tokyo, Japan), on 1/2 MS medium (MS medium). For biological assay using adult plants, seeds were grown directly in soil or 1:1 soil/vermiculite under a 12-h-light (75 μmol $m^{-1}$ $s^{-1}$, cool-white fluorescent light)/12-h-dark cycle at 22 °C.

The *joxQ* mutant was provided from XX and grown under the same condition as described above. XX

**UPLC-MS/MS analyses of wounding-induced JA and JA-Ile turnover in *A. thaliana* Col-0 and *joxQ* mutant line**

The leaves (ca. 100–200 mg FW) of the 5-week-old *A. thaliana* Col-0 and *joxQ* were nipped tightly with a pair of tweezers, avoiding the central veins. Damaged leaves were harvested at 0, 15, 30, 60, and 300 min after wounding. Each sample was weighed and frozen in liquid nitrogen. The frozen sample was ground using a Micro Homogenizing System (Micro Smash™; TOMY, Tokyo, Japan) with a zirconia bead and extracted in 1.5 mL EtOH for 16 h at 4 °C. After centrifugation at 4,300 rpm for 10 min, the supernatants were collected. The volatile components of the extract were removed by using nitrogen blowdown evaporator, and a solution of 150 µL 80% aq. MeOH was then added to the residue. The mixture was applied to an SPE C18 cartridge column (Sep-Pak C18 Plus Short Cartridge, 360 mg, Waters, Milford, MA, USA) and the column was eluted with a solution of 80% aq. MeOH (700 µL × 3). The volatile components and water in the eluents were removed by using nitrogen blowdown evaporator. The residue was dissolved with a solution of 100 µL MeOH, and the sample was examined by LC-MS/MS using an Ultivo Triple Quadrupole LC/MS system (Agilent Technologies, Santa Clara, CA, USA) operating in the negative mode. Poroshell 120 PFP 1.9 µm (Φ2.1 × 50 mm; Agilent Technologies) was used at 23 °C and a flow rate of 0.300 mL/min. The elution was performed using water-MeOH (95:5, v/v, isocratic), both containing 0.2% acetic acid (v/v). The liquid chromatograms were extracted by multiple reaction monitoring with the following ion transitions: 11-OH-JA 225 > 59 and 12-OH-JA 225 > 59.

**Pull-down experiments using fluorescein-tagged JAZ peptides**

For the pulldown experiment with Fl-JAZPs, purified GST-COI1 (5 nM), Fl-JAZPs (10 nM) and the ligands (1 - 100 μM) in 350 μL of incubation buffer were combined with anti-fluorescein antibody (ab19491; Abcam, Cambridge, UK ; 0.1 μL) and incubated for 10–15 h at 4 °C with rotation. After incubation, the samples were combined with SureBeads Protein A (10 μL in incubation buffer slurry; Bio-Rad, Hercules, CA, USA; 10 mg beads/ml). After 1.5 h of incubation at 4 °C with rotation, the samples were washed three times with 350 μL of phosphate-buffered saline containing 0.1% Tween 20. The washed beads were resuspended in 35 μL of SDS-PAGE loading buffer containing dithiothreitol (100 mM). After heating for 10 min at 60 °C, the samples were subjected to SDS-PAGE and analyzed by western blotting. The bound GST-COI1 was detected using anti-GST HRP conjugate (RPN1236; Cytiva ; 10000-fold dilution in blocking buffer).

**Growth inhibition assay**

Two-day-old Col-0 seedlings were transferred on a 1/2 MS plate in the absence or presence of 1–100 μm of each chemical and grown in a growth chamber under a 16:8 h light/dark photocycle at 22°C for 8 days. Images were then taken of each seedling with an α6400 digital camera (Sony Corp., Tokyo, Japan) and root length was measured using imagej, version 1.53S (http://imagej.net).

**Anthocyanin accumulation assay**

Seedlings were germinated on a 1/2 MS plate for 2 days and were transferred on 1/2 MS in the absence or presence of each chemical at concentrations in the range 1–100 μm to a growth chamber under a 16:8 h light/dark photocycle at 22°C for 8 days. Then, 16 seedlings were cut at the base of the leaf and weighed. For each sample, the seedlings were frozen and ground to a fine powder. Anthocyanin was extracted in aqueous methanol [HCl 0.1%; 50% methanol/sterilized water (v/v)] at 4°C for 3 h. The samples were then soaked in chloroform. Total anthocyanins were determined by measuring the $A_{530}$, and $A_{657}$ of the aqueous phase using a spectrophotometer (NanoPhotometerN60; Implen, München, Germany) and the content was calculated as: $A_{530} - 0.25 \times A_{657}$ per fresh weight (g).

**Plasmid constructions for JOX1/2/3/4 expression**

The coding sequence of *JOX1/2/3/4* (At5g05600, At3g55970, At2g38240, respectively) was obtained from TAIR database and JOX2/3/4 genes were amplified from an *A. thaliana* cDNA library using primers (JOX2-forward, 5′-ATGAACAAGAACAAGATTGATGTTAAGATCGAG-3′; JOX2-reverse, 5′-TCAACGAGGAGAAATATGAGATTCAACATG-3′, JOX3-forward, 5′-CATCATCATCATCACATGAATATCTTCCAAGACTGGC-3′; JOX3-reverse, 5′-TTAGCAGCCGGATCCCTATCGAGGGGATTTAAGTTCG-3′, JOX4-forward, 5′-CATCATCATCATCACATGGCTACATGCTGGCCTGAGC-3′; JOX4-reverse, 5′-TTAGCAGCCGGATCCTTATCTAGTTAATAACAGTG-3′). In the case of JOX1, we ordered Eurofins genomics to synthesis sequence and amplified them using primers (JOX1-forward, 5′- ATGAACAACCTAGACGAGATCAAGATCG-3′; JOX1-reverse, 5′- TCACCAATCTTTTGGAATCTCTAGGTAGTTC-3′). Using the In-Fusion HD Cloning kit (Takara Bio, Shiga, Japan), the PCR-amplified *JOX1-4* coding sequence was cloned into pET-*6His*-vector linearized with primers (JOX1-forward, 5′-CCAAAAGATTGGTGAGGATCCGGCTGCTAACAAAGCC -3′; JOX1-reverse, 5′-GTCTAGGTTGTTCATGTGATGATGATGATGATGGCCCATATG-3′, JOX2-forward,

5′- ATTTCTCCTCGTTGAGGATCCGGCTGCTAACAAAGC -3′; JOX2-reverse, 5′-CTTGTTCTTGTTCATGTGATGATGATGATGATGGCCCATATG-3′, JOX3/4-forward, 5′- GGATCCGGCTGCTAACAAAGCCC -3′; JOX3/4-reverse, 5′-GTGATGATGATGATGATGGCCCATATG -3′).

**Protein expression and purification**

The *6His-JOX1/2/3/4* genes in pET vector was expressed in *E. coli* BL21 (DE3) strain. *E. coli was* grown in LB medium supplemented with 50 μg/ml ampicillin and protein expression was induced by the addition of 0.2 mM isopropyl-β-D-thiogalactopyranoside when optical density at 600 nm got 0.6 with shaking at 16°C for 16 h. Cells were collected by centrifugation and lysed by sonication with lysis buffer (20 mM Tris–HCl, 150 mM NaCl, 20 mM imidazole [pH 7.5]). Recombinant protein was collected from Ni Sepharose™ High Performance (GE Healthcare) using elution buffer (20 mM Tris–HCl, 150 mM NaCl, 300 mM imidazole [pH 7.5])

The obtained crude fraction of tag-free COI1 was concentrated. The concentrated fraction was purified by gel filtration chromatography on the AKTA system (Amersham Pharmacia Biotech, column: Hiload 16/60 Superdex 200 prepgrade (1CV=120mL), elution buffer: 10 mM HEPES, 150 mM NaCl, pH 7.4, flowrate: 0.50 mL/min, detection: 280, 254, and 220nm, fraction volume: 2.0 mL, equilibration: 1.1 CV, elution 1.2 CV). The fractions of pure 6His-JOX1-4 were collected, and the concentration was determined by UV absorption at 280 nm using NanoPhotometerN60 (IMPLEN GmbH, Germany).

[1]H, [13]C NMR spectra

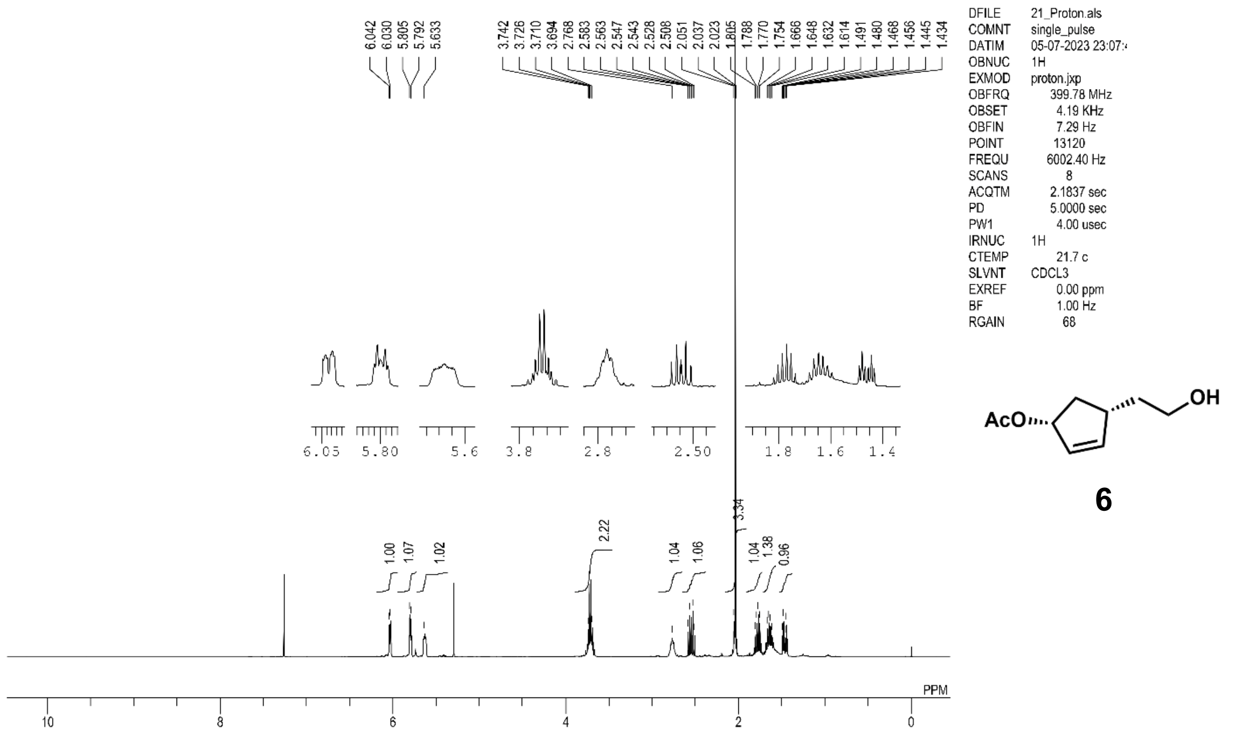


6

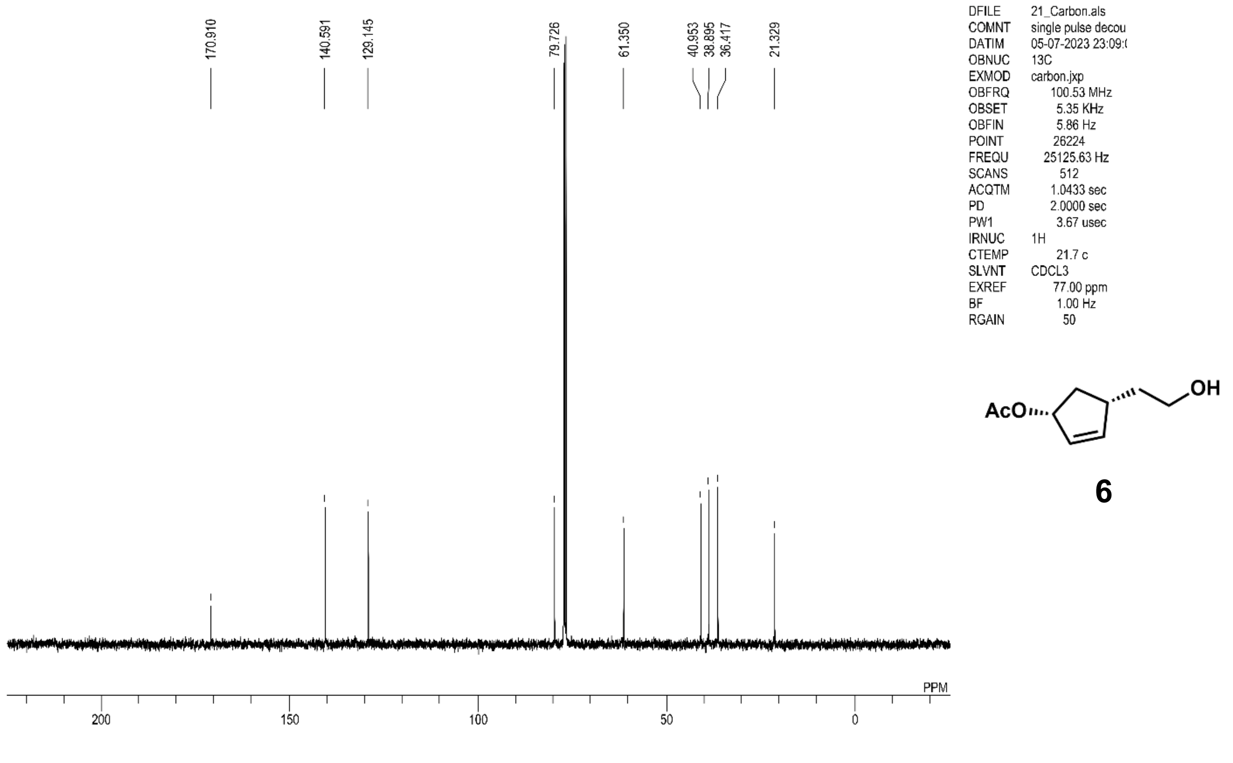


6

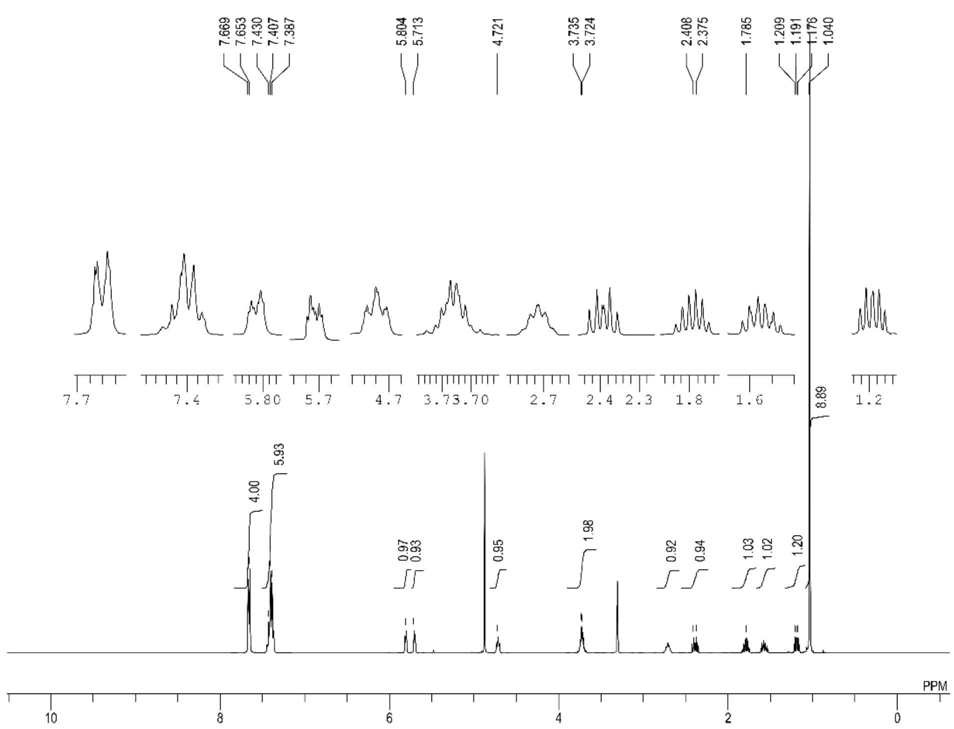

7.669 7.653 7.430 7.407 7.387 5.804 5.713 4.721 3.735 3.724 2.408 2.375 1.785 1.209 1.191 1.176 1.040
4.00 5.93 0.97 0.93 0.95 1.98 0.92 0.94 1.03 1.02 1.20 8.89
PPM
DFILE 22_MeOD_Proton.
COMNT single_pulse
DATIM 20-07-2023 20:16:
OBNUC 1H
EXMOD proton.jxp
OBFRQ 399.78 MHz
OBSET 4.19 KHz
OBFIN 7.29 Hz
POINT 13120
FREQU 6002.40 Hz
SCANS 8
ACQTM 2.1837 sec
PD 5.0000 sec
PW1 4.00 usec
IRNUC 1H
CTEMP 21.8 c
SLVNT CD3OD
EXREF 3.31 ppm
BF 1.00 Hz
RGAIN 64


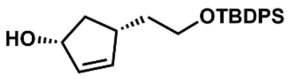

HO
OTBDPS


**7**

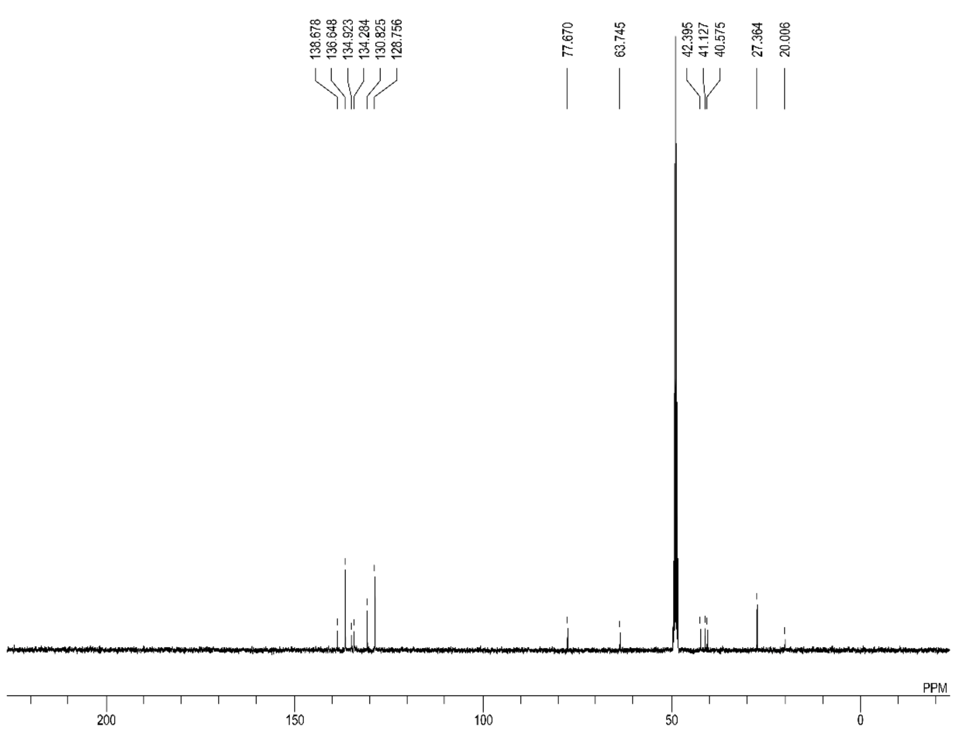

138.678 136.648 134.923 134.284 130.825 128.756 77.670 63.745 42.395 41.127 40.575 27.364 20.006
PPM
DFILE 22_MeOD_Carbon
COMNT single pulse decou
DATIM 12-07-2023 16:57:
OBNUC 13C
EXMOD carbon.jxp
OBFRQ 100.53 MHz
OBSET 5.35 KHz
OBFIN 5.86 Hz
POINT 26224
FREQU 25125.63 Hz
SCANS 512
ACQTM 1.0433 sec
PD 2.0000 sec
PW1 3.67 usec
IRNUC 1H
CTEMP 21.6 c
SLVNT CD3OD
EXREF 49.00 ppm
BF 1.00 Hz
RGAIN 50


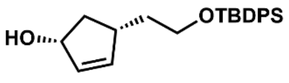

HO
OTBDPS


**7**

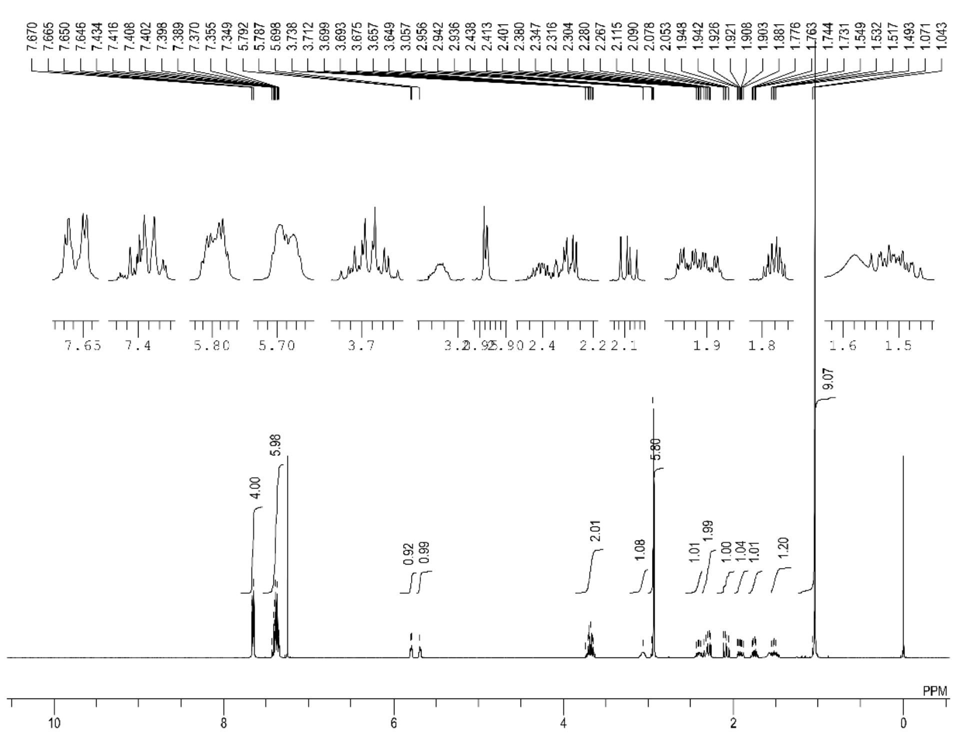


DFILE 23_Proton.als
COMNT single_pulse
DATIM 11-07-2023 21:37:[illegible]
OBNUC 1H
EXMOD proton.jxp
OBFRQ 399.78 MHz
OBSET 4.19 KHz
OBFIN 7.29 Hz
POINT 13120
FREQU 6002.40 Hz
SCANS 8
ACQTM 2.1837 sec
PD 5.0000 sec
PW1 4.00 usec
IRNUC 1H
CTEMP 21.8 c
SLVNT CDCL3
EXREF 0.00 ppm
BF 1.00 Hz
RGAIN 74

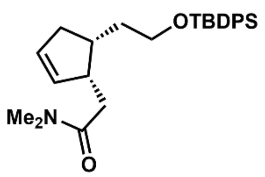


**8**

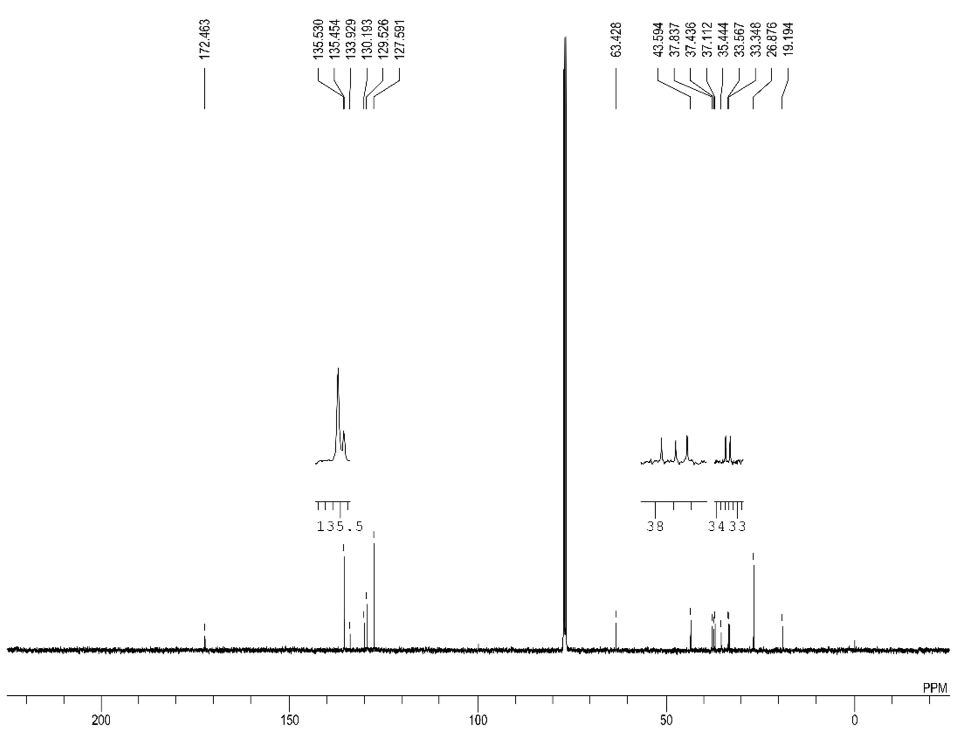


DFILE 23_Carbon.als
COMNT single pulse decou
DATIM 24-07-2023 16:05:[illegible]
OBNUC 13C
EXMOD carbon.jxp
OBFRQ 100.53 MHz
OBSET 5.35 KHz
OBFIN 5.86 Hz
POINT 26224
FREQU 25125.63 Hz
SCANS 2048
ACQTM 1.0433 sec
PD 2.0000 sec
PW1 3.67 usec
IRNUC 1H
CTEMP 21.9 c
SLVNT CDCL3
EXREF 77.00 ppm
BF 1.00 Hz
RGAIN 50

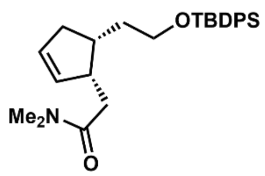


**8**

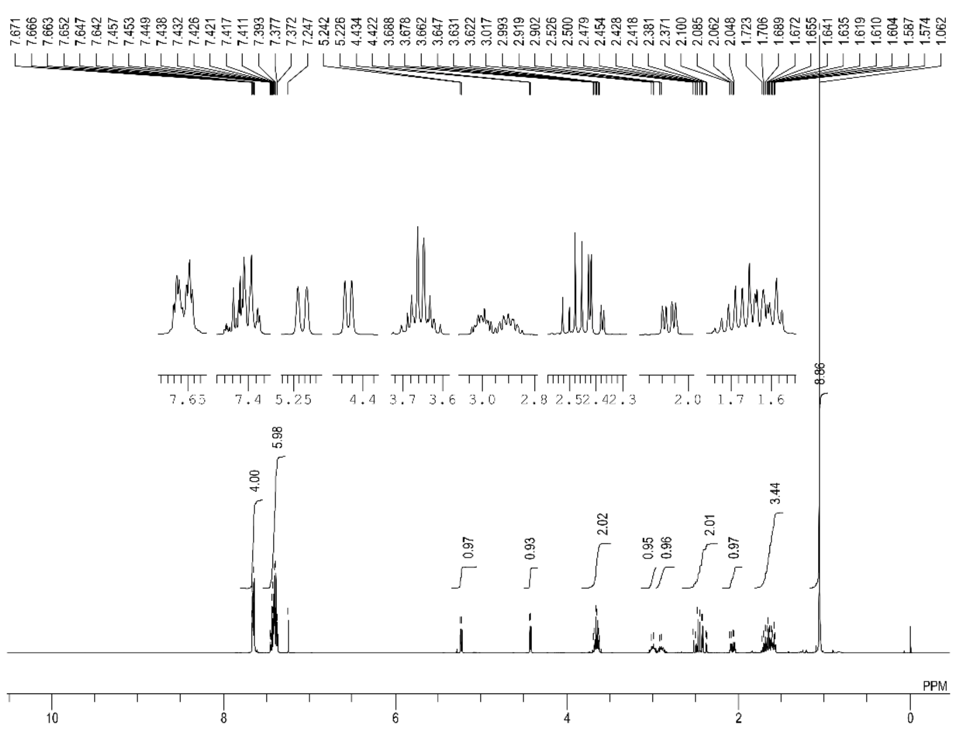
7.671 7.666 7.663 7.652 7.647 7.642 7.457 7.453 7.449 7.438 7.432 7.426 7.421 7.417 7.411 7.393 7.377 7.372 7.247 5.242 5.226 4.434 4.422 3.688 3.678 3.662 3.647 3.631 3.622 3.017 2.993 2.919 2.902 2.526 2.500 2.479 2.454 2.428 2.418 2.381 2.371 2.100 2.085 2.062 2.048 1.723 1.706 1.689 1.672 1.655 1.641 1.635 1.619 1.610 1.604 1.587 1.574 1.062
8.86
4.00
5.98
0.97
0.93
2.02
0.95
0.96
2.01
0.97
3.44
PPM
10
8
6
4
2
0
DFILE 24_Proton.als
COMNT single_pulse
DATIM 20-07-2023 22:18:[illegible]
OBNUC 1H
EXMOD proton.jxp
OBFRQ 399.78 MHz
OBSET 4.19 KHz
OBFIN 7.29 Hz
POINT 13120
FREQU 6002.40 Hz
SCANS 8
ACQTM 2.1837 sec
PD 5.0000 sec
PW1 4.00 usec
IRNUC 1H
CTEMP 21.8 c
SLVNT CDCL3
EXREF 0.00 ppm
BF 1.00 Hz
RGAIN 60

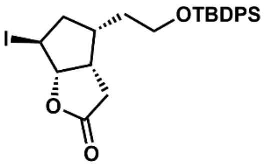
I
OTBDPS
O
O

**9**

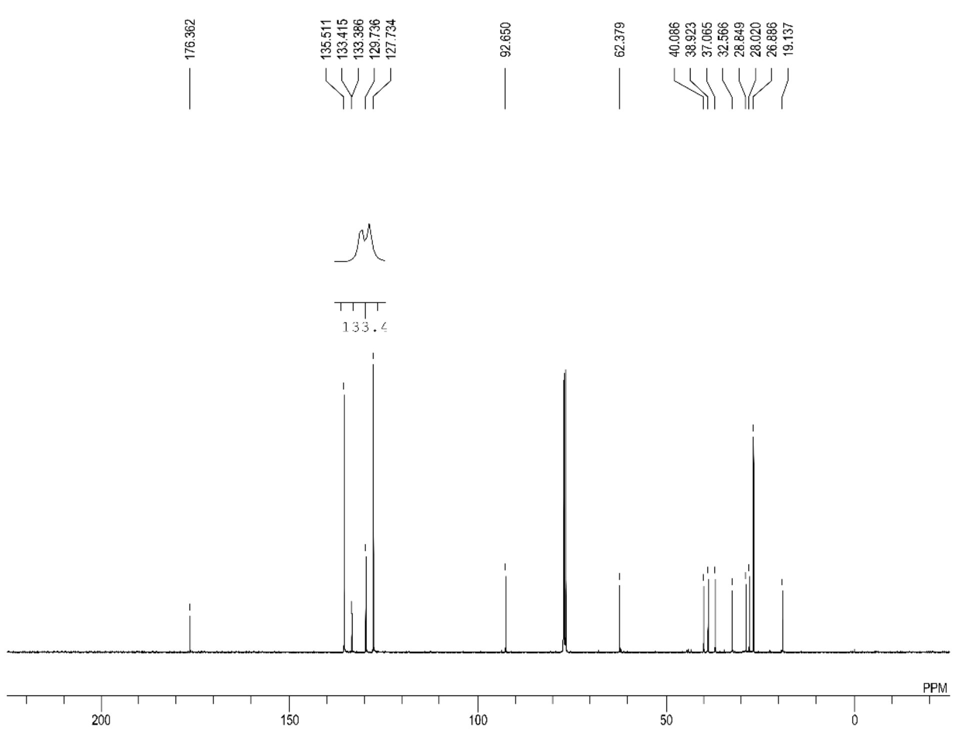
176.362
135.511 133.415 133.386 129.736 127.734
92.650
62.379
40.086 38.923 37.065 32.566 28.849 28.020 26.886 19.137
133.4
PPM
200
150
100
50
0
DFILE 24_Carbon.als
COMNT single pulse decou[illegible]
DATIM 20-07-2023 22:19:[illegible]
OBNUC 13C
EXMOD carbon.jxp
OBFRQ 100.53 MHz
OBSET 5.35 KHz
OBFIN 5.86 Hz
POINT 26224
FREQU 25125.63 Hz
SCANS 8192
ACQTM 1.0433 sec
PD 2.0000 sec
PW1 3.67 usec
IRNUC 1H
CTEMP 21.7 c
SLVNT CDCL3
EXREF 77.00 ppm
BF 1.00 Hz
RGAIN 50

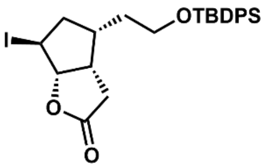
I
OTBDPS
O
O

**9**

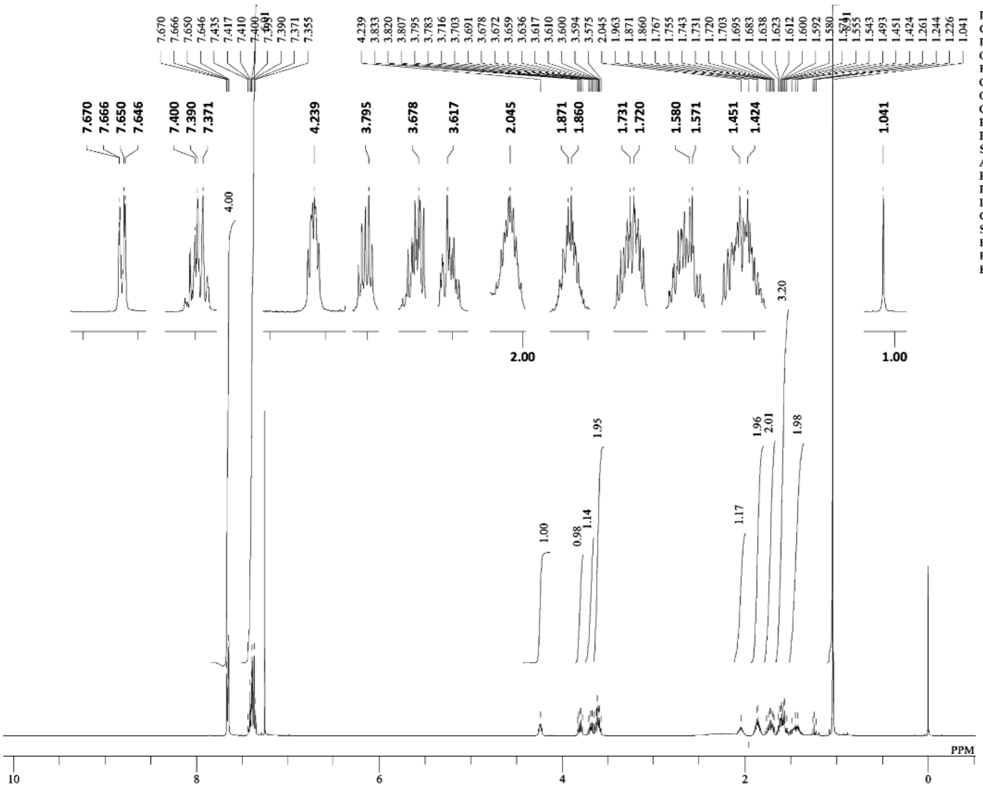
DFILE 2_1H_KN_exp42_step9_fr23-32_
COMNT single_pulse
DATIM 18-12-2023 15:52:13
OBNUC 1H
EXMOD proton.jxp
OBFRQ 399.78 MHz
OBSET 4.19 KHz
OBFIN 7.29 Hz
POINT 13120
FREQU 6002.40 Hz
SCANS 8
ACQTM 2.1837 sec
PD 5.0000 sec
PW1 4.00 usec
IRNUC 1H
CTEMP 20.6 c
SLVNT CDCL3
EXREF 0.00 ppm
BF 1.00 Hz
RGAIN 74
PPM

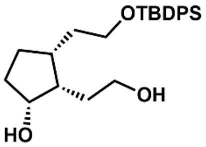
OTBDPS
OH
HO

**11**

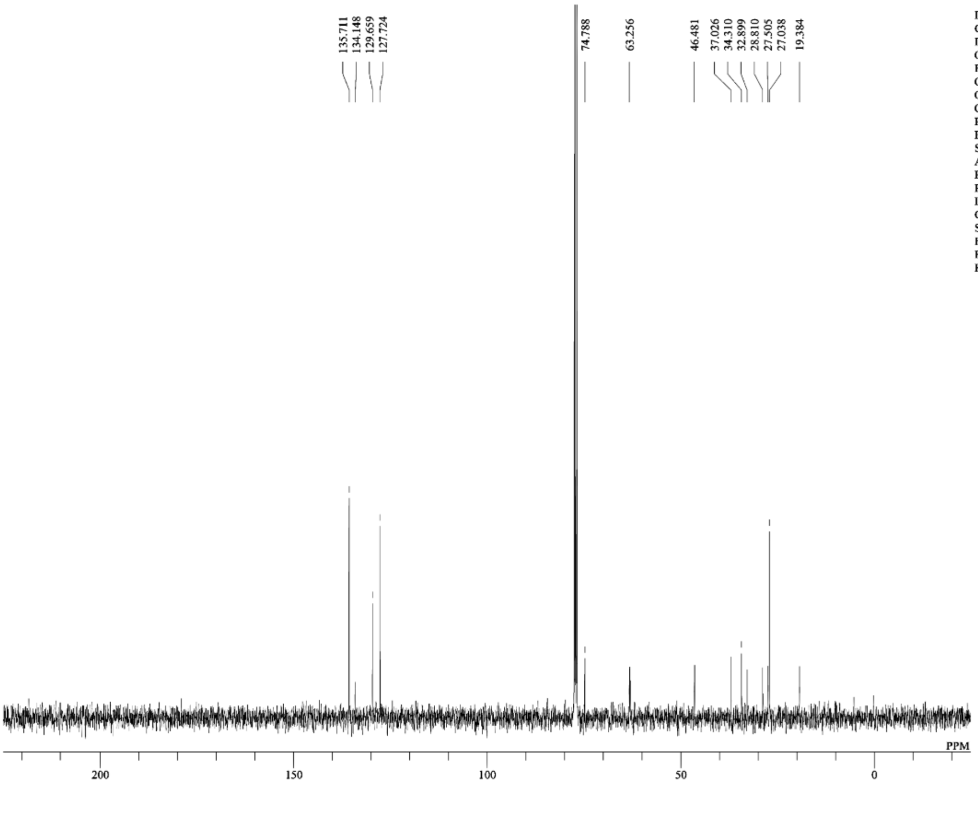
135.711
134.148
129.659
127.724
74.788
63.256
46.481
37.026
34.310
32.899
28.810
27.505
27.038
19.384
DFILE 2_13C_KN_exp42_step9_fr23-32
COMNT single pulse decoupled gated NOE
DATIM 18-12-2023 15:53:26
OBNUC 13C
EXMOD carbon.jxp
OBFRQ 100.53 MHz
OBSET 5.35 KHz
OBFIN 5.86 Hz
POINT 26224
FREQU 25125.63 Hz
SCANS 256
ACQTM 1.0433 sec
PD 2.0000 sec
PW1 3.67 usec
IRNUC 1H
CTEMP 20.6 c
SLVNT CDCL3
EXREF 224.97 ppm
BF 1.00 Hz
RGAIN 50
PPM

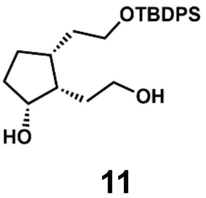
OTBDPS
OH
HO

**11**

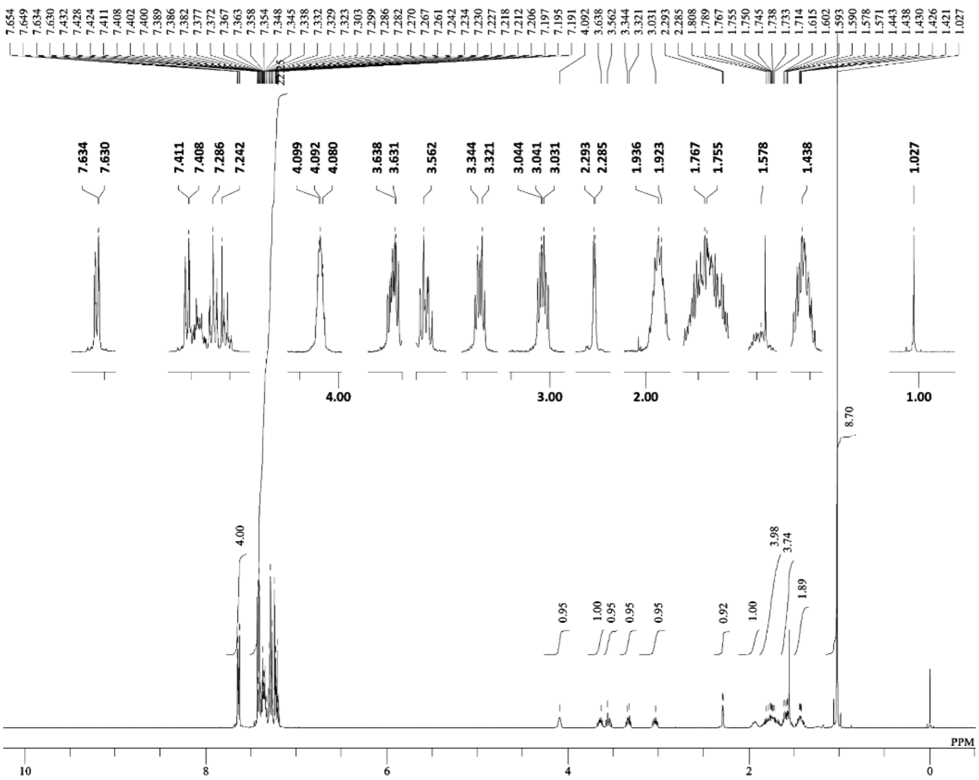


DFILE 3_1H_KN_exp69_step3_TM_Pro
COMNT single_pulse
DATIM 08-07-2024 18:50:21
OBNUC 1H
EXMOD proton.jxp
OBFRQ 399.78 MHz
OBSET 4.19 KHz
OBFIN 7.29 Hz
POINT 13120
FREQU 6002.40 Hz
SCANS 8
ACQTM 2.1837 sec
PD 5.0000 sec
PW1 3.60 usec
IRNUC 1H
CTEMP 24.6 c
SLVNT CDCL3
EXREF 0.00 ppm
BF 1.00 Hz
RGAIN 72

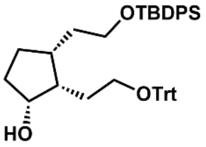


**12**

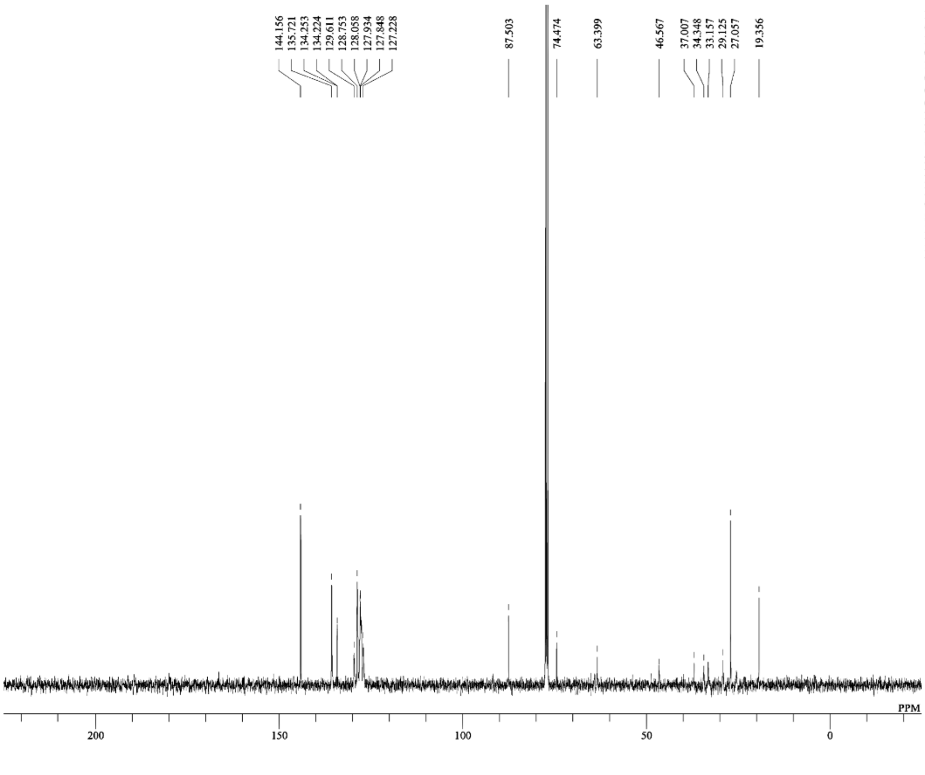


DFILE 3_13C_KN_exp69_step3_TM_Ca
COMNT single pulse decoupled gated NOE
DATIM 08-07-2024 18:23:13
OBNUC 13C
EXMOD carbon.jxp
OBFRQ 100.53 MHz
OBSET 5.35 KHz
OBFIN 5.86 Hz
POINT 26224
FREQU 25125.63 Hz
SCANS 512
ACQTM 1.0433 sec
PD 2.0000 sec
PW1 3.50 usec
IRNUC 1H
CTEMP 24.7 c
SLVNT CDCL3
EXREF 224.97 ppm
BF 1.00 Hz
RGAIN 50

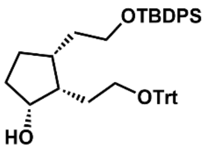


**12**

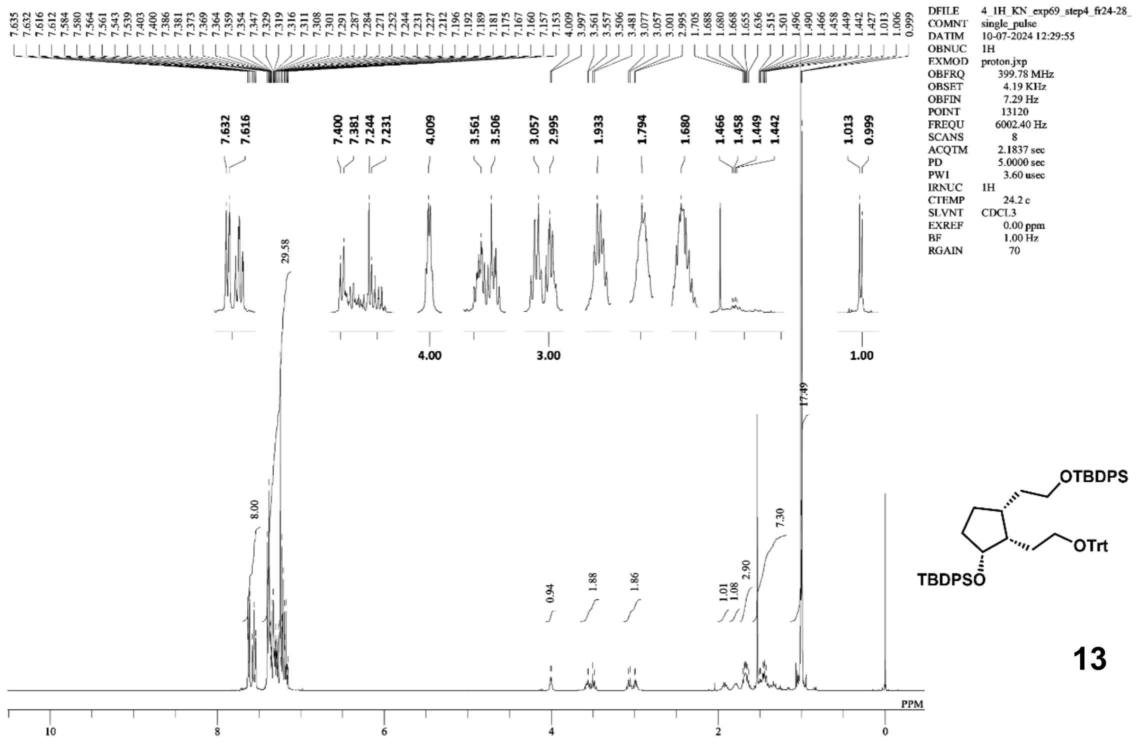

13

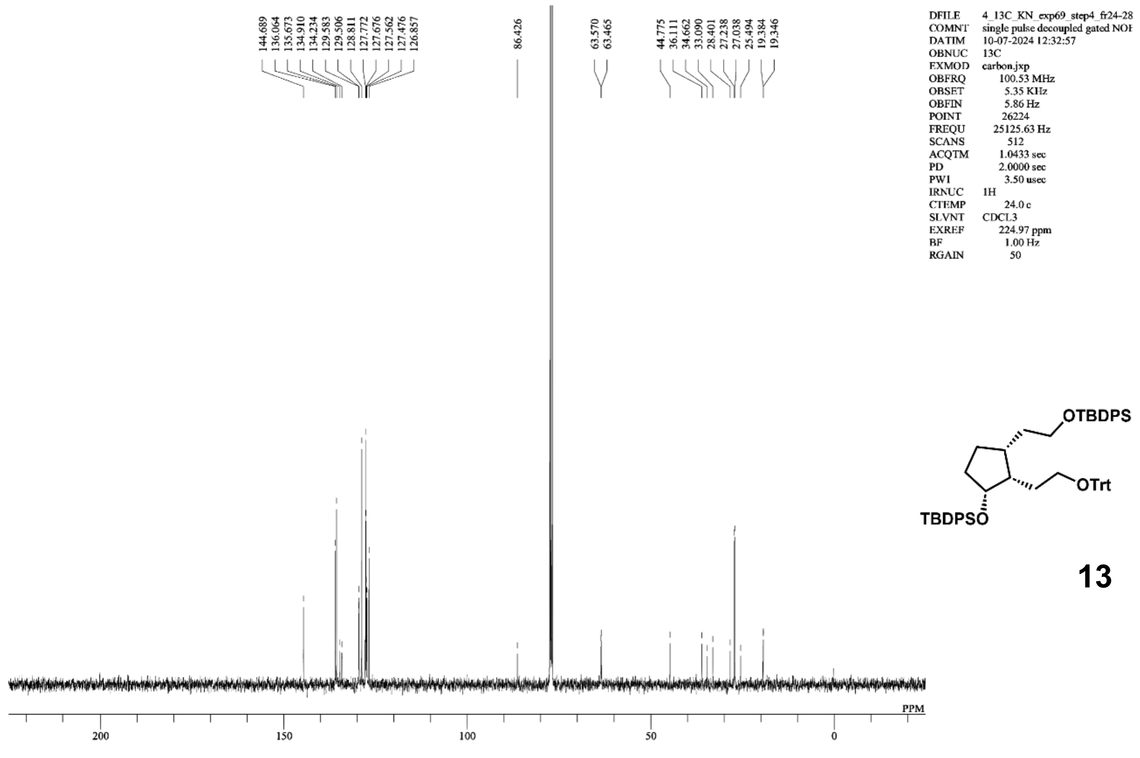

13

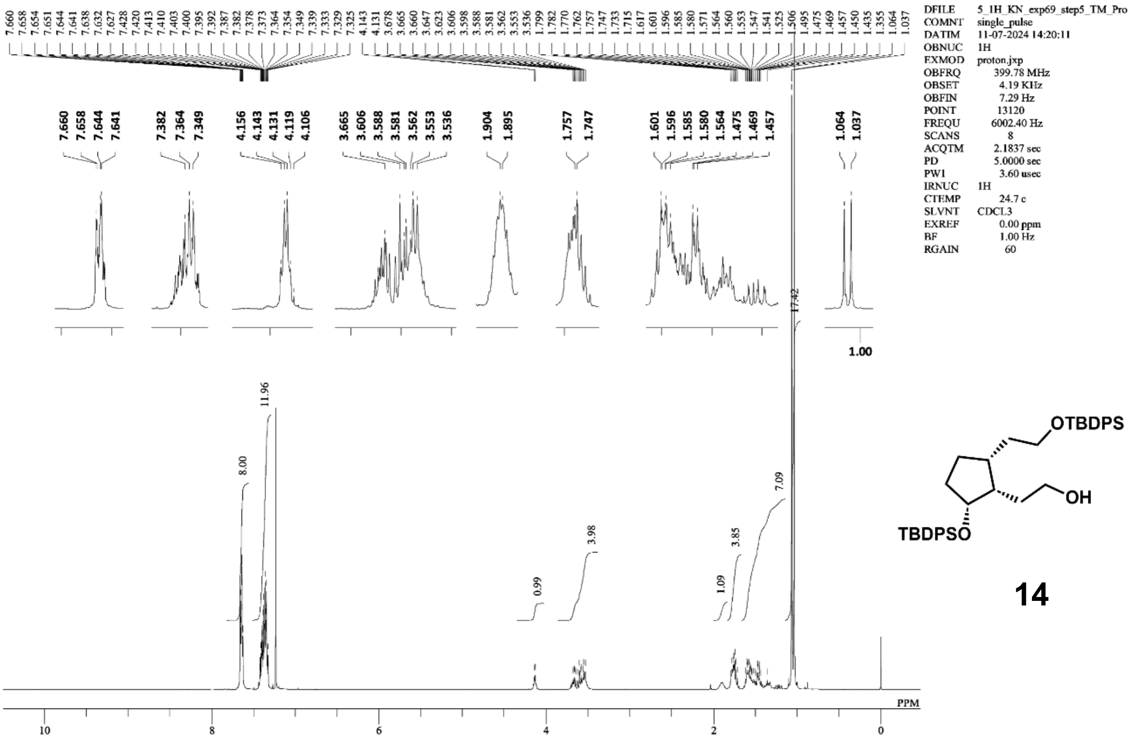
DFILE 5_1H_KN_exp69_step5_TM_Pro
COMNT single_pulse
DATIM 11-07-2024 14:20:11
OBNUC 1H
EXMOD proton.jxp
OBFRQ 399.78 MHz
OBSET 4.19 KHz
OBFIN 7.29 Hz
POINT 13120
FREQU 6002.40 Hz
SCANS 8
ACQTM 2.1837 sec
PD 5.0000 sec
PW1 3.60 usec
IRNUC 1H
CTEMP 24.7 c
SLVNT CDCL3
EXREF 0.00 ppm
BF 1.00 Hz
RGAIN 60
OTBDPS
OH
TBDPSO
PPM

14

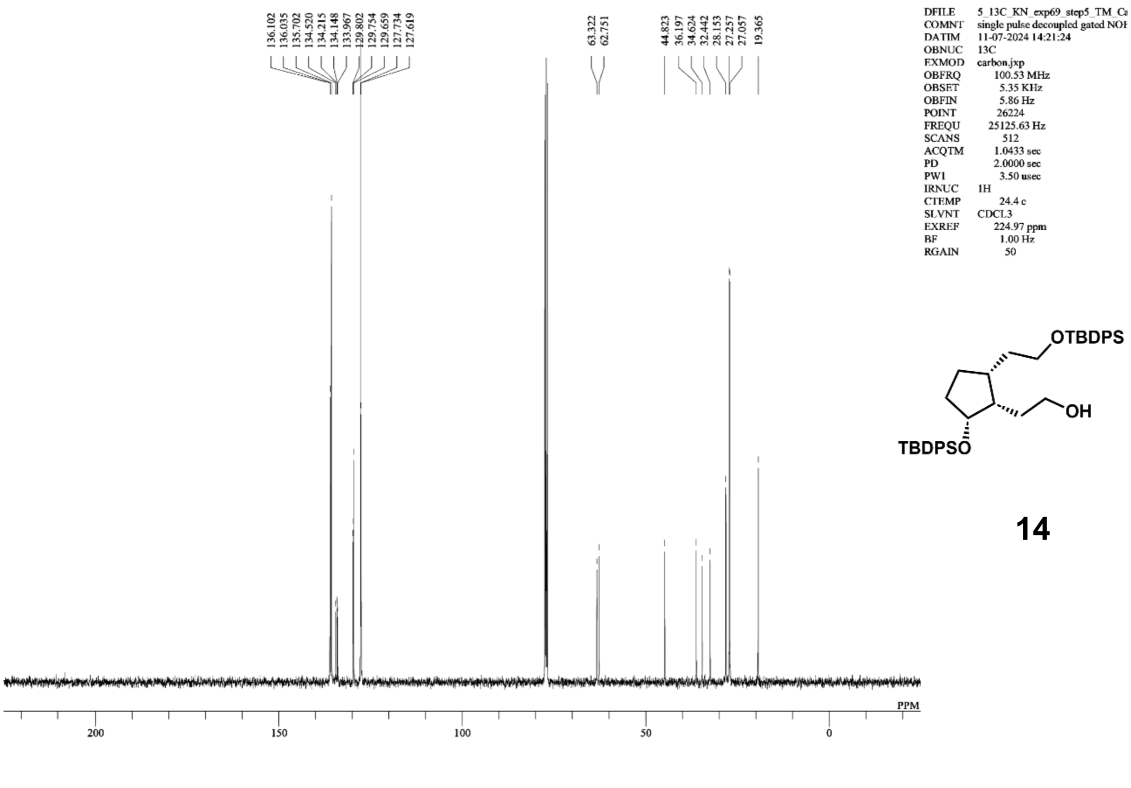
DFILE 5_13C_KN_exp69_step5_TM_Ca
COMNT single pulse decoupled gated NOE
DATIM 11-07-2024 14:21:24
OBNUC 13C
EXMOD carbon.jxp
OBFRQ 100.53 MHz
OBSET 5.35 KHz
OBFIN 5.86 Hz
POINT 26224
FREQU 25125.63 Hz
SCANS 512
ACQTM 1.0433 sec
PD 2.0000 sec
PW1 3.50 usec
IRNUC 1H
CTEMP 24.4 c
SLVNT CDCL3
EXREF 224.97 ppm
BF 1.00 Hz
RGAIN 50
OTBDPS
OH
TBDPSO
PPM

14

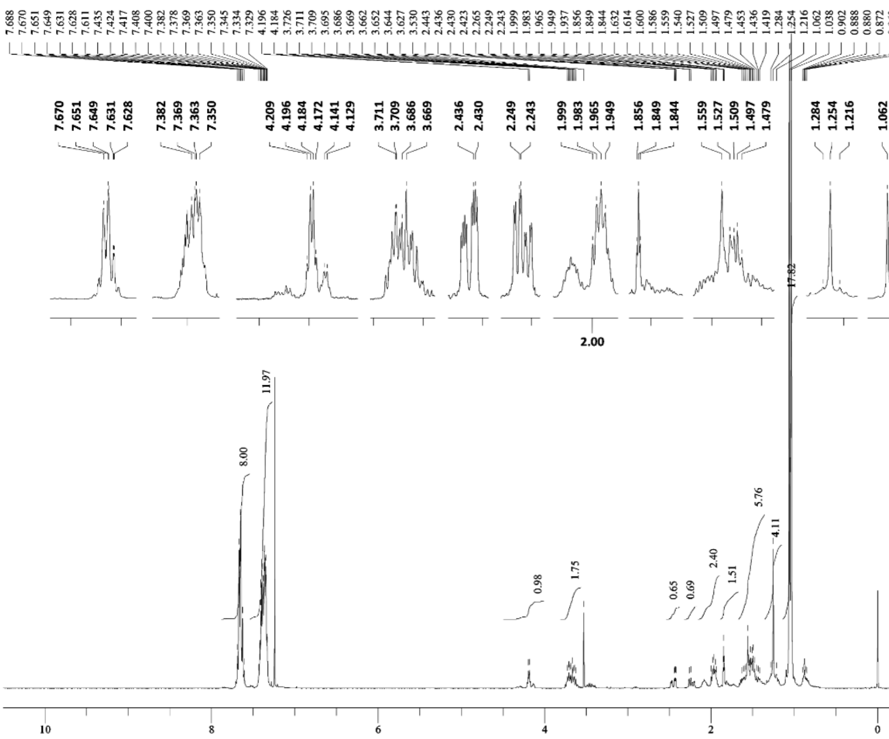
DFILE 7_1H_KN_exp69_step7_TM_Pro
COMNT single_pulse
DATIM 17-07-2024 19:41:12
OBNUC 1H
EXMOD proton.jxp
OBFRQ 399.78 MHz
OBSET 4.19 KHz
OBFIN 7.29 Hz
POINT 13120
FREQU 6002.40 Hz
SCANS 8
ACQTM 2.1837 sec
PD 5.0000 sec
PW1 3.60 usec
IRNUC 1H
CTEMP 22.1 c
SLVNT CDCL3
EXREF 0.00 ppm
BF 1.00 Hz
RGAIN 66
PPM

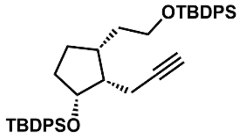
OTBDPS
TBDPSO

**2**

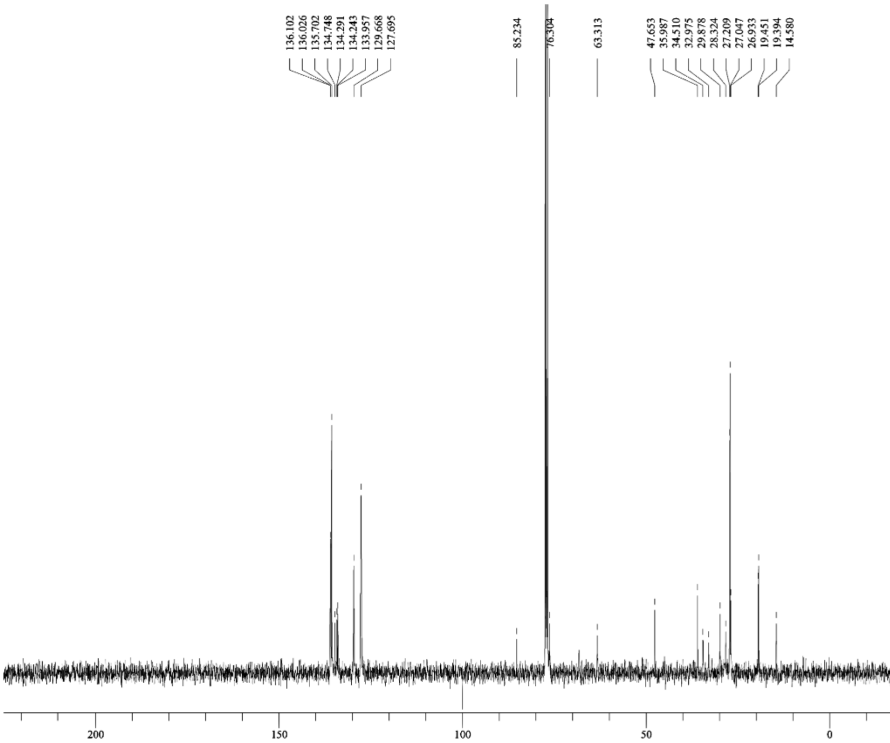
DFILE 7_13C_KN_exp69_step7_TM_Ca
COMNT single pulse decoupled gated NOE
DATIM 17-07-2024 21:54:52
OBNUC 13C
EXMOD carbon.jxp
OBFRQ 100.53 MHz
OBSET 5.35 KHz
OBFIN 5.86 Hz
POINT 26224
FREQU 25125.63 Hz
SCANS 512
ACQTM 1.0433 sec
PD 2.0000 sec
PW1 3.50 usec
IRNUC 1H
CTEMP 22.1 c
SLVNT CDCL3
EXREF 224.97 ppm
BF 1.00 Hz
RGAIN 50
PPM

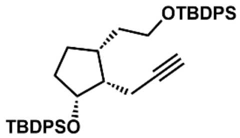
OTBDPS
TBDPSO

**2**

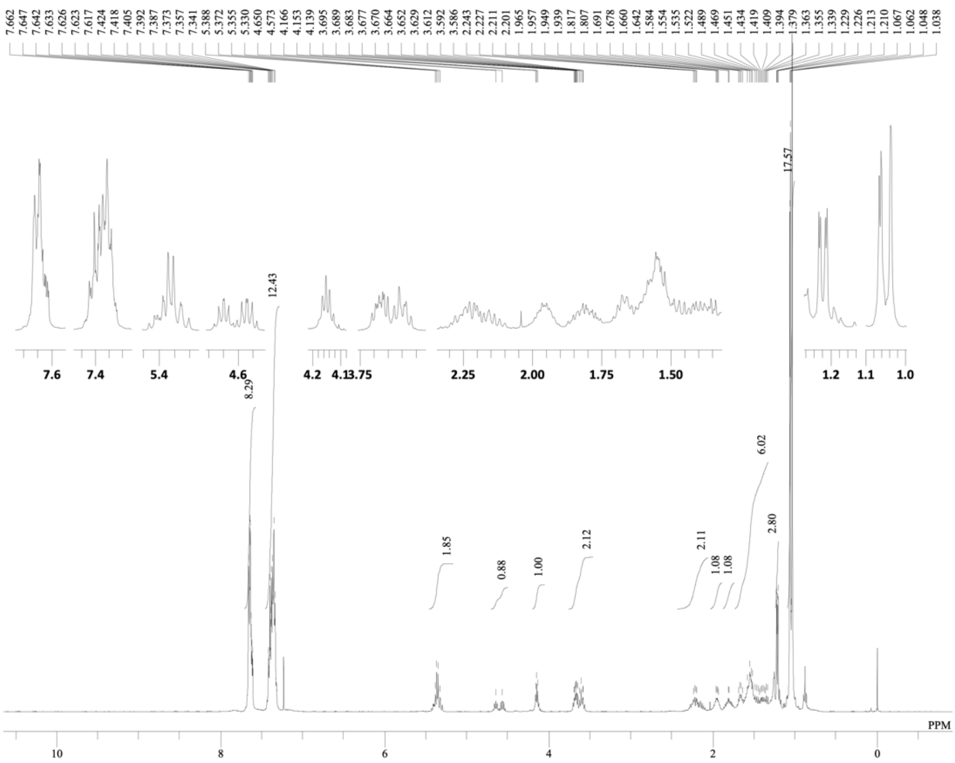


DFILE TO-ex11-st2_Proton-6-1.als
COMNT single_pulse
DATIM 15-11-2024 22:02:15
OBNUC 1H
EXMOD proton.jxp
OBFRQ 399.78 MHz
OBSET 4.19 KHz
OBFIN 7.29 Hz
POINT 13120
FREQU 6002.40 Hz
SCANS 8
ACQTM 2.1837 sec
PD 5.0000 sec
PW1 3.60 usec
IRNUC 1H
CTEMP 20.8 c
SLVNT CDCL3
EXREF 0.00 ppm
BF 1.00 Hz
RGAIN 60

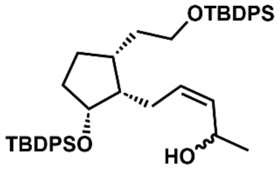


**3**

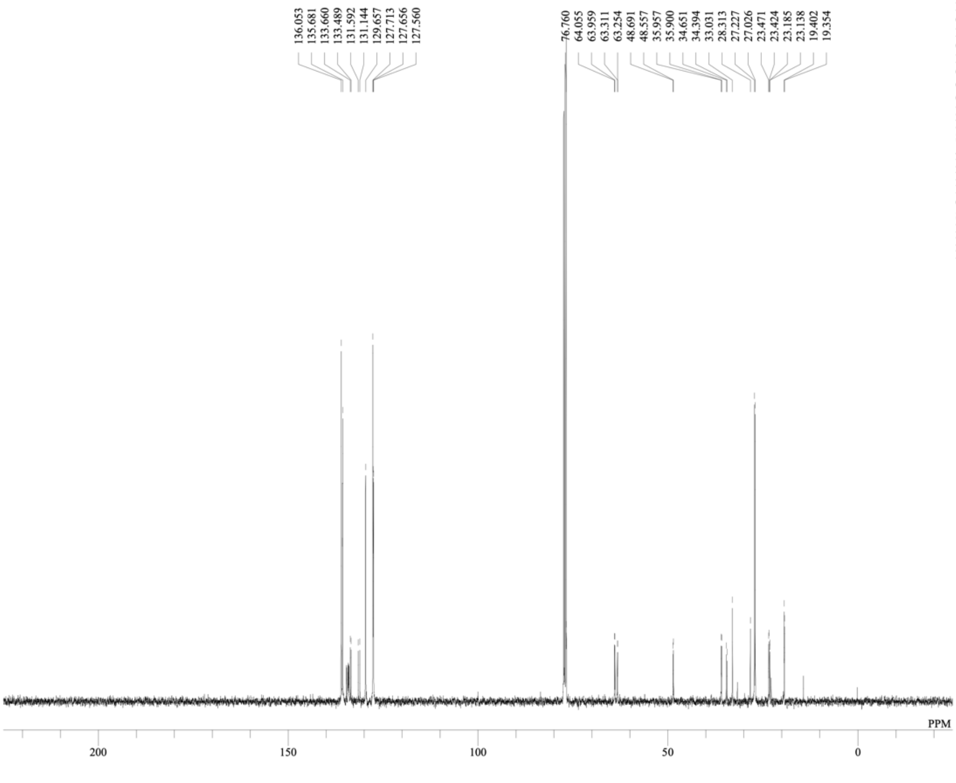


DFILE TO-ex11-st2_Carbon-2-1.als
COMNT single pulse decoupled gated NOE
DATIM 15-11-2024 21:30:40
OBNUC 13C
EXMOD carbon.jxp
OBFRQ 100.53 MHz
OBSET 5.35 KHz
OBFIN 5.86 Hz
POINT 26224
FREQU 25125.63 Hz
SCANS 601
ACQTM 1.0433 sec
PD 2.0000 sec
PW1 3.50 usec
IRNUC 1H
CTEMP 20.8 c
SLVNT CDCL3
EXREF 77.16 ppm
BF 1.00 Hz
RGAIN 50

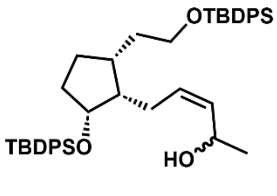


**3**

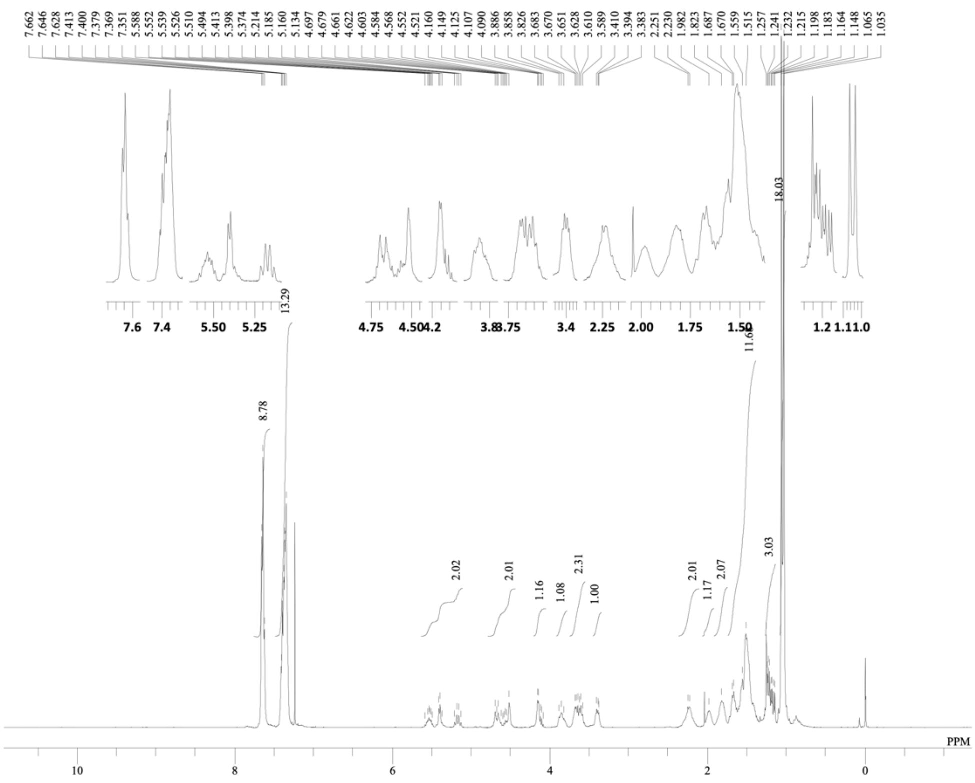
DFILE TO-ex12-st1_Proton-4-1.als
COMNT single_pulse
DATIM 22-11-2024 16:23:50
OBNUC 1H
EXMOD proton.jxp
OBFRQ 399.78 MHz
OBSET 4.19 KHz
OBFIN 7.29 Hz
POINT 13120
FREQU 6002.40 Hz
SCANS 8
ACQTM 2.1837 sec
PD 5.0000 sec
PW1 3.60 usec
IRNUC 1H
CTEMP 20.6 c
SLVNT CDCL3
EXREF 0.00 ppm
BF 1.00 Hz
RGAIN 68
PPM

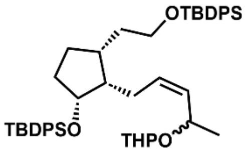
OTBDPS
TBDPSO
THPO

**4**

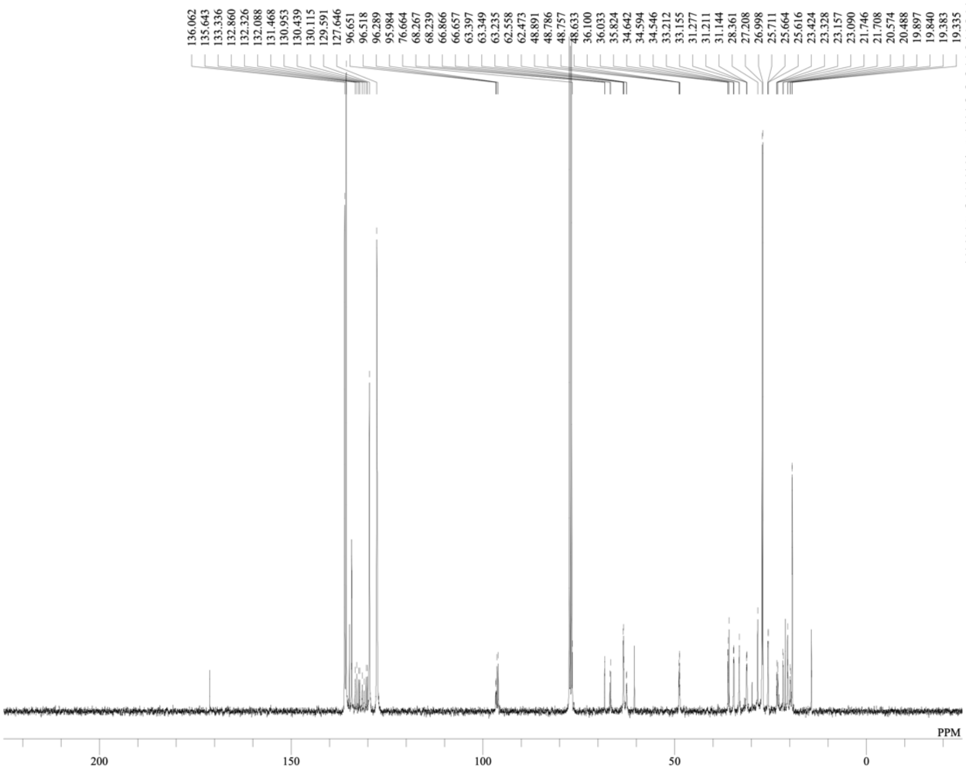
DFILE TO-ex12-st1_Carbon-1-1.als
COMNT single pulse decoupled gated NOE
DATIM 21-11-2024 21:26:26
OBNUC 13C
EXMOD carbon.jxp
OBFRQ 100.53 MHz
OBSET 5.35 KHz
OBFIN 5.86 Hz
POINT 26224
FREQU 25125.63 Hz
SCANS 1024
ACQTM 1.0433 sec
PD 2.0000 sec
PW1 3.50 usec
IRNUC 1H
CTEMP 20.7 c
SLVNT CDCL3
EXREF 77.16 ppm
BF 1.00 Hz
RGAIN 50
PPM

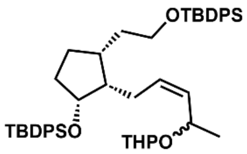
OTBDPS
TBDPSO
THPO

**4**

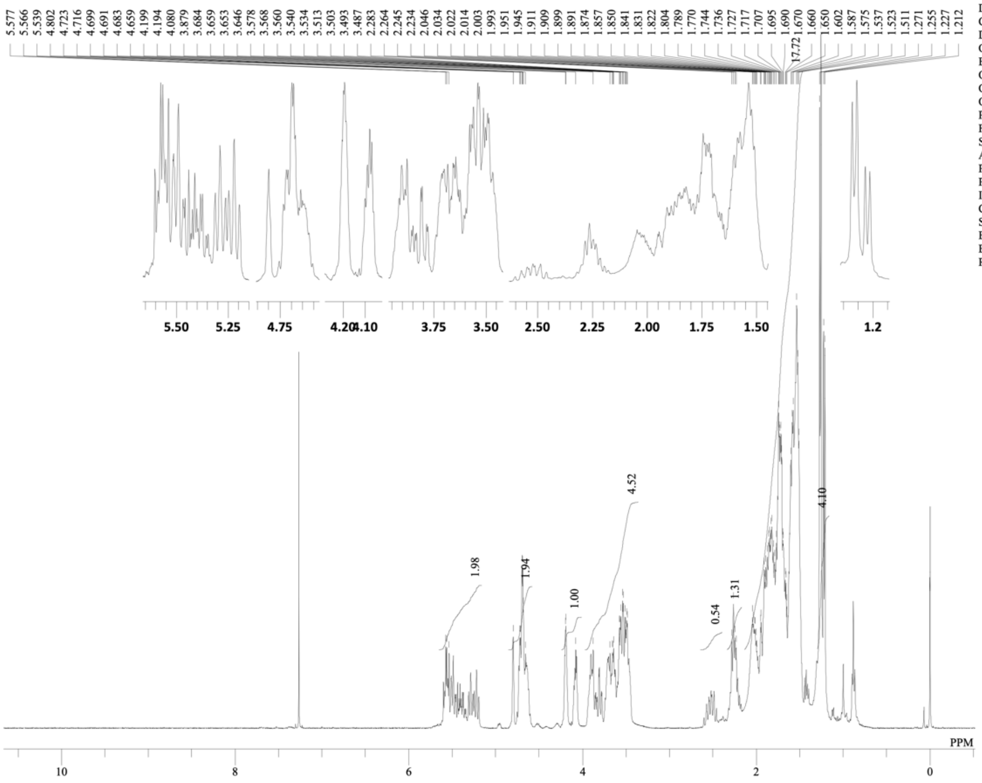


DFILE TO-ex12-st2_Proton-3-1.als
COMNT single_pulse
DATIM 27-11-2024 15:12:02
OBNUC 1H
EXMOD proton.jxp
OBFRQ 399.78 MHz
OBSET 4.19 KHz
OBFIN 7.29 Hz
POINT 13120
FREQU 6002.40 Hz
SCANS 8
ACQTM 2.1837 sec
PD 5.0000 sec
PW1 3.60 usec
IRNUC 1H
CTEMP 20.7 c
SLVNT CDCL3
EXREF 0.00 ppm
BF 1.00 Hz
RGAIN 68

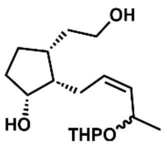


**5**

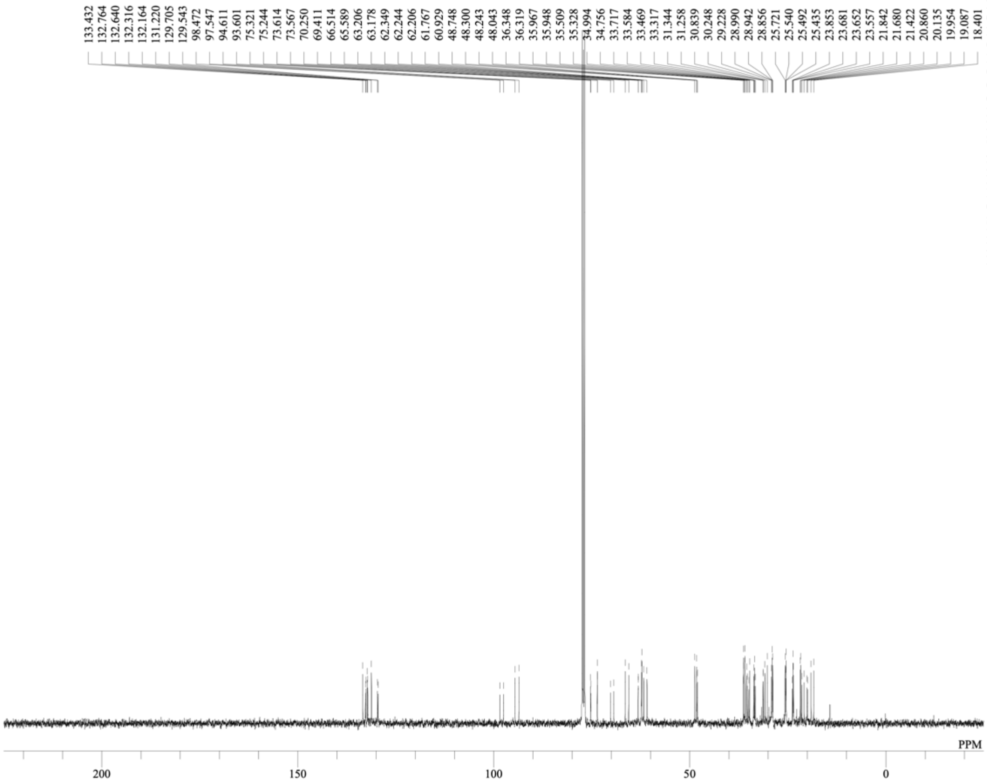


DFILE To-ex12-st2_Carbon-1-1.als
COMNT single pulse decoupled gated NOE
DATIM 27-11-2024 12:49:10
OBNUC 13C
EXMOD carbon.jxp
OBFRQ 100.53 MHz
OBSET 5.35 KHz
OBFIN 5.86 Hz
POINT 26224
FREQU 25125.63 Hz
SCANS 1024
ACQTM 1.0433 sec
PD 2.0000 sec
PW1 3.50 usec
IRNUC 1H
CTEMP 21.0 c
SLVNT CDCL3
EXREF 77.16 ppm
BF 1.00 Hz
RGAIN 50

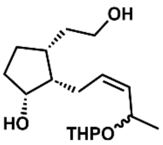


**5**

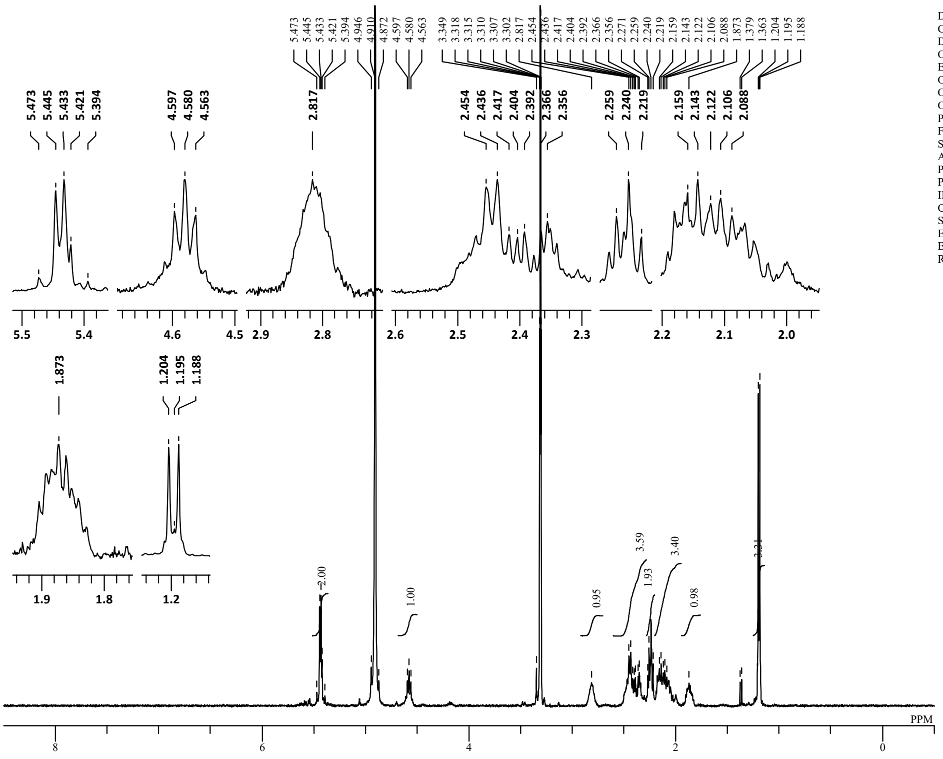


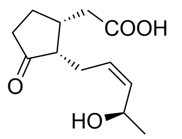


(*3R,7S,11R*)-11-OH-JA

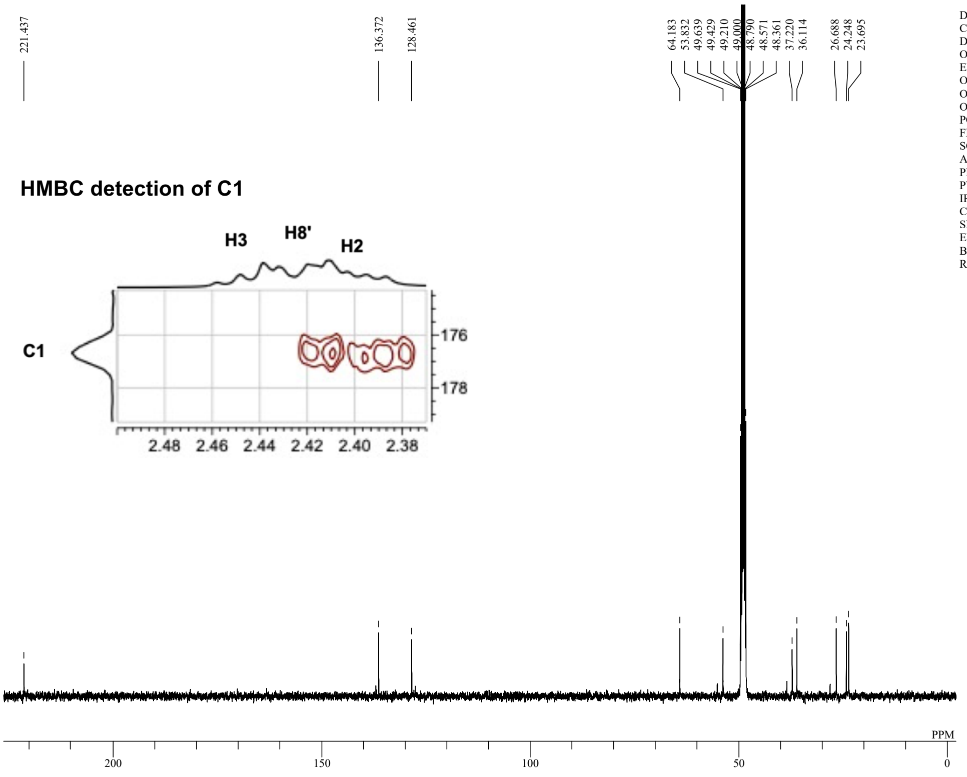


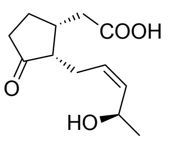


(*3R,7S,11R*)-11-OH-JA

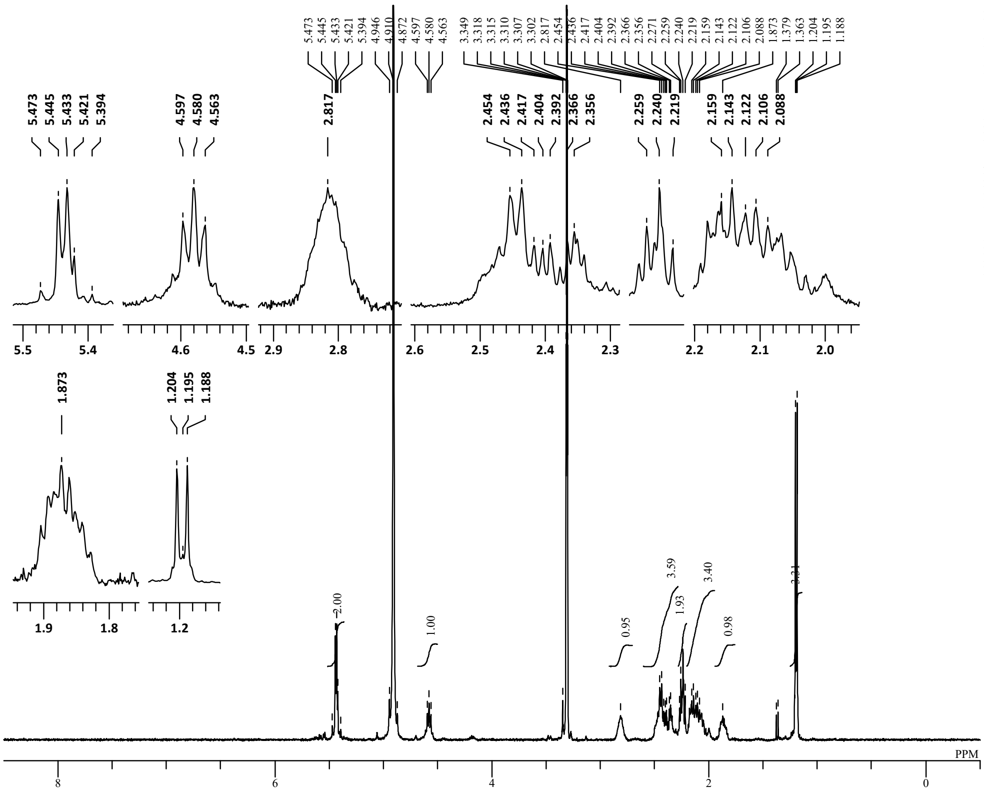


| | |
|---|---|
| DFILE | TK04198-11S-OH-JA_Proton-1-1 |
| COMNT | single_pulse |
| DATIM | 10-03-2025 23:11:05 |
| OBNUC | 1H |
| EXMOD | proton.jxp |
| OBFRQ | 399.78 MHz |
| OBSET | 4.19 KHz |
| OBFIN | 7.29 Hz |
| POINT | 16400 |
| FREQU | 7503.00 Hz |
| SCANS | 8 |
| ACQTM | 2.1837 sec |
| PD | 5.0000 sec |
| PW1 | 3.60 usec |
| IRNUC | 1H |
| CTEMP | 18.7 c |
| SLVNT | CD3OD |
| EXREF | 3.31 ppm |
| BF | 1.00 Hz |
| RGAIN | 86 |

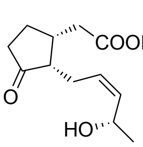


(*3R,7S,11S*)-11-OH-JA

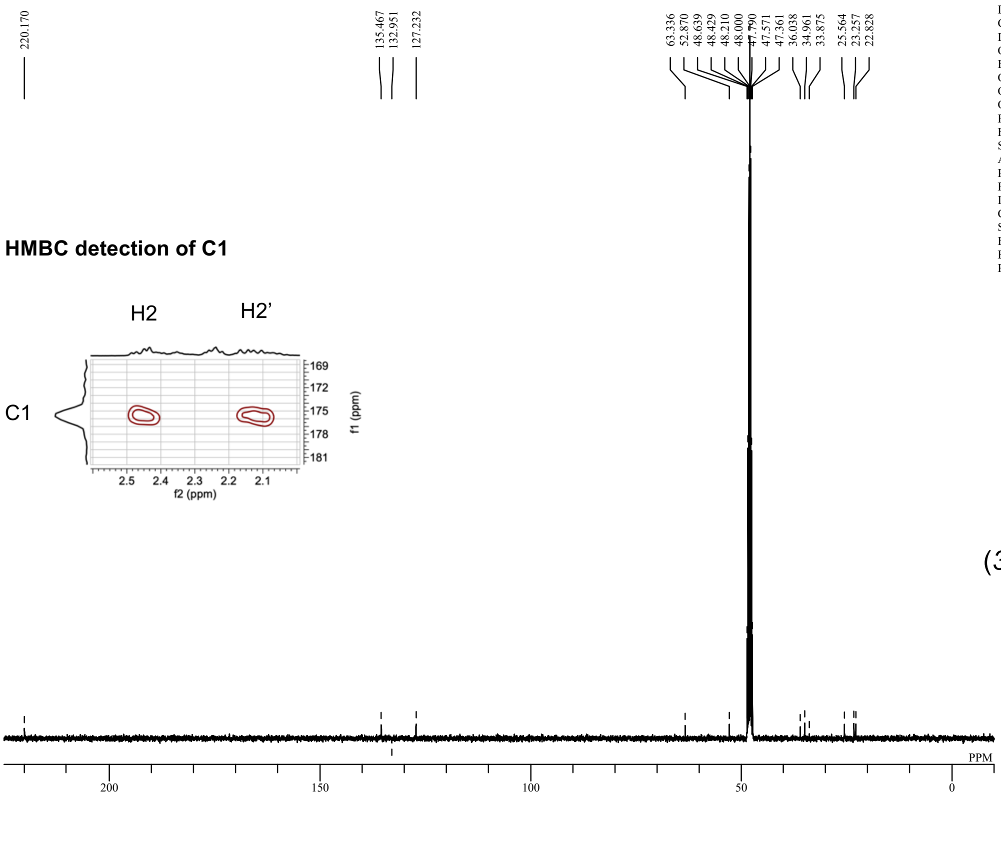


| | |
|---|---|
| DFILE | 3R7S11S-11OH-JA_Carbon-1-1.a |
| COMNT | single pulse decoupled gated NOE |
| DATIM | 14-07-2025 20:51:59 |
| OBNUC | 13C |
| EXMOD | carbon.jxp |
| OBFRQ | 100.53 MHz |
| OBSET | 5.35 KHz |
| OBFIN | 5.86 Hz |
| POINT | 26224 |
| FREQU | 25125.63 Hz |
| SCANS | 481 |
| ACQTM | 1.0433 sec |
| PD | 2.0000 sec |
| PW1 | 3.50 usec |
| IRNUC | 1H |
| CTEMP | 21.5 c |
| SLVNT | CD3OD |
| EXREF | 48.00 ppm |
| BF | 1.00 Hz |
| RGAIN | 50 |

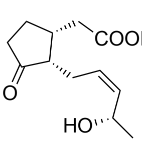


(*3R,7S,11S*)-11-OH-JA

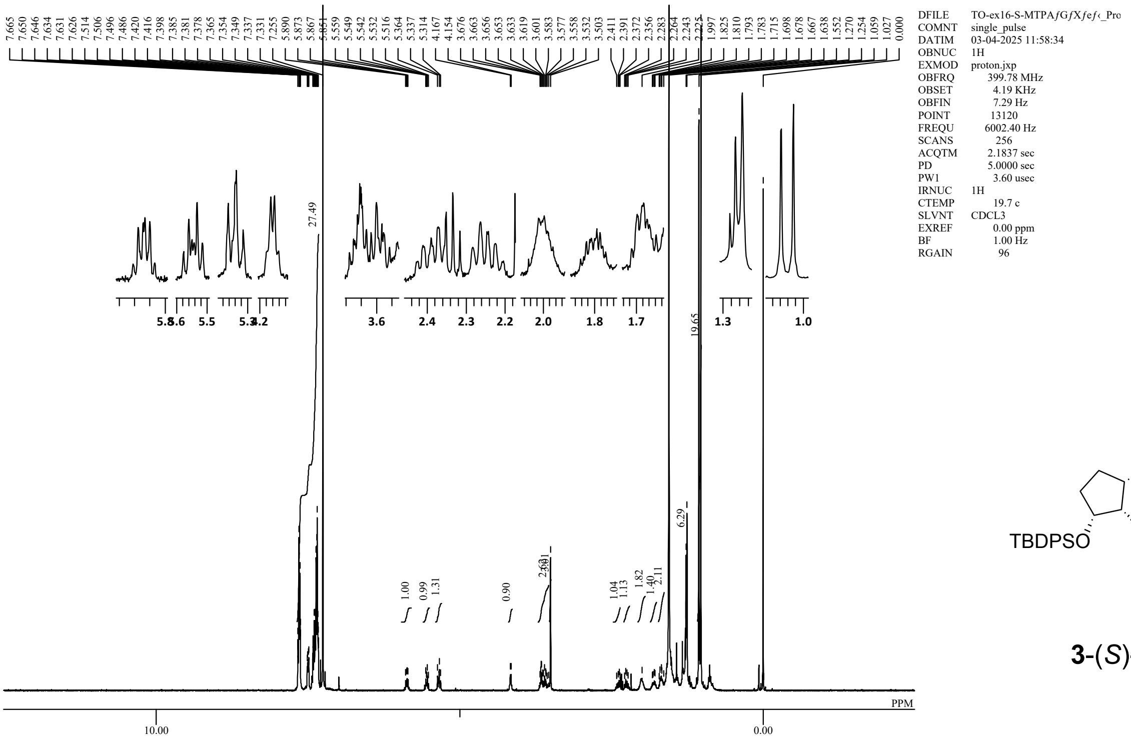


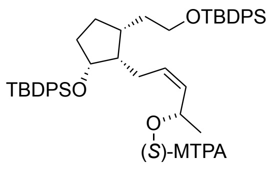


**3**-(*S*)-MTPA

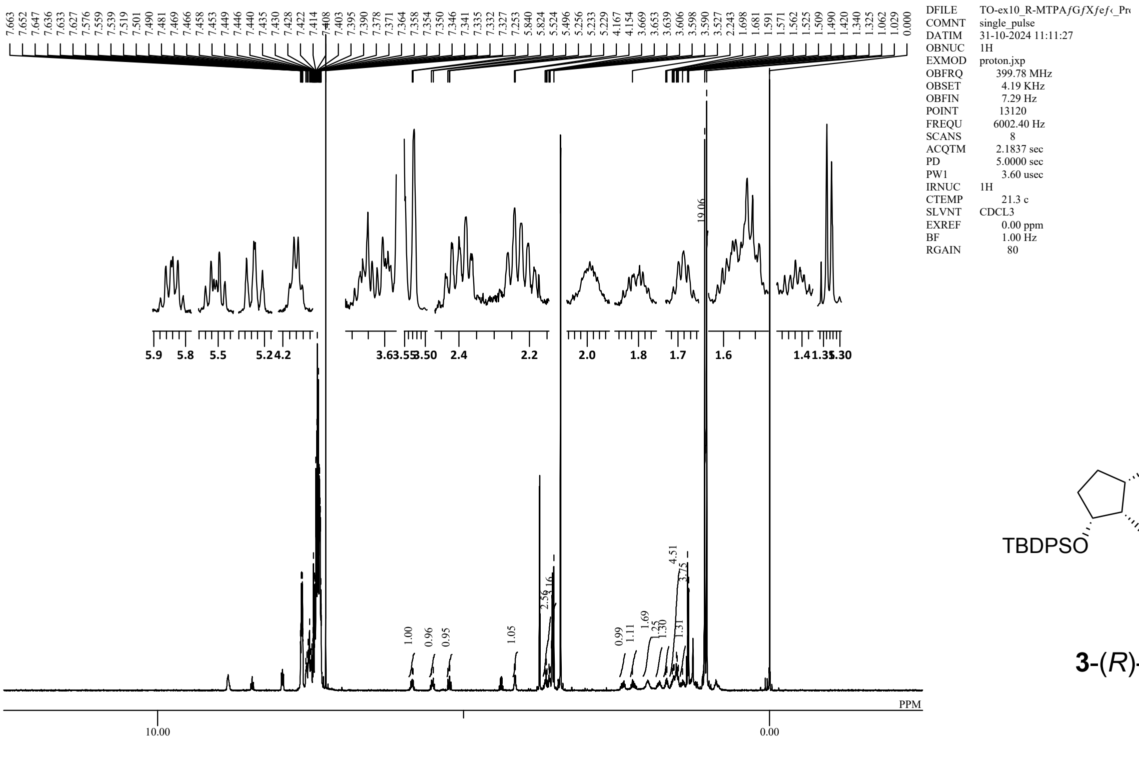


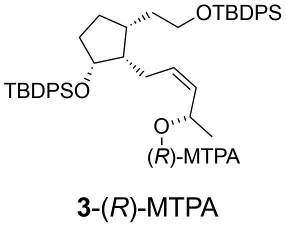


**3**-(*R*)-MTPA
